# ARACRA: Automated RNA-seq Analysis for Chemical Risk Assessment

**DOI:** 10.64898/2026.04.07.716912

**Authors:** Shubh Sharma, Saurav Kumar, Judit Biosca-Brull, Deepika Deepika, Vikas Kumar

## Abstract

Transcriptomic analysis is considered a powerful approach for biomarker discovery, however still exploring large scale omics dataset to extract meaningful biological insights remains a challenge for biologists. To address this gap, we present ARACRA a fully automated RNA-seq analysis pipeline including entire transcriptomics workflow from raw FASTQ files to the transcriptomics Point of Departure (tPoD) with human-in-the-loop review process. Overall, the analysis is performed in two phases: Phase 1 carries out the acquisition of raw reads, pre-alignment quality control, alignment to reference genome and quantification of gene expression. Whereas, Phase 2 performs statistical analysis including Differential Gene Expression analysis and Dose-Response modelling. Two phases are separated by an extensive quality control step which allows the user to visually inspect the quality of data processed and helps in filtering noise and outlier samples. ARACRA facilitates end-to-end analysis of RNA-Seq data through an interactive web-based application developed on nextflow and streamlit for minimizing computational complexities while ensuring correct downstream processing.

**Availability and implementation:** ARACRA is freely available online at the GitHub with MIT License and stream lit-based web application: <u>ARACRA</u>. Researchers can use the demo data or even upload their own data to do the analysis.

Fig 1:
Overall Architecture of ARACRA

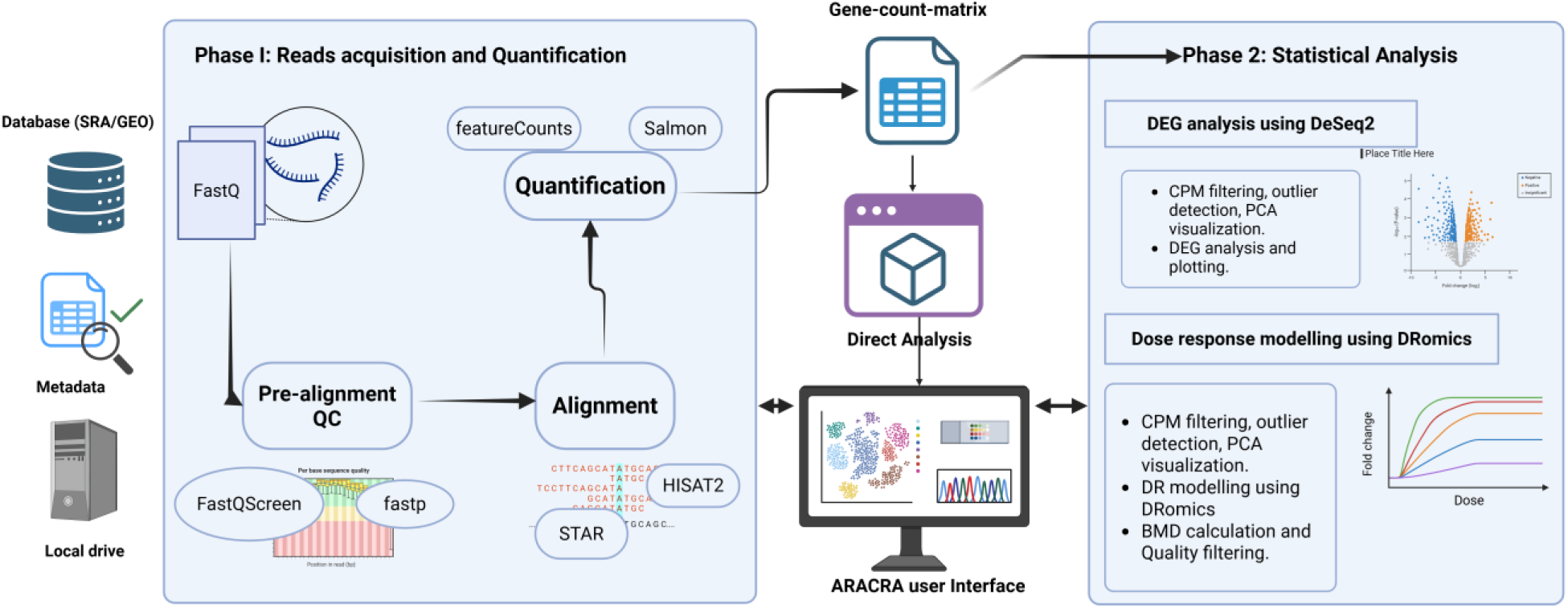

## 1. Introduction

Transcriptomics, comprehensive study of total RNA (coding and non-coding) within a cell or tissue, has become a powerful approach to characterize gene regulation and associated changes to external stressors and conditions [1] [2]. Several strategies are available for measuring transcriptome changes, including microarrays, whole RNA sequencing, and targeted RNA sequencing, which offer higher sensitivity and resolution [3]. In toxicology and risk assessment, RNA-seq in particular has emerged as a new approach methodology (NAM) for biomarker discovery, capturing molecular signals that can be associated with perturbed biological pathways relevant to adverse health outcomes [ [4]. Analyzing the changes in gene expression upon exposure can support multiple applications: identification of shared transcriptional signatures and pathway perturbation patterns can support chemical grouping and read-across, constructing dose-response curves from expression profiles at different exposure concentrations which can then be used to derive Point of Departure (PoD), Benchmark Dose (BMD) contributing in quantitative risk assessment [5] [6].

Numerous bioinformatics platforms and computational pipelines have been developed to facilitate the processing and analysis of HTTr data. Examples include Zavolan-Lab Automated RNA-Seq Pipeline (ZARP) and metaTP, both of which integrate different bioinformatics tools for RNA-seq data analysis [7] [8]. For upstream read processing, nf-core/rnaseq [9] provides a community-curated Nextflow pipeline covering quality control (QC), alignment, and gene quantification with comprehensive Quality Control reporting. Galaxy offers an accessible web-based platform for general-purpose RNA-seq analysis [10]. For dose-response modeling speciifically, BMDExpress 2 [11], maintained by the National Toxicology Program, supports BMD analysis across multiple microarray and sequencing platforms with GO and Reactome pathway classification. BMDx (Serra et al., 2020) and its successor BMDx2 [12] provide R-Shiny interfaces for BMD computation with AOP-based mechanistic annotation. DRomics [13] offers a specialized R framework for omics-scale dose-response analysis with built-in model selection and BMD calculation. In this line, Verheijen et al., 2022 developed the Regulatory Omics Data Analysis Framework (R-ODAF), a framework specifically designed for omics data analysis in regulatory applications. R-ODAF provides an automated pipeline for regulatory-grade transcriptomic analysis including differential expression and robust gene filtering [14].

These tools collectively cover the full analytical steps from raw reads to biological interpretation. However, in practice, a complete toxicogenomics dose-response analysis from FASTQ files to transcriptomic points of departure (tPODs) requires sequential use of multiple independent tools: a read-processing pipeline for QC and quantification, a statistical environment for differential expression, a dose-response modeling tool for BMD estimation, and pathway analysis software for tPOD derivation. Each transition between tools introduces manual format conversions, parameter re-specification, and potential for inconsistency. Notably, upstream pipelines such as nf-core/rnaseq produce count matrices but do not perform differential expression or dose-response modeling; conversely, BMD tools such as BMDExpress and BMDx accept pre-processed expression matrices but do not handle read-level QC or alignment. No single existing tool spans the complete workflow from raw sequencing reads through to regulatory-quality pathway-level tPODs within one reproducible, automated framework. Furthermore, the growing use of targeted TempO-Seq platforms in high-throughput toxicogenomics introduces additional analytical requirements like probe-based alignment, multi-probe gene aggregation, and platform-specific QC that are not natively supported by general-purpose RNA-seq pipelines. To address this, we developed ARACRA (Automated RNA-seq Analysis for Chemical Risk Assessment), a Nextflow DSL2 pipeline with an interactive Streamlit web interface that integrates the complete toxicogenomics workflow into a single, reproducible framework. ARACRA does not provide any new algorithm for RNA-seq data analysis, rather, it orchestrates well-established bioinformatics tools to facilitate end-to-end analysis for non-coding experts, who are interested in overall biology of chemical effects. ARACRA provides: (i) SRA data retrieval through alignment, QC, and quantification for both RNA-seq and TempO-Seq platforms; (ii) interactive multi-layer quality control with human-in-the-loop outlier review; (iii) integrated DRomics dose-response modeling with quality filters; (iv) pathway-level tPOD computation across GO, KEGG, and MSigDB gene set collections with multiple tPOD derivation methods; and (v) a web interface enabling interactive parameter tuning and result visualization without command-line expertise. For validation, ARACRA was applied to a publicly available TempO-Seq dataset from Beal et al. (2024) examining BPA and 11 alternatives exposure in MCF-7 cells [15].

## 2. Implementation

### 2.1 Architecture overview

ARACRA is implemented in Nextflow DSL2 workflow management system with R scripts for statistical analysis and a Python-based Streamlit web interface for user interaction. The pipeline is divided into two phases separated by an interactive quality control gate. Phase 1 includes read acquisition, pre- alignment quality control (QC), alignment, post-alignment QC, gene quantification, and outlier detection. A detailed QC report is generated at the end of Phase 1 for users to visually inspect the quality of data and remove poor quality samples from downstream processing. Phase 2 includes differential expression of genes analysis and dose-response (DR) modelling for biological interpretation of the RNA-seq data. This separation ensures interactive data visualization and inspection to ensure high quality data is processed, excluding any technical artifacts under the user’s supervision.

ARACRA additionally supports a Direct Analysis mode in which users can provide a pre-existing count matrix and metadata file, bypassing the preprocessing phase entirely. In this mode, a PCA-based QC check runs first, followed by the same interactive review gate before analysis proceeds. It also allows the user to analyze Tempo-Seq data (targeted RNA-seq data) which uses specific probe set to identify transcriptomic perturbations following chemical exposure. Fig 2. represents the overall architecture of ARACRA.

**Fig 2:**
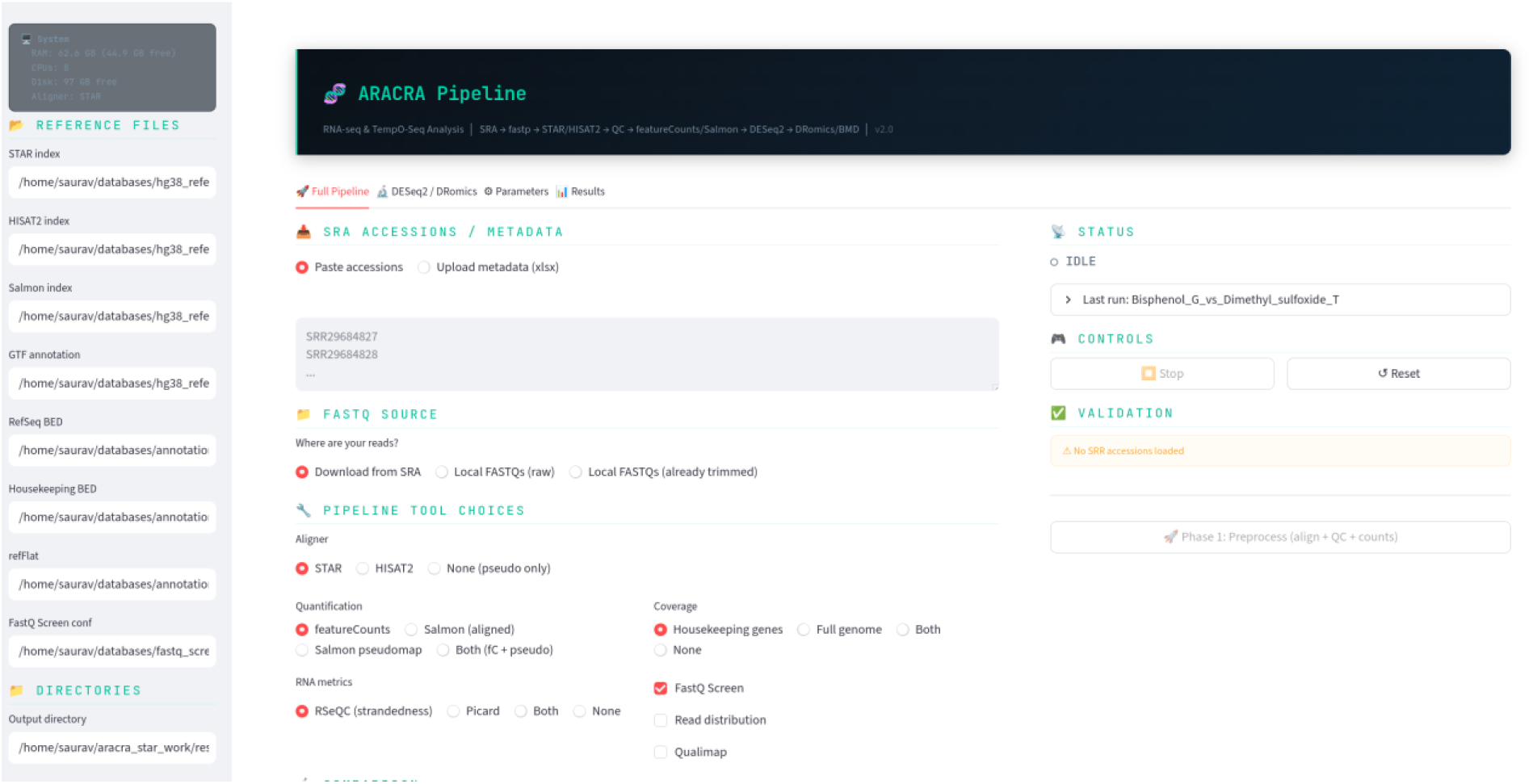
**User Interface of ARACRA**

### 2.2. Phase 1: Reads acquisition and Quantification

#### 2.2.1 Data Retrieval and Pre-alignment Quality Control

The pipeline provides two options for retrieving data. Data can be downloaded directly from Sequence Read Archive (SRA) database using SRA accession number or retrieving the in-house data from local directory in FASTQ format.

Accession numbers provided directly retrieve the raw data from the SRA database and is converted into FASTQ format using the fasterq-dump tool provided by SRA toolkit. Whereas local data, directly acquired from the directory through an assigned ID and proceeds for quality check and control. Adapter trimming and quality filtering are performed by fastp [16], producing per-sample QC reports. It utilizes an adaptable configuration file with default parameters including a 4-base pair sliding window and a minimum Phred quality score threshold of Q30. For each sample, fastp generates standardized JSON and HTML reports detailing critical metrics:

- Per-base quality distribution before and after filtering
- Read length distribution and duplication rates
- Adapter content and GC content bias
- Pass/fail read count ratios.

Contamination screening is optionally performed using FastQ Screen [17] against reference genomes for common contaminants. FastQ Screen maps a subset of reads against comprehensive reference databases and identify multi-genome contamination and detect residual ribosomal RNA (rRNA). Required FastQ Screen indices for different target and contaminant genomes along with the corresponding configuration files—will be downloaded and built during the initial computational environment setup using the get_genomes protocol.

#### 2.2.2 Alignment

ARACRA provides two distinct splice-aware RNA-Seq aligners- HISAT2 (Hierarchical Indexing for Spliced Alignment of Transcripts) and STAR (Spliced Transcripts Alignment to a Reference) [18] [19]. Two aligners are provided for optimal use of computational resources where HISAT2 can be configured with system with low resources (<32 GB RAM) while STAR requires high computational resources (>32 GB RAM).

During the initial environment setup, the bash script handles the preparation of reference indices. For HISAT2, the script downloads a pre-built, transcriptome-aware human index (grch38_tran.tar.gz) directly, which saves time building the index from scratch. For STAR, the setup file uses standard human genome FASTA and corresponding GENCODE GTF file to build the index once which can be referred to every time. However, for targeted TempO-Seq data, the pipeline employs a fundamentally different, dynamic indexing strategy. Because TempO-Seq relies on a highly restricted set of target-specific, ∼50-base pair probes rather than the full transcriptome, mapping against the standard human genome would be computationally inefficient and prone to off-target mapping.

Instead, the pipeline parses the user-provided manifest file during execution to generate a custom, miniature FASTA file containing exclusively the exact probe sequences. Using STAR, the localized index is then built directly from this customized FASTA. To prevent memory segmentation faults caused by this drastically reduced file size, the pipeline automatically calculates and adjusts the index scaling parameter (--genomeSAindexNbases) prior to generation. Once this custom index is built, the TempO-Seq reads are mapped with strictly tuned STAR parameters explicitly disabling splicing (--alignIntronMax 1) and severely restricting mismatch tolerances to maximize the accuracy of probe-level quantification.

#### 2.2.3 Quantification

Standard alignment-based quantification is executed using featureCounts from the Subread package and Salmon [20] [21]. This utility processes the filtered BAM files generated by STAR or HISAT2 alongside the reference GTF annotation to accurately assign reads to known genomic features. featureCounts dynamically adapts to the empirically inferred library strandedness and read layout to ensure high-fidelity gene-level aggregation. Salmon provides higher accuracy in terms of identification and handling of novel transcripts and isoforms. It uses the exact library strandness from STAR and performs quasi-mapping against the reference transcriptome. A rapid pseudo-alignment option is also available in which salmon bypasses the resource-heavy STAR/HISAT2 genomic alignments entirely, deploying Salmon’s quasi-mapping algorithm directly on the trimmed FASTQ files with automatic library-type detection (-l A). For both Salmon pathways, the resulting transcript-level expression estimates are seamlessly aggregated into a robust gene-level count matrix using an automated transcript-to-gene dictionary (tx2gene).

For targeted TempO-Seq data, the pipeline employs a specialized quantification workflow. Probe-level alignment frequencies are extracted directly from the STAR-generated mapping files using samtools idxstats. A custom Python aggregation script then processes these frequencies, utilizing the provided BioSpyder manifest file to map and consolidate probe counts directly back to their respective target genes.

#### 2.2.4 Post Alignment Quality Control

Extensive Quality control is performed after alignment of reads to ensure high-fidelity data is processed downstream. It integrates RSeQC computational suite [22] to perform structural profiling of the aligned libraries which uses several modules such as inferexperiment.py, read_distribution.py and geneBody_coverage.py to validate library strandness, categorization of mapped reads across distinct genomic features and evaluating potential 5’-> 3’ transcript degradation respectively. Apart from RSeQC, Picard and Qualimap [23] is also configured in the pipeline to provide visual diagnostics of the samples. A detailed summary report is generated in a single, interactive HTML dashboard using MultiQC [24] to provide the users with a holistic overview of the samples. Samples with more than 70% of reads having a Phred quality score below Q30 is flagged as poor-quality samples and is removed from further analysis. Samples with less than 70% of reads uniquely mapped to the reference genome is flagged as outliers and left to the user for further inspection before deciding their inclusion or removal from downstream analysis. These thresholds are based on the R-ODAF pipeline developed by Verheijen et al., 2022.

#### 2.2.5 Exploratory data analysis and layer II outlier detection

After gene quantification, data is normalized using variance-stabilizing transformation (VST) and if batch variable exists, batch correction is done using ComBat-seq [25]. Principal Component Analysis is done on batch corrected counts and visualized in two-dimensional space. Within each experimental group, samples with Euclidean distances exceeding the group mean plus two standard deviations in PC1-PC2 space are flagged as outlier. JSON reports are generated and visualized in graphical user interface for human in loop intervention to include or exclude flagged outliers.

### 2.3. Phase 2: Statistical Analysis

Counts per Million (CPM)-based gene filtering is done to filter out noise genes (genes must exceed 1 CPM in at least 75% of samples within at least one experimental group by default). DESeq2 [26] is used for differential expression analysis on raw integer counts using batch (if available) and treatment as covariate to ensure no over correction is implied. For dose-response studies with multiple treatment concentrations, per-dose contrasts against the control group are computed in addition to the overall treated-vs-control comparison. Gene symbols are resolved using the org.Hs.eg.db annotation database, with fallback to symbols embedded in Salmon/GENCODE transcript identifiers when available. Results include volcano plots, PCA plots, and CSV files of all results, significant DEGs, and per-dose comparisons. Dose-response analysis is performed using DRomics R package to analyze the response of genes across different doses. Table 1 provides a list of bioinformatics tools used in ARACRA Genes are tested for response using one of the three tests: quadratic, linear or ANOVA with FDR threshold of 0.05. Dose response curves are fit using drcfit () function testing 5 different models () and selects the best-fitting model via the Akaike Information Criterion corrected for small samples (AICc). Fitting is parallelized across available CPU cores using SNOW clusters to manage computation time, particularly for datasets with hundreds of responsive genes. Benchmark Dose (BMD) values are derived from each fitted curve using bmdcalc () function, which reports both the zSD metric (the dose producing a response equal to one standard deviation of the control distribution) and the xfold metric (the dose producing a specified percent change from baseline). It also provides BMD calculations using bootstrap confidence intervals (bmdboot function) with a configurable number of iterations [13]. For each gene, ARACRA computes three metrics: variance explained by the fitted curve relative to total variance, signal to noise ratio (maximum predicted response change divided by residual standard deviation), and effect size (response range as a fraction of baseline expression). To ensure reliability of BMD estimates, several filters are available which can be configured independently. The max-dose filter removes genes whose BMD exceeds the highest experimentally tested dose, as such estimates require extrapolation beyond the observed data range. The extrapolation flag identifies genes whose BMD falls below the lowest tested dose divided by a configurable factor (default: 10), indicating that the estimated potency relies on downward extrapolation; these genes are flagged but retained, with warnings propagated to pathway-level results. The BMDU/BMDL ratio filter removes genes for which the bootstrap confidence interval is excessively wide (ratio exceeding 40 for NTP), indicating poor precision in the BMD estimate. Finally, the fold-change pre-filter removes genes whose maximum expression difference between any treated group and controls falls below a user-defined threshold in the transformed expression space, ensuring that only biologically meaningful responses contribute to downstream tPOD estimation. This filter is applied after model fitting to avoid disrupting the internal consistency of the DRomics itemselect object.

**Table 1:**
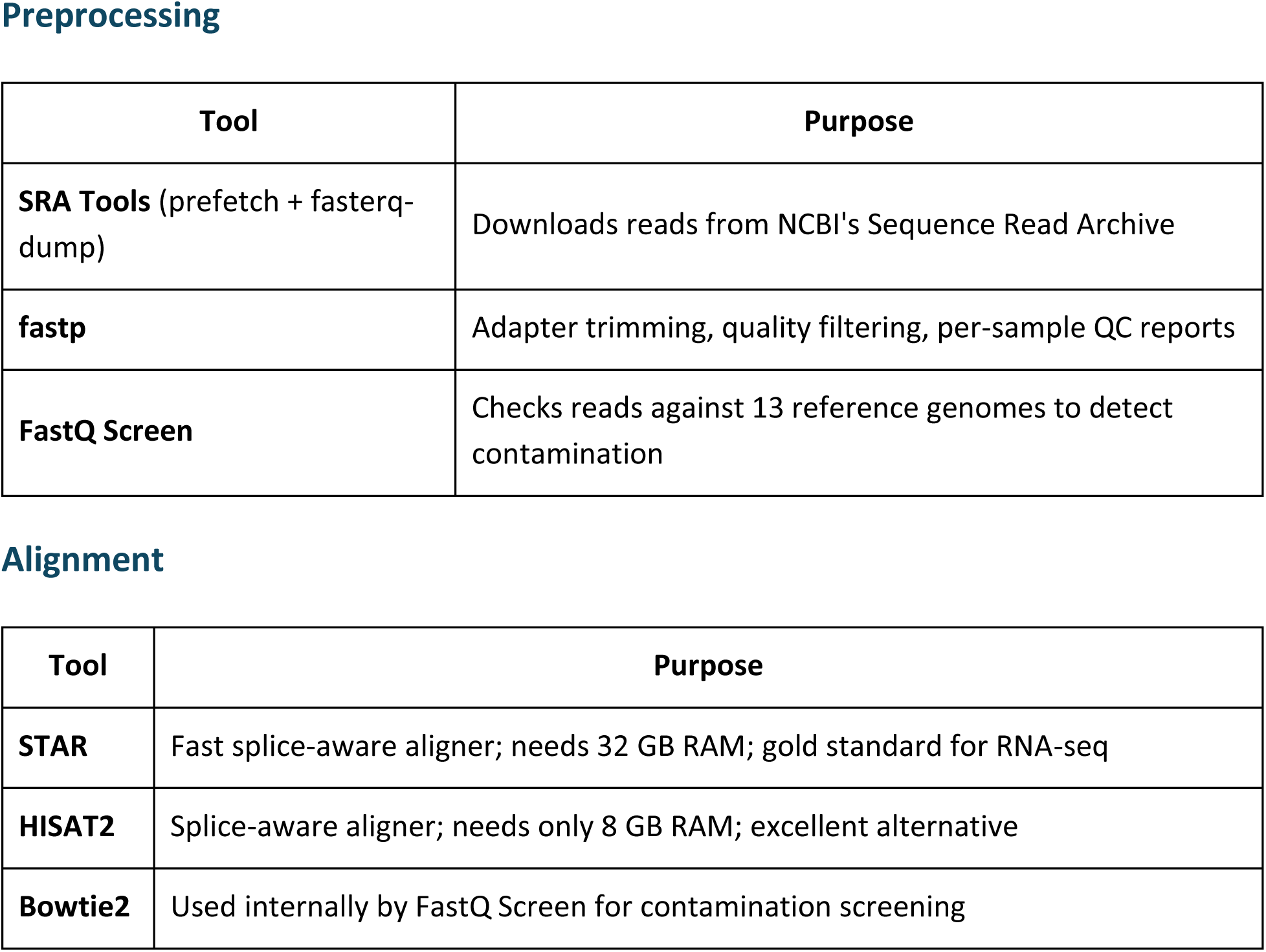
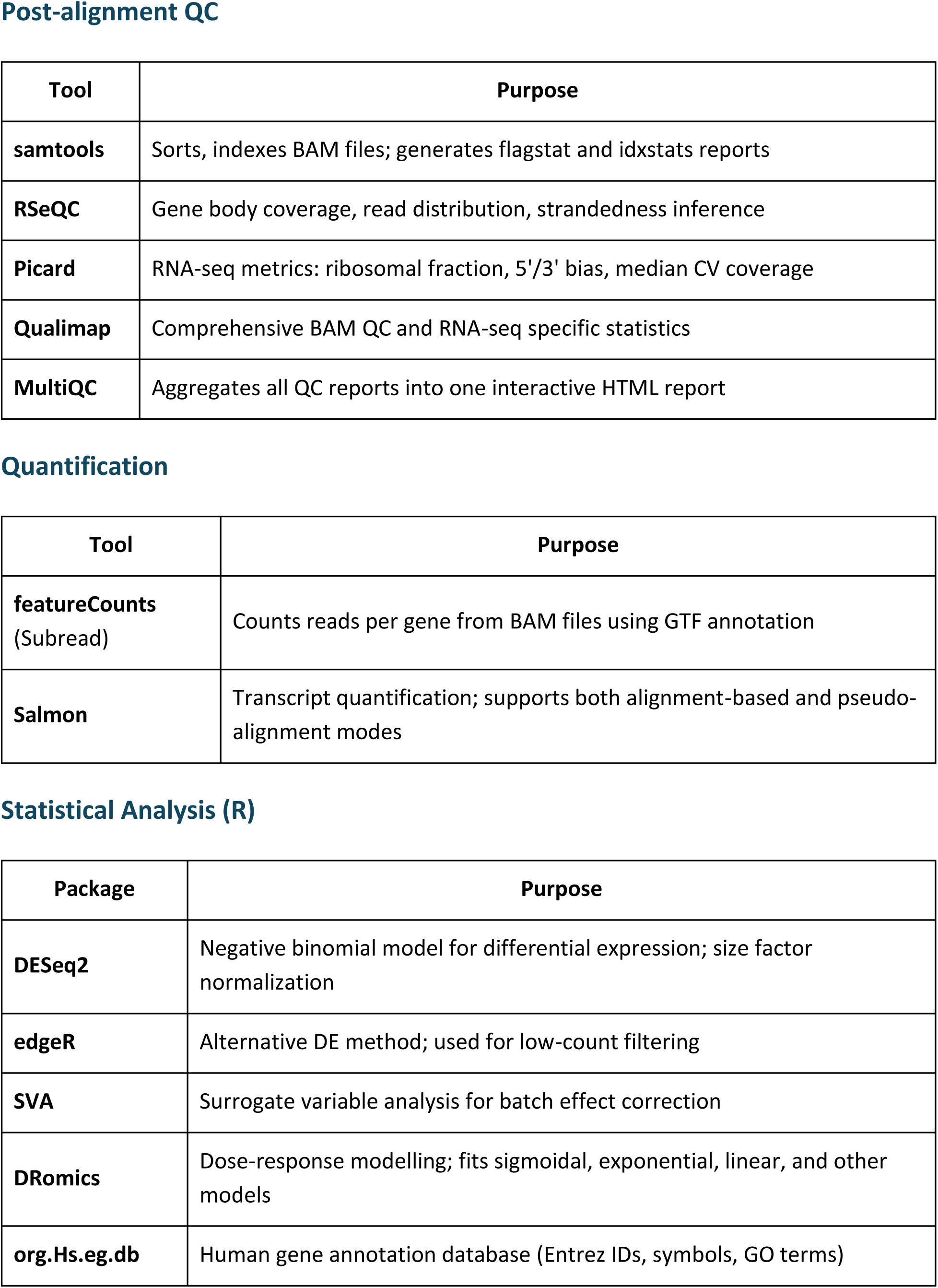
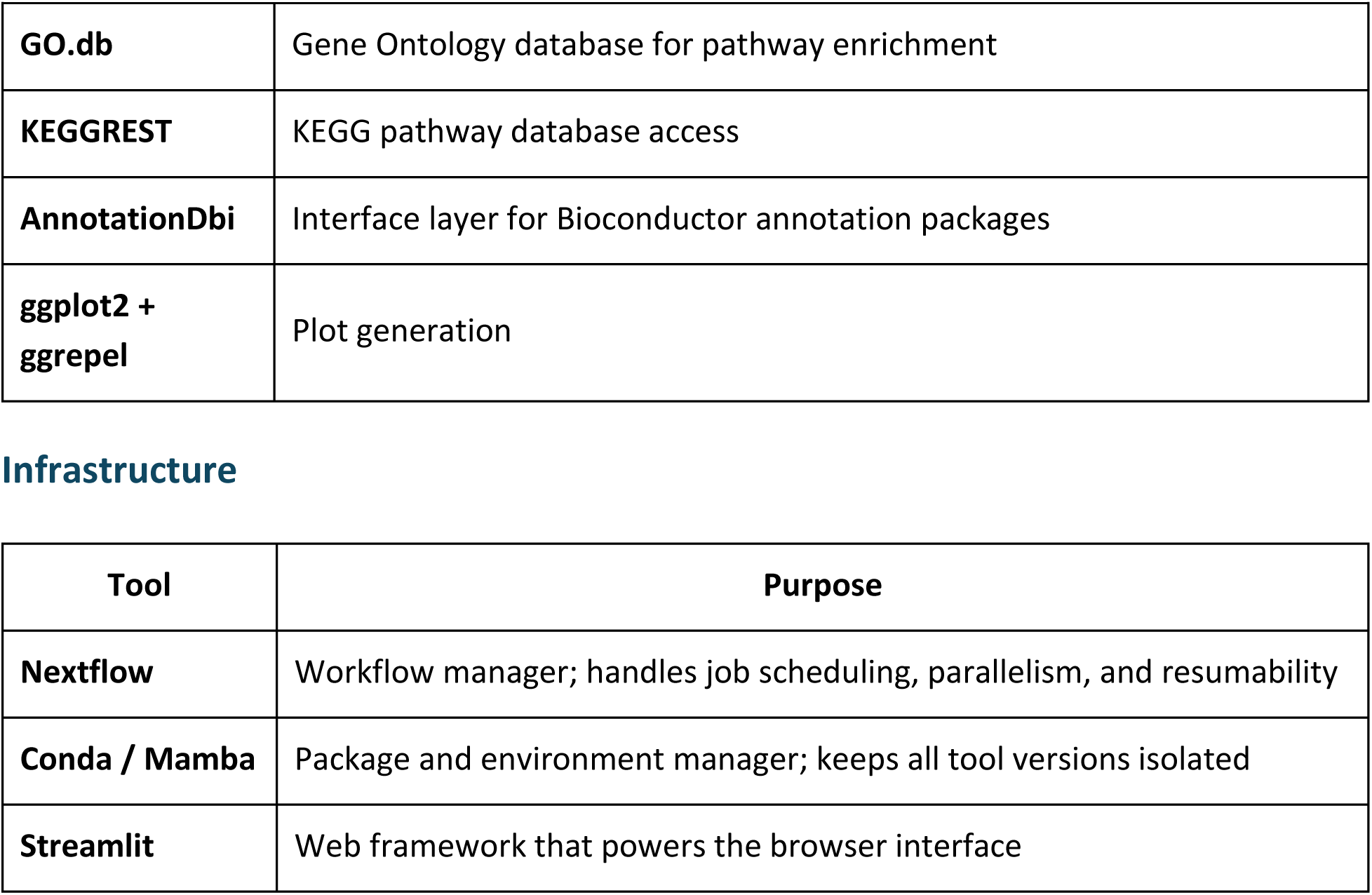
Various bioinformatics tools used in ARACRA along with their purpose.

ARACRA computes pathway-level tPODs following the NTP 2018 gene set-based approach [27]. Quality-filtered BMD genes are mapped to four pathway collections GO Biological Process, KEGG (via org.Hs.eg.db) [28], MSigDB Hallmark, and MSigDB C2 (via msigdbr) [29] with pathway sizes derived from the full genome annotation to ensure accurate coverage estimates. For each pathway, the median BMD of its member genes serves as the sensitivity estimate; pathways qualify if they contain at least 3 BMD genes, cover at least 5% of total membership, and (for GO terms) fall between 10 and 2,000 total genes to exclude both unstable small terms and overly generic ones. The tPOD is defined as the median BMD of the most sensitive qualifying pathway, with per-collection breakdowns reported separately. As a complement, following distribution-based tPOD methods are computed directly from the ranked gene-level BMD distribution: the 25th ranked gene BMD [30], the 5th and 10th percentile BMDs [31], the first mode via ECDF maximum curvature [32], enabling direct comparison between pathway-based and distribution-based frameworks within a single analysis.

### 2.4 Resource management, Initial setup and reproducibility

The setup.sh installation script initiates the setup of ARACRA and subsequent dependencies. It auto-detects system resources (CPU cores, available RAM, disk space) and configures the Nextflow process labels accordingly. Lightweight processes such as fastp trimming are parallelized across available cores, while memory-intensive processes such as STAR alignment are limited to two concurrent instances to prevent out-of-memory failures. On systems with less than 32 GB RAM, HISAT2 is automatically recommended over STAR, and the generated configuration reflects this recommendation.

### 2.5 Web Interface

A streamlit-based web interface is provided for interactive usage and visualization (Fig 2). The interface is organized into four tabs: Full Pipeline (two-phase workflow from raw reads), Direct Analysis (count matrix input), Parameters (configurable thresholds for DESeq2 and DRomics), and Results (interactive display of analysis outputs). The sidebar provides reference file paths and execution profiles installed during the setup phase. Detailed description of User interface is provided in the supplementary material S2_User_manual.docx. The interface requires no installation beyond the conda environment created by setup.sh.

### 2.6 Availability

ARACRA is available freely at the Github: ARACRA . A detailed user manual for first time setup and usage is also available with a need for broader community interaction and enrichment.

## 3. Validation

ARACRA was used to analyse public dataset from Beal et al., 2024 consisting of HTTr data of Bisphenol A and eleven data-poor alternatives. Reads of 286 samples were retrieved and processed through the application to generate a quantified gene-expression matrix, which was used later for DEG analysis and BMD modelling. For BPA, ARACRA identified 1,181 dose-responsive genes from 11,889 analyzed (9.9%), fitted dose-response models to 892, and retained 282 BMDs after applying NTP 2018 and EFSA quality filters. The BMDU/BMDL ratio filter (threshold < 40) was the dominant quality filter, removing 62.5% of BMDs, consistent with the inherent imprecision of bootstrap confidence intervals for omics-scale dose-response data. For 2,4’-BPA, the pipeline identified 1,459 responsive genes with 1,127 fitted models and 404 quality-filtered BMDs, demonstrating higher transcriptomic activity than BPA. Detailed analysis is available in the supplementary S1.docx and table 2 reports range of tPoD from different methods for the chemicals. The order of potency of the chemicals was similar to what Beal et al., 2024 reported with BPA and 2,4’ BPA having most number of differentially expressed genes and genes which were responding to changes in doses. Whereas, Bisphenol BP and Tetramethylol Bisphenol A were inactive.

**Table 2:**
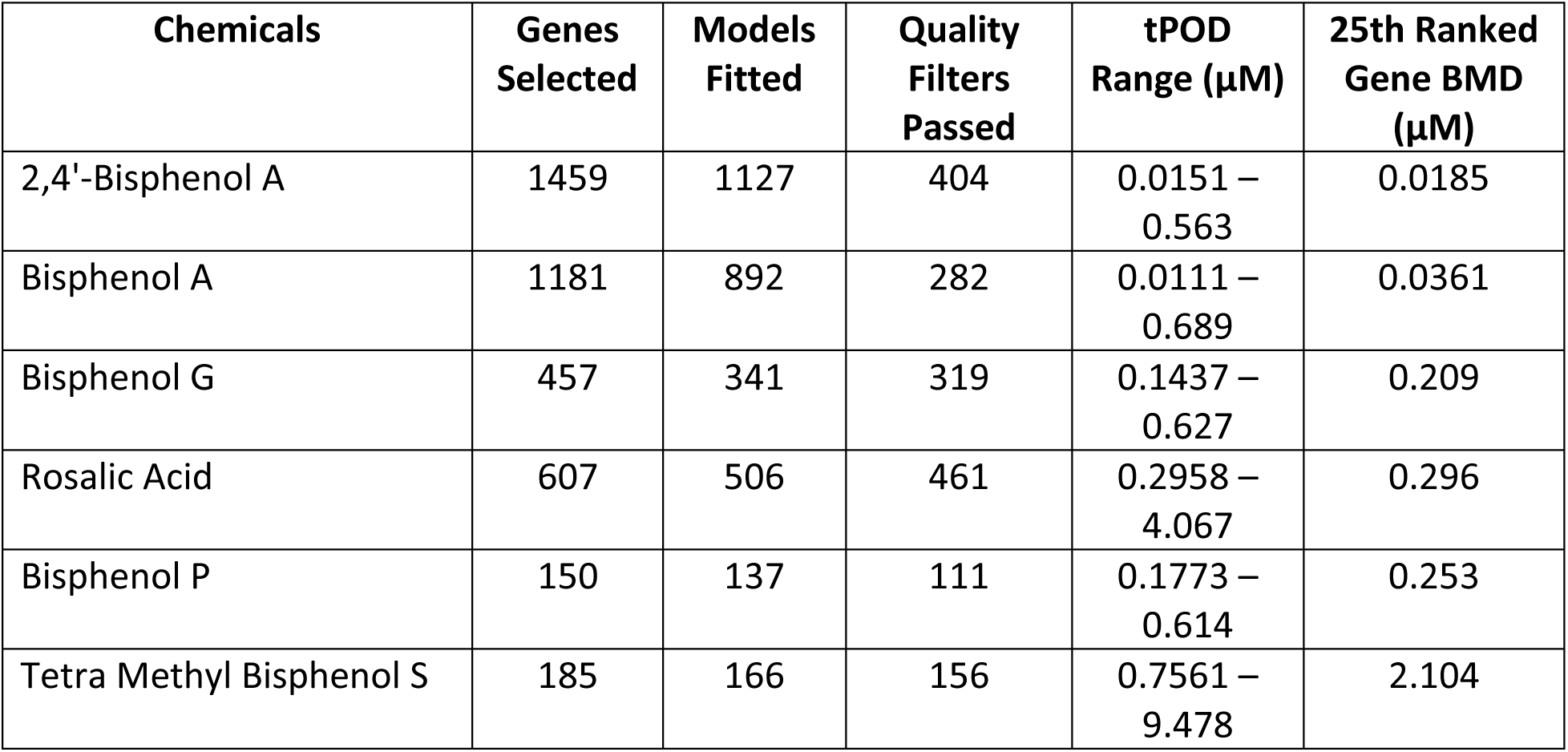
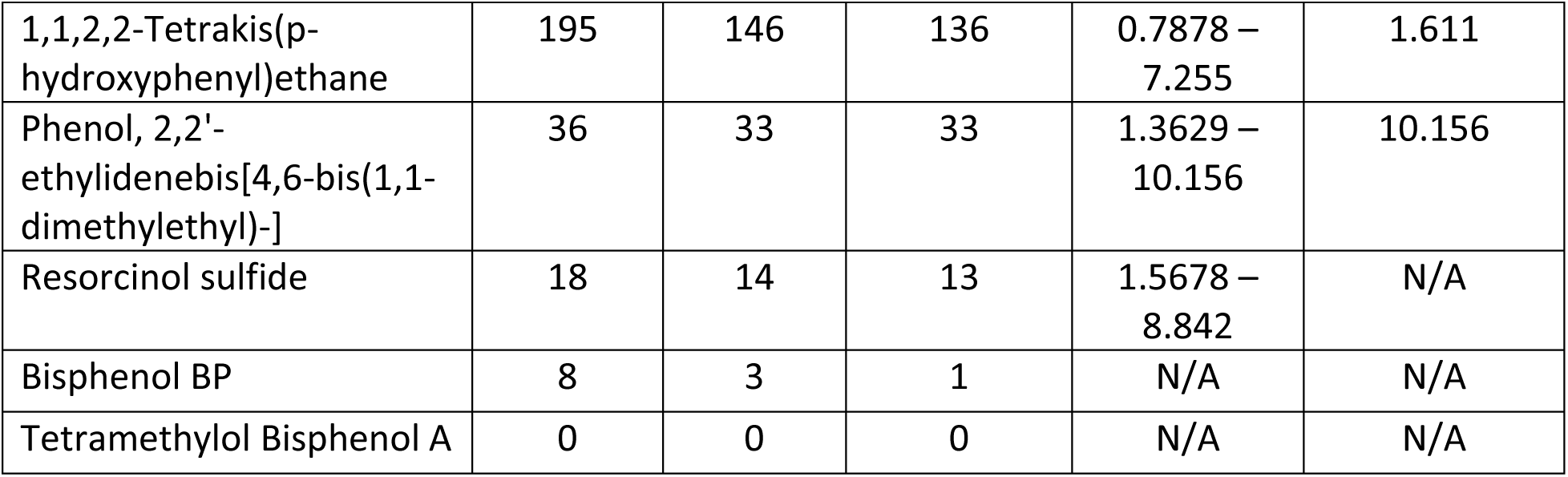
Range of tPoDs derived from ARACRA for BPA and alternatives.

| **Chemicals** | **Genes Selected** | **Models Fitted** | **Quality Filters Passed** | **tPOD Range (μM)** | **25th Ranked Gene BMD (μM)** |
| --- | --- | --- | --- | --- | --- |
| 2,4'-Bisphenol A | 1459 | 1127 | 404 | 0.0151 – 0.563 | 0.0185 |
| Bisphenol A | 1181 | 892 | 282 | 0.0111 – 0.689 | 0.0361 |
| Bisphenol G | 457 | 341 | 319 | 0.1437 – 0.627 | 0.209 |
| Rosalic Acid | 607 | 506 | 461 | 0.2958 – 4.067 | 0.296 |
| Bisphenol P | 150 | 137 | 111 | 0.1773 – 0.614 | 0.253 |
| Tetra Methyl Bisphenol S | 185 | 166 | 156 | 0.7561 – 9.478 | 2.104 |
| 1,1,2,2-Tetrakis(p-hydroxyphenyl)ethane | 195 | 146 | 136 | 0.7878 – 7.255 | 1.611 |
| Phenol, 2,2'-ethylidenebis[4,6-bis(1,1-dimethylethyl)-] | 36 | 33 | 33 | 1.3629 – 10.156 | 10.156 |
| Resorcinol sulfide | 18 | 14 | 13 | 1.5678 – 8.842 | N/A |
| Bisphenol BP | 8 | 3 | 1 | N/A | N/A |
| Tetramethylol Bisphenol A | 0 | 0 | 0 | N/A | N/A |

## 4. Discussion

Although robust tools exist for individual analytical tasks, end-to-end analysis from raw sequencing reads to transcriptomic point of departure (tPOD) derivation still typically requires manual transfer of data, parameters, and decisions across multiple software environments. In regulatory and applied toxicogenomics settings, this fragmentation can reduce reproducibility, track flow of data, and may require consistent human engagement for processing and outputs. ARACRA does not introduce a new statistical method; rather, its contribution lies in the structured orchestration of established tools within a single reproducible workflow, coupled to an interactive review layer that preserves expert oversight at critical decision points. ARACRA is available freely on GitHub. InSilicoVida-Research-Lab/ARACRA for processing raw fastq files and deriving tPODs for the chemicals. Considering that one of the major issues in RNA-seq data analysis is reproducibility and false positives [32], every parameter used, exclusion and inclusion decision of the samples and intermediate result is preserved in structured JSON outputs for reproducibility and efficient preprocessing protocol established on the foundations of regulatory pipelines with user review. ARACRA derives its operation using well established bioinformatics tools and eliminates the need for manual command-line wrapping of the operations. It is based on general purpose RNA-seq analysis tools such as nf-core or Galaxy but also is tailored specifically for chemical exposure data by including dose-response analysis through DRomics.

In the analysis of dataset from Beal et al., 2024, similar patterns of results were obtained. In our analysis also we found that 2,4’BPA produced maximum effect at the concentration of 10 micromolar and also was transcriptionally more active than BPA (25th ranked gene tPOD lower than that of BPA). The tPoD range derived in ARACRA from various methods had comparable range reported by Beal *et* al, who used R-ODAF pipeline along with BMD express for calculating tPOD. ARACRA’s 25th ranked gene BMD point estimates fell within the BMCL–BMCU confidence intervals reported by Beal et al. for 8 of 10 chemicals with derivable tPODs, demonstrating numerical as well as ordinal concordance. The primary difference was observed for RS, for which ARACRA did not derive a Rank25 tPOD due to fewer than 25 genes passing quality filters, likely reflecting the stricter combined effect of the max-dose and BMDU/BMDL filters applied in ARACRA. Several genes were found to be differentially expressed which were involved in endocrine disruption from previous studies such as PGR, GREB1. GREB1 acts as a co-activator for ERα, stabilizing interactions between ER and other co-factors, and modulating the expression of ER target genes, and therefore alteration in expression of GREB1 can be a potential cause of endocrine disruption resulting in increased cell proliferation [33]. PGR gene encodes for a nuclear receptor for progesterone, a key steroid hormone involved in reproductive tissue differentiation, ovulation, and pregnancy. Both GREB1 and PGR are induced by estrogen via ERα and their disruption by EDCs can lead to impaired cell cycle regulation and other processes [34]. Similar results were reported by Beall et al., 2024, ranking the chemicals based on the activity and transcriptomics response setting a benchmark for this analysis. However, to validate these results and identify BMD for ERα genes activated at lower doses, further investigations are required.

Within the EU-PARC project where an effort is being made for using transcriptomics data in an harmonized way for transparent data processing, metadata harmonization, standardized reporting, and cross-study comparability, ARACRA not only provides an interactive computational convenience for the users but also aims for standardizing workflow and processing of the transcriptomics data for preserving traceability, comparability, and interpretability.

## 5. Conclusion and Future Work

ARACRA provides an easy-to-use platform which can be configured locally for end-to-end RNA seq analysis and used in the field of chemical risk assessment. It eliminates the need for manual command line actions with an interactive user interface for overall review by the experts, thereby making RNA-seq analysis easier for non-coding researchers. It also provides reproducibility and transparency by storing the results and parameters used for future reference and downstream reporting.

Some limitations exist, which will be improved in further course of time with wider community support. In the current setup file, only human transcriptome is referenced, limiting its applicability to rodent toxicogenomic data. Also, unlike BMDx2, ARACRA does not currently integrate Adverse Outcome Pathway (AOP) mapping. Future development will focus on multi-species support, integration of AOP frameworks for mechanistic anchoring. For statistical analysis, multiple options will be made available with respect to packages being used (for example: edgeR, BMDexpress). Also, WGCNA and GSEA modules will be added for understanding synergetic effect of the genes and their biological interpretation through pathways. As the next step, individual steps of the pipeline will be containerized as MCP (Model Context Protocol) server, which will be available for developing AI-agent (LLM based) assisted transcriptomics analysis where LLM models can trigger automated analysis steps using harmonized metadata (using OECD Omics Reporting Framework). This architecture aims to significantly lower the technical barrier for researchers, abstracting computational complexities with human-in-loop biological interpretation and hypothesis generation.

## Supporting information

csv file

HTML file

## Data Availability

High-Throughput RNA-Seq data was used from GEO database under the accession number GSE271332. All the codes used in this work is available on GitHub: <u>InSilicoVida-Research-Lab/ARACRA</u>

## Supplementary File

Supplementary file S1 and S2. Full tutorial for the application is available in supplementary file S2_User_Manual.docx.

