## Supplement file 1 for "ARACRA: Automated RNA-seq Analysis for Chemical Risk Assessment"

**Supplementary document**

**Validation:**

The dataset having accession number GSE271332 was analyzed using ARACRA using the metadata information S4_metadata.csv. The dataset had Tempo-Seq HTTr data of BPA and 11 alternatives tested at different concentrations in 3 batches as per Beal *et* al., 2024. In total, 286 samples were present, raw data of which was retrieved from SRA database. TempoSeq manifest file provided by BioSpyder Technologies was used to build the local reference index against which the samples were mapped and quantified.

**Overall QC report from MultiQC revealed following information:**

Fastp filtering retained an average of 99.83% of reads per sample across 286 samples, with negligible adapter contamination (maximum 2.27%) and minimal low-quality or too-short read fractions (<0.15% and <1.3%, respectively), confirming high raw sequencing quality. STAR alignment yielded a mean unique mapping rate of 94.6%, with only 6 samples having unique mapping rate less than 70%. These 6 samples were removed from the further analysis. Library sequencing depth ranged from approximately 0.4 to 7.3 million reads per sample (mean 3.6 million; median 3.7 million), with the majority of libraries (250/286) exceeding 2 million reads. FastQ Screen confirmed predominantly human origin of the sequenced reads, with no detectable contamination from common laboratory organisms. Being TempoSeq data, duplication rate was high as it is probe targeted mechanism of sequencing. A detailed Multi-QC report is attached as S3_multiqc.html.

**Further Downstream processing:**

Gene-counts matrix was then used for differential gene expression analysis and dose-response modelling. An additional layer quality control step was performed which help the users to filter noise genes in the dataset. As soon as the user select chemical and its corresponding control in the menu, subset of count tables is created specific to samples for those chemicals by mapping through accession numbers between count matrix and metadata. Batch correction was performed using ComBat Seq method and samples were visualized using PCA analysis. Genes having less than 1 CPM within its treatment group were removed from analysis. Approximately 11500 genes passed the threshold per chemical and were considered for DEG and dose response analysis. Subsequent data were retrieved from raw count matrix and utilized by DeSeq2 package with treatment as design (Chemical v/s control) and batch as covariate for identifying differentially expressed genes. Overall results of DEGs identified are summarized in table 1 below:

| **Chemical Name** | **PCA Outliers Removed** | **Total Genes in Matrix** | **Genes After Low-Count Filtering** | **Upregulated** | **Downregulated** | **Dose Levels Tested** |
| --- | --- | --- | --- | --- | --- | --- |
| 1,1,2,2-Tetrakis(p-hydroxyphenyl)ethane | 1 | 19686 | 12017 | 9 | 2 | 5 |
| 2,2'-Methylenebis(4-methyl-6-tert-butylphenol) | N/A | 19686 | 11370 | 346 | 163 | 5 |
| 2,4'-Bisphenol A | 2 | 19686 | 11884 | **554** | 203 | 5 |
| Bisphenol A | 0 | 19686 | 11886 | 497 | 148 | 5 |
| Bisphenol BP | 0 | 19686 | 12063 | 0 | 0 | 5 |
| Bisphenol G | 1 | 19686 | 11582 | 245 | 42 | 5 |
| Bisphenol P | 0 | 19686 | 11498 | 63 | 15 | 5 |
| Phenol, 2,2'-ethylidenebis[4,6-bis(1,1-dimethylethyl)- | 2 | 19686 | 11888 | 0 | 0 | 5 |
| Resorcinol Sulfide | 1 | 19686 | 11690 | 2 | 0 | 5 |
| Rosalic Acid | 0 | 19686 | 11683 | 87 | 8 | 5 |
| Tetra Methyl Bisphenol S | 1 | 19686 | 9607 | 19 | 2 | 5 |
| Tetramethylol Bisphenol A | 0 | 19686 | 7493 | 3 | 6 | 5 |

Table 1: Summary of DEGs when compared overall exposure of chemicals against corresponding control samples.
