## Supplement file 2 for "ARACRA: Automated RNA-seq Analysis for Chemical Risk Assessment"

### **ARACRA Pipeline — User Manual**

**Version 2.0** | RNA-seq & TempO-Seq Analysis Platform

#### **Table of Contents**

1. [What is ARACRA?](https://euc-word-edit.officeapps.live.com/we/wordeditorframe.aspx?new=1&ui=en-US&rs=en-US&wdenableroaming=1&mscc=1&wdodb=1&hid=0B0F05A2-2059-C000-8565-3BB3825CD4F9.0&uih=sharepointcom&wdlcid=en-US&jsapi=1&jsapiver=v2&corrid=ce4cd52c-9b2c-c199-53a6-cbabf31cabe6&usid=ce4cd52c-9b2c-c199-53a6-cbabf31cabe6&newsession=1&sftc=1&uihit=docaspx&muv=1&ats=PairwiseBroker&cac=1&sams=1&mtf=1&sfp=1&sdp=1&hch=1&hwfh=1&wopisrc=https%3A%2F%2Frovira-my.sharepoint.com%2Fpersonal%2Fv7549573_epp_urv_cat%2F_vti_bin%2Fwopi.ashx%2Ffiles%2F4cf11d1fd11c4219b20c732a97ad725f&dchat=1&sc=%7B%22pmo%22%3A%22https%3A%2F%2Frovira-my.sharepoint.com%22%2C%22pmshare%22%3Atrue%7D&ctp=LeastProtected&rct=Normal&wdorigin=DocLib&wdhostclicktime=1774874517185&afdflight=91&csiro=1&wdredirectionreason=Unified_SingleFlush#1-what-is-aracra)
2. [What You'll Need](https://euc-word-edit.officeapps.live.com/we/wordeditorframe.aspx?new=1&ui=en-US&rs=en-US&wdenableroaming=1&mscc=1&wdodb=1&hid=0B0F05A2-2059-C000-8565-3BB3825CD4F9.0&uih=sharepointcom&wdlcid=en-US&jsapi=1&jsapiver=v2&corrid=ce4cd52c-9b2c-c199-53a6-cbabf31cabe6&usid=ce4cd52c-9b2c-c199-53a6-cbabf31cabe6&newsession=1&sftc=1&uihit=docaspx&muv=1&ats=PairwiseBroker&cac=1&sams=1&mtf=1&sfp=1&sdp=1&hch=1&hwfh=1&wopisrc=https%3A%2F%2Frovira-my.sharepoint.com%2Fpersonal%2Fv7549573_epp_urv_cat%2F_vti_bin%2Fwopi.ashx%2Ffiles%2F4cf11d1fd11c4219b20c732a97ad725f&dchat=1&sc=%7B%22pmo%22%3A%22https%3A%2F%2Frovira-my.sharepoint.com%22%2C%22pmshare%22%3Atrue%7D&ctp=LeastProtected&rct=Normal&wdorigin=DocLib&wdhostclicktime=1774874517185&afdflight=91&csiro=1&wdredirectionreason=Unified_SingleFlush#2-what-youll-need)
3. [First-Time Setup: Linux](https://euc-word-edit.officeapps.live.com/we/wordeditorframe.aspx?new=1&ui=en-US&rs=en-US&wdenableroaming=1&mscc=1&wdodb=1&hid=0B0F05A2-2059-C000-8565-3BB3825CD4F9.0&uih=sharepointcom&wdlcid=en-US&jsapi=1&jsapiver=v2&corrid=ce4cd52c-9b2c-c199-53a6-cbabf31cabe6&usid=ce4cd52c-9b2c-c199-53a6-cbabf31cabe6&newsession=1&sftc=1&uihit=docaspx&muv=1&ats=PairwiseBroker&cac=1&sams=1&mtf=1&sfp=1&sdp=1&hch=1&hwfh=1&wopisrc=https%3A%2F%2Frovira-my.sharepoint.com%2Fpersonal%2Fv7549573_epp_urv_cat%2F_vti_bin%2Fwopi.ashx%2Ffiles%2F4cf11d1fd11c4219b20c732a97ad725f&dchat=1&sc=%7B%22pmo%22%3A%22https%3A%2F%2Frovira-my.sharepoint.com%22%2C%22pmshare%22%3Atrue%7D&ctp=LeastProtected&rct=Normal&wdorigin=DocLib&wdhostclicktime=1774874517185&afdflight=91&csiro=1&wdredirectionreason=Unified_SingleFlush#3-first-time-setup-linux)
4. [First-Time Setup: Windows (via WSL2)](https://euc-word-edit.officeapps.live.com/we/wordeditorframe.aspx?new=1&ui=en-US&rs=en-US&wdenableroaming=1&mscc=1&wdodb=1&hid=0B0F05A2-2059-C000-8565-3BB3825CD4F9.0&uih=sharepointcom&wdlcid=en-US&jsapi=1&jsapiver=v2&corrid=ce4cd52c-9b2c-c199-53a6-cbabf31cabe6&usid=ce4cd52c-9b2c-c199-53a6-cbabf31cabe6&newsession=1&sftc=1&uihit=docaspx&muv=1&ats=PairwiseBroker&cac=1&sams=1&mtf=1&sfp=1&sdp=1&hch=1&hwfh=1&wopisrc=https%3A%2F%2Frovira-my.sharepoint.com%2Fpersonal%2Fv7549573_epp_urv_cat%2F_vti_bin%2Fwopi.ashx%2Ffiles%2F4cf11d1fd11c4219b20c732a97ad725f&dchat=1&sc=%7B%22pmo%22%3A%22https%3A%2F%2Frovira-my.sharepoint.com%22%2C%22pmshare%22%3Atrue%7D&ctp=LeastProtected&rct=Normal&wdorigin=DocLib&wdhostclicktime=1774874517185&afdflight=91&csiro=1&wdredirectionreason=Unified_SingleFlush#4-first-time-setup-windows-via-wsl2)
5. [Starting the App](https://euc-word-edit.officeapps.live.com/we/wordeditorframe.aspx?new=1&ui=en-US&rs=en-US&wdenableroaming=1&mscc=1&wdodb=1&hid=0B0F05A2-2059-C000-8565-3BB3825CD4F9.0&uih=sharepointcom&wdlcid=en-US&jsapi=1&jsapiver=v2&corrid=ce4cd52c-9b2c-c199-53a6-cbabf31cabe6&usid=ce4cd52c-9b2c-c199-53a6-cbabf31cabe6&newsession=1&sftc=1&uihit=docaspx&muv=1&ats=PairwiseBroker&cac=1&sams=1&mtf=1&sfp=1&sdp=1&hch=1&hwfh=1&wopisrc=https%3A%2F%2Frovira-my.sharepoint.com%2Fpersonal%2Fv7549573_epp_urv_cat%2F_vti_bin%2Fwopi.ashx%2Ffiles%2F4cf11d1fd11c4219b20c732a97ad725f&dchat=1&sc=%7B%22pmo%22%3A%22https%3A%2F%2Frovira-my.sharepoint.com%22%2C%22pmshare%22%3Atrue%7D&ctp=LeastProtected&rct=Normal&wdorigin=DocLib&wdhostclicktime=1774874517185&afdflight=91&csiro=1&wdredirectionreason=Unified_SingleFlush#5-starting-the-app)
6. [Web Interface Walkthrough](https://euc-word-edit.officeapps.live.com/we/wordeditorframe.aspx?new=1&ui=en-US&rs=en-US&wdenableroaming=1&mscc=1&wdodb=1&hid=0B0F05A2-2059-C000-8565-3BB3825CD4F9.0&uih=sharepointcom&wdlcid=en-US&jsapi=1&jsapiver=v2&corrid=ce4cd52c-9b2c-c199-53a6-cbabf31cabe6&usid=ce4cd52c-9b2c-c199-53a6-cbabf31cabe6&newsession=1&sftc=1&uihit=docaspx&muv=1&ats=PairwiseBroker&cac=1&sams=1&mtf=1&sfp=1&sdp=1&hch=1&hwfh=1&wopisrc=https%3A%2F%2Frovira-my.sharepoint.com%2Fpersonal%2Fv7549573_epp_urv_cat%2F_vti_bin%2Fwopi.ashx%2Ffiles%2F4cf11d1fd11c4219b20c732a97ad725f&dchat=1&sc=%7B%22pmo%22%3A%22https%3A%2F%2Frovira-my.sharepoint.com%22%2C%22pmshare%22%3Atrue%7D&ctp=LeastProtected&rct=Normal&wdorigin=DocLib&wdhostclicktime=1774874517185&afdflight=91&csiro=1&wdredirectionreason=Unified_SingleFlush#6-web-interface-walkthrough)
   a. [The Sidebar](https://euc-word-edit.officeapps.live.com/we/wordeditorframe.aspx?new=1&ui=en-US&rs=en-US&wdenableroaming=1&mscc=1&wdodb=1&hid=0B0F05A2-2059-C000-8565-3BB3825CD4F9.0&uih=sharepointcom&wdlcid=en-US&jsapi=1&jsapiver=v2&corrid=ce4cd52c-9b2c-c199-53a6-cbabf31cabe6&usid=ce4cd52c-9b2c-c199-53a6-cbabf31cabe6&newsession=1&sftc=1&uihit=docaspx&muv=1&ats=PairwiseBroker&cac=1&sams=1&mtf=1&sfp=1&sdp=1&hch=1&hwfh=1&wopisrc=https%3A%2F%2Frovira-my.sharepoint.com%2Fpersonal%2Fv7549573_epp_urv_cat%2F_vti_bin%2Fwopi.ashx%2Ffiles%2F4cf11d1fd11c4219b20c732a97ad725f&dchat=1&sc=%7B%22pmo%22%3A%22https%3A%2F%2Frovira-my.sharepoint.com%22%2C%22pmshare%22%3Atrue%7D&ctp=LeastProtected&rct=Normal&wdorigin=DocLib&wdhostclicktime=1774874517185&afdflight=91&csiro=1&wdredirectionreason=Unified_SingleFlush#61-the-sidebar)
   b. [Full Pipeline Tab](https://euc-word-edit.officeapps.live.com/we/wordeditorframe.aspx?new=1&ui=en-US&rs=en-US&wdenableroaming=1&mscc=1&wdodb=1&hid=0B0F05A2-2059-C000-8565-3BB3825CD4F9.0&uih=sharepointcom&wdlcid=en-US&jsapi=1&jsapiver=v2&corrid=ce4cd52c-9b2c-c199-53a6-cbabf31cabe6&usid=ce4cd52c-9b2c-c199-53a6-cbabf31cabe6&newsession=1&sftc=1&uihit=docaspx&muv=1&ats=PairwiseBroker&cac=1&sams=1&mtf=1&sfp=1&sdp=1&hch=1&hwfh=1&wopisrc=https%3A%2F%2Frovira-my.sharepoint.com%2Fpersonal%2Fv7549573_epp_urv_cat%2F_vti_bin%2Fwopi.ashx%2Ffiles%2F4cf11d1fd11c4219b20c732a97ad725f&dchat=1&sc=%7B%22pmo%22%3A%22https%3A%2F%2Frovira-my.sharepoint.com%22%2C%22pmshare%22%3Atrue%7D&ctp=LeastProtected&rct=Normal&wdorigin=DocLib&wdhostclicktime=1774874517185&afdflight=91&csiro=1&wdredirectionreason=Unified_SingleFlush#62-full-pipeline-tab)
   c. [DESeq2 / DRomics Tab](https://euc-word-edit.officeapps.live.com/we/wordeditorframe.aspx?new=1&ui=en-US&rs=en-US&wdenableroaming=1&mscc=1&wdodb=1&hid=0B0F05A2-2059-C000-8565-3BB3825CD4F9.0&uih=sharepointcom&wdlcid=en-US&jsapi=1&jsapiver=v2&corrid=ce4cd52c-9b2c-c199-53a6-cbabf31cabe6&usid=ce4cd52c-9b2c-c199-53a6-cbabf31cabe6&newsession=1&sftc=1&uihit=docaspx&muv=1&ats=PairwiseBroker&cac=1&sams=1&mtf=1&sfp=1&sdp=1&hch=1&hwfh=1&wopisrc=https%3A%2F%2Frovira-my.sharepoint.com%2Fpersonal%2Fv7549573_epp_urv_cat%2F_vti_bin%2Fwopi.ashx%2Ffiles%2F4cf11d1fd11c4219b20c732a97ad725f&dchat=1&sc=%7B%22pmo%22%3A%22https%3A%2F%2Frovira-my.sharepoint.com%22%2C%22pmshare%22%3Atrue%7D&ctp=LeastProtected&rct=Normal&wdorigin=DocLib&wdhostclicktime=1774874517185&afdflight=91&csiro=1&wdredirectionreason=Unified_SingleFlush#63-deseq2--dromics-tab)
   d. [Parameters Tab](https://euc-word-edit.officeapps.live.com/we/wordeditorframe.aspx?new=1&ui=en-US&rs=en-US&wdenableroaming=1&mscc=1&wdodb=1&hid=0B0F05A2-2059-C000-8565-3BB3825CD4F9.0&uih=sharepointcom&wdlcid=en-US&jsapi=1&jsapiver=v2&corrid=ce4cd52c-9b2c-c199-53a6-cbabf31cabe6&usid=ce4cd52c-9b2c-c199-53a6-cbabf31cabe6&newsession=1&sftc=1&uihit=docaspx&muv=1&ats=PairwiseBroker&cac=1&sams=1&mtf=1&sfp=1&sdp=1&hch=1&hwfh=1&wopisrc=https%3A%2F%2Frovira-my.sharepoint.com%2Fpersonal%2Fv7549573_epp_urv_cat%2F_vti_bin%2Fwopi.ashx%2Ffiles%2F4cf11d1fd11c4219b20c732a97ad725f&dchat=1&sc=%7B%22pmo%22%3A%22https%3A%2F%2Frovira-my.sharepoint.com%22%2C%22pmshare%22%3Atrue%7D&ctp=LeastProtected&rct=Normal&wdorigin=DocLib&wdhostclicktime=1774874517185&afdflight=91&csiro=1&wdredirectionreason=Unified_SingleFlush#64-parameters-tab)
   e. [Results Tab](https://euc-word-edit.officeapps.live.com/we/wordeditorframe.aspx?new=1&ui=en-US&rs=en-US&wdenableroaming=1&mscc=1&wdodb=1&hid=0B0F05A2-2059-C000-8565-3BB3825CD4F9.0&uih=sharepointcom&wdlcid=en-US&jsapi=1&jsapiver=v2&corrid=ce4cd52c-9b2c-c199-53a6-cbabf31cabe6&usid=ce4cd52c-9b2c-c199-53a6-cbabf31cabe6&newsession=1&sftc=1&uihit=docaspx&muv=1&ats=PairwiseBroker&cac=1&sams=1&mtf=1&sfp=1&sdp=1&hch=1&hwfh=1&wopisrc=https%3A%2F%2Frovira-my.sharepoint.com%2Fpersonal%2Fv7549573_epp_urv_cat%2F_vti_bin%2Fwopi.ashx%2Ffiles%2F4cf11d1fd11c4219b20c732a97ad725f&dchat=1&sc=%7B%22pmo%22%3A%22https%3A%2F%2Frovira-my.sharepoint.com%22%2C%22pmshare%22%3Atrue%7D&ctp=LeastProtected&rct=Normal&wdorigin=DocLib&wdhostclicktime=1774874517185&afdflight=91&csiro=1&wdredirectionreason=Unified_SingleFlush#65-results-tab)
7. [Your Metadata File](https://euc-word-edit.officeapps.live.com/we/wordeditorframe.aspx?new=1&ui=en-US&rs=en-US&wdenableroaming=1&mscc=1&wdodb=1&hid=0B0F05A2-2059-C000-8565-3BB3825CD4F9.0&uih=sharepointcom&wdlcid=en-US&jsapi=1&jsapiver=v2&corrid=ce4cd52c-9b2c-c199-53a6-cbabf31cabe6&usid=ce4cd52c-9b2c-c199-53a6-cbabf31cabe6&newsession=1&sftc=1&uihit=docaspx&muv=1&ats=PairwiseBroker&cac=1&sams=1&mtf=1&sfp=1&sdp=1&hch=1&hwfh=1&wopisrc=https%3A%2F%2Frovira-my.sharepoint.com%2Fpersonal%2Fv7549573_epp_urv_cat%2F_vti_bin%2Fwopi.ashx%2Ffiles%2F4cf11d1fd11c4219b20c732a97ad725f&dchat=1&sc=%7B%22pmo%22%3A%22https%3A%2F%2Frovira-my.sharepoint.com%22%2C%22pmshare%22%3Atrue%7D&ctp=LeastProtected&rct=Normal&wdorigin=DocLib&wdhostclicktime=1774874517185&afdflight=91&csiro=1&wdredirectionreason=Unified_SingleFlush#7-your-metadata-file)
8. [Understanding the Two-Phase Design](https://euc-word-edit.officeapps.live.com/we/wordeditorframe.aspx?new=1&ui=en-US&rs=en-US&wdenableroaming=1&mscc=1&wdodb=1&hid=0B0F05A2-2059-C000-8565-3BB3825CD4F9.0&uih=sharepointcom&wdlcid=en-US&jsapi=1&jsapiver=v2&corrid=ce4cd52c-9b2c-c199-53a6-cbabf31cabe6&usid=ce4cd52c-9b2c-c199-53a6-cbabf31cabe6&newsession=1&sftc=1&uihit=docaspx&muv=1&ats=PairwiseBroker&cac=1&sams=1&mtf=1&sfp=1&sdp=1&hch=1&hwfh=1&wopisrc=https%3A%2F%2Frovira-my.sharepoint.com%2Fpersonal%2Fv7549573_epp_urv_cat%2F_vti_bin%2Fwopi.ashx%2Ffiles%2F4cf11d1fd11c4219b20c732a97ad725f&dchat=1&sc=%7B%22pmo%22%3A%22https%3A%2F%2Frovira-my.sharepoint.com%22%2C%22pmshare%22%3Atrue%7D&ctp=LeastProtected&rct=Normal&wdorigin=DocLib&wdhostclicktime=1774874517185&afdflight=91&csiro=1&wdredirectionreason=Unified_SingleFlush#8-understanding-the-two-phase-design)
9. [Bioinformatics Tools Reference](https://euc-word-edit.officeapps.live.com/we/wordeditorframe.aspx?new=1&ui=en-US&rs=en-US&wdenableroaming=1&mscc=1&wdodb=1&hid=0B0F05A2-2059-C000-8565-3BB3825CD4F9.0&uih=sharepointcom&wdlcid=en-US&jsapi=1&jsapiver=v2&corrid=ce4cd52c-9b2c-c199-53a6-cbabf31cabe6&usid=ce4cd52c-9b2c-c199-53a6-cbabf31cabe6&newsession=1&sftc=1&uihit=docaspx&muv=1&ats=PairwiseBroker&cac=1&sams=1&mtf=1&sfp=1&sdp=1&hch=1&hwfh=1&wopisrc=https%3A%2F%2Frovira-my.sharepoint.com%2Fpersonal%2Fv7549573_epp_urv_cat%2F_vti_bin%2Fwopi.ashx%2Ffiles%2F4cf11d1fd11c4219b20c732a97ad725f&dchat=1&sc=%7B%22pmo%22%3A%22https%3A%2F%2Frovira-my.sharepoint.com%22%2C%22pmshare%22%3Atrue%7D&ctp=LeastProtected&rct=Normal&wdorigin=DocLib&wdhostclicktime=1774874517185&afdflight=91&csiro=1&wdredirectionreason=Unified_SingleFlush#9-bioinformatics-tools-reference)
10. [Output Files & Folders](https://euc-word-edit.officeapps.live.com/we/wordeditorframe.aspx?new=1&ui=en-US&rs=en-US&wdenableroaming=1&mscc=1&wdodb=1&hid=0B0F05A2-2059-C000-8565-3BB3825CD4F9.0&uih=sharepointcom&wdlcid=en-US&jsapi=1&jsapiver=v2&corrid=ce4cd52c-9b2c-c199-53a6-cbabf31cabe6&usid=ce4cd52c-9b2c-c199-53a6-cbabf31cabe6&newsession=1&sftc=1&uihit=docaspx&muv=1&ats=PairwiseBroker&cac=1&sams=1&mtf=1&sfp=1&sdp=1&hch=1&hwfh=1&wopisrc=https%3A%2F%2Frovira-my.sharepoint.com%2Fpersonal%2Fv7549573_epp_urv_cat%2F_vti_bin%2Fwopi.ashx%2Ffiles%2F4cf11d1fd11c4219b20c732a97ad725f&dchat=1&sc=%7B%22pmo%22%3A%22https%3A%2F%2Frovira-my.sharepoint.com%22%2C%22pmshare%22%3Atrue%7D&ctp=LeastProtected&rct=Normal&wdorigin=DocLib&wdhostclicktime=1774874517185&afdflight=91&csiro=1&wdredirectionreason=Unified_SingleFlush#10-output-files--folders)
11. [Troubleshooting](https://euc-word-edit.officeapps.live.com/we/wordeditorframe.aspx?new=1&ui=en-US&rs=en-US&wdenableroaming=1&mscc=1&wdodb=1&hid=0B0F05A2-2059-C000-8565-3BB3825CD4F9.0&uih=sharepointcom&wdlcid=en-US&jsapi=1&jsapiver=v2&corrid=ce4cd52c-9b2c-c199-53a6-cbabf31cabe6&usid=ce4cd52c-9b2c-c199-53a6-cbabf31cabe6&newsession=1&sftc=1&uihit=docaspx&muv=1&ats=PairwiseBroker&cac=1&sams=1&mtf=1&sfp=1&sdp=1&hch=1&hwfh=1&wopisrc=https%3A%2F%2Frovira-my.sharepoint.com%2Fpersonal%2Fv7549573_epp_urv_cat%2F_vti_bin%2Fwopi.ashx%2Ffiles%2F4cf11d1fd11c4219b20c732a97ad725f&dchat=1&sc=%7B%22pmo%22%3A%22https%3A%2F%2Frovira-my.sharepoint.com%22%2C%22pmshare%22%3Atrue%7D&ctp=LeastProtected&rct=Normal&wdorigin=DocLib&wdhostclicktime=1774874517185&afdflight=91&csiro=1&wdredirectionreason=Unified_SingleFlush#11-troubleshooting)

#### **1. What is ARACRA?**

ARACRA is a Streamlit-based graphical user interface that integrates a complete RNA-seq processing pipeline with downstream differential expression (DEG) and dose-response (DRomics) analysis.

**What it does, start to finish:**

```
Raw reads (SRA or local FASTQ)
    ↓  fastp          — quality trimming & adapter removal
    ↓  STAR / HISAT2  — alignment to the human genome (hg38)
    ↓  QC suite       — flagstat, RSeQC, Picard, Qualimap, MultiQC
    ↓  featureCounts / Salmon  — gene-level quantification
    ↓  [you review QC, decide which samples to keep]
    ↓  DESeq2         — differential expression analysis
    ↓  DRomics / BMD  — dose-response modelling & benchmark dose

```

It also supports **TempO-Seq** (BioSpyder targeted sequencing) alongside standard whole-transcriptome RNA-seq.

#### **2. What You'll Need**

##### **Hardware minimums**

| **Resource** | **Minimum** | **Recommended** |
| --- | --- | --- |
| RAM | 16 GB | 64 GB+ |
| CPU cores | 4 | 16+ |
| Free disk space | 100 GB | 500 GB+ |
| Internet connection | Required for first setup | — |

**RAM note:** STAR alignment requires ~32 GB RAM to load the human genome index. If your machine has less than 32 GB, the setup script will automatically recommend **HISAT2**, which only needs ~8 GB RAM. You won't lose any important functionality — HISAT2 is a solid, widely used aligner.

##### **Operating system**

- **Linux** (Ubuntu 20.04+ or any modern distro) — runs natively
- **Windows 10/11** — runs through WSL2 (Windows Subsystem for Linux)
- **macOS** — technically possible but not officially tested

#### **3. First-Time Setup: Linux**

This section assumes you're starting from a fresh Linux machine with no special software installed. Everything else will be installed automatically.

##### **Step 1 — Get the code**

Open a terminal and run:

```
# If you haven't already, install git
sudo apt update && sudo apt install -y git

# Clone (or download and extract) the ARACRA folder
cd ~/Downloads
# (the ARACRA folder should already be here if you received it directly)
cd ARACRA

```

##### **Step 2 — Check your disk space**

The setup downloads about 10–15 GB of reference genome data. Make sure you have room:

df -h ~

If you want to store the databases somewhere other than ~/databases (e.g. a large external drive), note the path — you'll pass it to setup with --db-dir.

##### **Step 3 — Run setup**

bash setup.sh

The setup script will:

1. Detect your CPU count, RAM, and available disk
2. Automatically install **Miniforge3** (a lightweight conda) if conda isn't already present
3. Create a conda environment called test_ARACRA with every tool pre-installed
4. Download the **hg38 human genome**, GTF annotation, transcriptome FASTA, and all annotation files
5. Build alignment indexes (STAR takes 30–45 min; HISAT2 index is downloaded pre-built)
6. Build the Salmon index with genome decoys
7. Install all required **R packages** including DESeq2, DRomics, edgeR, and pathway databases

**Expect this to take 1–3 hours** the first time, mostly waiting for downloads and index building.

###### **Optional setup flags**

```
# Skip building alignment indexes (if you already have them elsewhere)
bash setup.sh --skip-index

# Skip downloading FastQ Screen genomes (saves ~14 GB, disables contamination screening)
bash setup.sh --skip-screen

# Store databases on a different drive
bash setup.sh --db-dir=/mnt/bigdrive/databases

# Combine flags
bash setup.sh --skip-screen --db-dir=/mnt/bigdrive/databases

```

##### **Step 4 — Verify the install**

After setup completes, you should see a summary like:

```
 ✔ nextflow       version 24.x.x
  ✔ STAR           2.7.x
  ✔ hisat2         2.2.x
  ✔ samtools       1.x
  ✔ fastp          0.23.x
  ✔ featureCounts  2.x
  ✔ salmon         1.10.x
  ✔ multiqc        1.x
  ✔ Rscript        4.x
  ✔ streamlit      1.x

```

If any tool shows ✘ NOT FOUND, see the [Troubleshooting](https://euc-word-edit.officeapps.live.com/we/wordeditorframe.aspx?new=1&ui=en-US&rs=en-US&wdenableroaming=1&mscc=1&wdodb=1&hid=0B0F05A2-2059-C000-8565-3BB3825CD4F9.0&uih=sharepointcom&wdlcid=en-US&jsapi=1&jsapiver=v2&corrid=ce4cd52c-9b2c-c199-53a6-cbabf31cabe6&usid=ce4cd52c-9b2c-c199-53a6-cbabf31cabe6&newsession=1&sftc=1&uihit=docaspx&muv=1&ats=PairwiseBroker&cac=1&sams=1&mtf=1&sfp=1&sdp=1&hch=1&hwfh=1&wopisrc=https%3A%2F%2Frovira-my.sharepoint.com%2Fpersonal%2Fv7549573_epp_urv_cat%2F_vti_bin%2Fwopi.ashx%2Ffiles%2F4cf11d1fd11c4219b20c732a97ad725f&dchat=1&sc=%7B%22pmo%22%3A%22https%3A%2F%2Frovira-my.sharepoint.com%22%2C%22pmshare%22%3Atrue%7D&ctp=LeastProtected&rct=Normal&wdorigin=DocLib&wdhostclicktime=1774874517185&afdflight=91&csiro=1&wdredirectionreason=Unified_SingleFlush#11-troubleshooting) section.

#### **4. First-Time Setup: Windows (via WSL2)**

Windows doesn't run Linux tools natively, but **WSL2** (Windows Subsystem for Linux 2) gives you a full Linux environment right inside Windows — no virtual machine needed. ARACRA runs inside WSL2 exactly like it would on a Linux machine.

##### **Step 1 — Enable WSL2**

Open **PowerShell as Administrator** (right-click the Start menu → "Windows PowerShell (Admin)") and run:

wsl --install

This installs WSL2 with Ubuntu. **Restart your computer** when prompted.

If you already have WSL1, upgrade it:

wsl --set-default-version 2

##### **Step 2 — Set up Ubuntu**

After restart, search "Ubuntu" in the Start menu and open it. The first time it runs, it will ask you to create a username and password — choose something simple, since you'll type the password occasionally.

```
# Update the package list
sudo apt update && sudo apt upgrade -y

# Install some basics
sudo apt install -y git curl wget unzip build-essential

```

##### **Step 3 — Allocate enough RAM to WSL2**

By default, WSL2 caps itself at 50% of your system RAM. If you want to use STAR (which needs ~32 GB), you may need to raise this limit.

Create or edit the file C:\Users\YourName\.wslconfig (replace YourName with your Windows username):

[wsl2]
memory=48GB
processors=12
swap=8GB

Then restart WSL from PowerShell:

wsl --shutdown

Re-open Ubuntu. Run free -h to confirm the RAM is now available.

##### **Step 4 — Decide where to put your data**

WSL2 has its own filesystem, separate from Windows. Your Windows C:\ drive is mounted at /mnt/c/ inside WSL. For best performance, keep your data **inside WSL** (e.g. ~/Downloads/ARACRA), not on the Windows drive — filesystem crossover is slow.

```
# Copy files from Windows into WSL (if needed)
cp -r /mnt/c/Users/YourName/Downloads/ARACRA ~/Downloads/ARACRA

```

##### **Step 5 — Run setup (same as Linux)**

cd ~/Downloads/ARACRA
bash setup.sh

Everything from this point forward is identical to the Linux instructions. The setup script handles everything automatically.

##### **Step 6 — Accessing the web interface from Windows**

When you launch the app inside WSL2, the Streamlit server starts at <http://localhost:8501>. Open any Windows browser (Chrome, Edge, Firefox) and go to:

<http://localhost:8501>

It works. WSL2 forwards ports to Windows automatically.

**Tip:** If the page doesn't load, try <http://127.0.0.1:8501> instead.

#### **5. Starting the App**

Each time you want to use ARACRA (after the one-time setup is done):

cd ~/Downloads/ARACRA
bash run_app.sh

You'll see output like:

```
 You can now view your Streamlit app in your browser.
  Local URL: http://localhost:8501

```

Open that URL in your browser. Leave the terminal running in the background — the pipeline continues even if you close the browser tab.

**The pipeline keeps running even if you close the browser.** It runs as a detached background process. You can come back hours later, re-open the app, and see your results.

#### **6. Web Interface Walkthrough**

The ARACRA interface has two main areas: the **sidebar** on the left for configuration, and **four tabs** in the main area for different stages of your analysis.

##### **6.1 The Sidebar**

The sidebar is your control panel. It stays visible from every tab.

###### **System Info Box**

At the very top, a small box shows your machine's live stats:

```
System
  RAM:   62.8 GB  (58.2 GB free)
  CPUs:  16
  Disk:  420 GB free
  Aligner: STAR

```

The **Aligner** line tells you which aligner the setup script recommended based on your RAM. You can override this in the Full Pipeline tab.

###### **Reference Files**

These paths are auto-populated from your setup. You usually don't need to touch them unless you want to use a custom reference.

| **Field** | **What it points to** |
| --- | --- |
| STAR index | Pre-built STAR genome index (~28 GB) |
| HISAT2 index | Pre-built HISAT2 splice-aware index |
| Salmon index | Selective alignment index with decoys |
| GTF annotation | GENCODE v44 gene annotation |
| RefSeq BED | Gene body coordinates for RSeQC |
| Housekeeping BED | Housekeeping gene coordinates for coverage QC |
| refFlat | Picard RNA metrics reference |
| FastQ Screen conf | Contamination screen genome list |

###### **Directories**

- **Output directory** — where results, count matrices, QC reports, and analysis outputs are saved. Default: ~/aracra_star_work/results
- **Work directory** — Nextflow's internal scratch space for intermediate files. Default: ~/aracra_star_work/work

###### **Executor**

- **Profile** — standard runs everything locally on your machine. slurm submits jobs to an HPC cluster (requires a configured SLURM environment).
- **Library layout** — SE for single-end reads, PE for paired-end reads.
- **Resume previous run** — Keep this checked. It tells Nextflow to skip steps that already completed successfully, so if something fails partway through, you restart without repeating hours of alignment work.

###### **Platform**

- **RNA-seq (whole transcriptome)** — standard mode; aligns to the full hg38 genome
- **TempO-Seq (targeted)** — for BioSpyder TempO-Seq data; aligns to a probe reference instead of the whole genome. When selected, a file uploader appears for your **BioSpyder manifest CSV**.

###### **Downstream**

Checkboxes to enable/disable the analysis steps that run after quantification:

- **DESeq2** — differential expression analysis (treatment vs. control)
- **DRomics / BMD** — dose-response modelling and benchmark dose calculation

###### **System Check**

Click **▶ Check Tools** to verify that every required tool is properly installed and accessible. A green checkmark means it's working; a red ✘ means something is wrong.

This is the first thing to check if your pipeline fails unexpectedly.

##### **6.2 Full Pipeline Tab**

This is where most users spend their time. It runs the complete pipeline from raw reads to quantified counts, with an optional QC review gate before statistical analysis.

###### **FASTQ Source**

Choose where your sequencing data comes from:

- **Download from SRA** — enter SRA accession numbers (e.g. SRR12345678) directly in the app; the pipeline downloads them automatically using prefetch and fasterq-dump
- **Local raw FASTQs** — you already have FASTQ files; provide the folder path
- **Local pre-trimmed FASTQs** — your reads are already trimmed with fastp/Trimmomatic; skip the trimming step

###### **Metadata Upload**

Upload your sample metadata as a CSV or Excel file. This tells the pipeline which samples belong to which treatment groups, what doses were used, and which samples are controls.

At minimum, your file needs columns for:

| **Column** | **Purpose** |
| --- | --- |
| Sample_Name | SRR accession or file basename |
| Treatment | Chemical or treatment name |
| Dose | Dose level (numeric, in µM or your units) |
| Type | treatment or control |

The app auto-detects common column names (e.g. sample, chemical, conc, condition) and maps them. A column-mapping interface appears so you can confirm or correct the mapping.

###### **Aligner Selection**

Choose between **STAR** and **HISAT2**:

- **STAR** — faster, more accurate for standard RNA-seq; needs ~32 GB RAM
- **HISAT2** — uses only ~8 GB RAM; excellent for machines with less memory

###### **Optional QC Modules**

Toggle additional QC steps on or off:

| **Module** | **What it does** | **Cost** |
| --- | --- | --- |
| FastQ Screen | Screens for non-human contamination | Moderate time |
| Read distribution | RSeQC read distribution across gene features | Fast |
| Qualimap | Deep BAM QC including RNA-seq metrics | Slow |
| Gene body coverage (housekeeping) | 5'→3' coverage uniformity on stable genes | Fast |
| Gene body coverage (full) | Coverage across all genes | Moderate |
| Picard RNA metrics | Ribosomal fraction, 5'/3' bias | Fast |
| Salmon (alignment-based) | Transcript-level quantification | Moderate |
| Salmon (pseudoalignment) | Ultra-fast pseudo-mapping | Fast |

###### **Quantification**

- **featureCounts** — gene-level counts from BAM files; enabled by default
- **Salmon (aligned)** — uses the BAM file for accurate transcript quantification
- **Infer strandedness** — auto-detects library strandedness using RSeQC; recommended

###### **Run Controls**

- **▶ Start Pipeline** — launches Phase 1 (preprocessing + QC + quantification)
- **⏹ Stop** — gracefully terminates the running pipeline
- **Progress bars** — show per-process completion (Download → Fastp → STAR → QC → featureCounts → MultiQC)
- **Live log** — streams the Nextflow log in real time; last 80 lines shown

###### **QC Review Gate**

After Phase 1 completes, the app shows a **QC review panel** before any statistical analysis runs. This is intentional — you get to see the data quality and decide which samples to include.

The panel shows two layers of outlier detection:

**Layer 1 — Alignment QC** (Grubbs test + IQR fences):

- Flags samples with abnormally low mapping rates, too few reads, or excessive multi-mapping
- A sample is only flagged if *both* the Grubbs test and IQR method agree

**Layer 2 — Expression PCA**:

- Runs PCA on the count matrix and flags samples that cluster far from their treatment group
- A scatter plot shows PC1 vs PC2, with outliers marked

Flagged samples are **pre-ticked for exclusion**, but you make the final call. Untick a sample to keep it. Tick a sample that wasn't auto-flagged if you want to remove it manually.

Once you're happy with your selection, click **▶ Run Phase 2 (DESeq2 / DRomics)** to launch the statistical analysis.

##### **6.3 DESeq2 / DRomics Tab**

This tab lets you run just the statistical analysis — useful when you already have a count matrix from a previous run (or from another pipeline entirely).

###### **Direct Analysis Mode**

Upload a count matrix CSV/TSV directly. The first column should be gene IDs; remaining columns are samples. No alignment needed.

###### **QC Check (Step 1)**

Before running DESeq2, you can run a standalone PCA QC check on your count matrix. This generates the PCA plot and outlier report so you can review sample quality before committing to the full analysis.

###### **Treatment & Control Selection**

Dropdowns to select which treatment group to compare against which control. The run is named automatically (e.g. BisphenolA_vs_Control).

###### **Launch Analysis**

Click **▶ Run DESeq2** or **▶ Run DRomics** (or both). The analysis runs in a few minutes for typical datasets.

##### **6.4 Parameters Tab**

Fine-tune the statistical parameters before running your analysis. Sensible defaults are pre-filled; most users won't need to change these.

###### **DESeq2 Parameters**

| **Parameter** | **Default** | **Meaning** |
| --- | --- | --- |
| FDR (strict) | 0.01 | Adjusted p-value cutoff for the "stringent" DEG list |
| FDR (relaxed) | 0.05 | Adjusted p-value cutoff for the main DEG list |
| log₂ fold-change threshold | 1.0 | Minimum fold-change magnitude (log₂ scale) for a gene to be called DE |
| Read count threshold | 1,000,000 | Samples below this read count are flagged as low-quality |

###### **DRomics / BMD Parameters**

| **Parameter** | **Default** | **Meaning** |
| --- | --- | --- |
| FDR (gene selection) | 0.05 | Significance threshold for selecting dose-responsive genes |
| Model selection criterion | AICc | Information criterion for choosing between dose-response models |
| Transformation method | auto | Data transformation before fitting (auto, log, sqrt, none) |
| BMD z-factor | 1.0 | Number of standard deviations from baseline for BMD definition |
| BMD x-factor | 10.0 | Percentage change from baseline for BMD definition (alternative) |
| Bootstrap iterations | 1000 | Iterations for BMD confidence interval estimation |
| Use MSigDB pathways | Yes | Include MSigDB hallmark/KEGG/GO pathway sets for tPOD calculation |

###### **NTP 2018 / EFSA Quality Filters**

These filters clean up BMD estimates before calculating the tPOD:

| **Filter** | **Default** | **Purpose** |
| --- | --- | --- |
| Remove BMDs > max dose | On | Removes BMDs that exceed the highest tested dose (extrapolation) |
| Extrapolation factor | 10× | Flags BMDs more than 10× below the lowest tested dose |
| BMDU/BMDL ratio | 40 | Removes genes with very wide confidence intervals (uncertain BMDs) |
| Min fold-change | 0.0 | Optional: remove genes with small effect sizes |

##### **6.5 Results Tab**

Once a run completes, the Results tab shows a history of all your analysis runs and their outputs.

###### **DESeq2 Results**

- **Metric cards** — samples analyzed, total genes tested, DEGs at FDR < 0.05, upregulated count, downregulated count
- **PCA plot** — batch-corrected VST-normalized PCA showing sample clustering
- **Volcano plot** — fold-change vs. significance; colored by up/down/non-significant
- **DEG table** — top differentially expressed genes with fold-change, p-value, and adjusted p-value
- **Download buttons** — export All Results, Filtered DEGs, Upregulated, or Downregulated gene lists as CSV

###### **DRomics / BMD Results**

- **Metric cards** — genes analyzed, dose-responsive genes, models fitted, median gene BMD, tPOD
- **Model quality panel** — median pseudo-R², signal-to-noise ratio, percentage of genes with R² > 0.5
- **tPOD highlight box** — the transcriptomic point of departure with the driving pathway
- **All tPOD methods table** — compares tPOD estimates from multiple calculation methods (NTP 2018, median, Q25, etc.)
- **tPOD by gene set collection** — breakdown by KEGG, GO Biological Process, GO Molecular Function, Hallmarks
- **Top 5 most sensitive pathways** — the pathways with the lowest median BMD
- **Dose-response curves** — fitted curves for top responding genes
- **BMD distribution** — histogram of all gene-level BMD estimates
- **Pathway sensitivity plot** — ranked pathway tPOD values
- **Download buttons** — BMD results, top sensitive genes, dose-response models, pathway summary, model quality metrics

#### **7. Your Metadata File**

The metadata file is the key input that connects your samples to their experimental context. Get this right and everything else follows.

##### **Required format**

A CSV or Excel file with one row per sample:

Sample_Name,Treatment,Dose,Type,Batch
SRR12345678,Bisphenol A,0.1,treatment,1
SRR12345679,Bisphenol A,1.0,treatment,1
SRR12345680,Bisphenol A,10.0,treatment,1
SRR12345681,Control,0,control,1
SRR12345682,Control,0,control,1
SRR12345683,Control,0,control,1

##### **Column guide**

- **Sample_Name** — must match your SRR accession exactly (for SRA downloads), or the FASTQ filename without the .fastq.gz extension (for local files)
- **Treatment** — the chemical or treatment name. Identical strings = same group. Spaces are fine.
- **Dose** — numeric. Use the same units throughout (µM is conventional). Controls should have dose = 0.
- **Type** — treatment for treated samples, control for vehicle/untreated controls
- **Batch** *(optional)* — integer batch number if samples were processed in multiple batches. DESeq2 will include batch as a covariate in the design.

##### **Tips**

- Column names are flexible — the app will auto-detect sample, chemical, conc, group, etc. and map them. You'll see a confirmation screen before the pipeline starts.
- If you have multiple treatment groups, include all of them in one file. You'll select the specific treatment vs. control comparison when running DESeq2.
- Dose values for DRomics must be numeric and monotonically increasing. Categorical dose labels won't work.

#### **8. Understanding the Two-Phase Design**

ARACRA deliberately splits the pipeline into two phases with a **human review step in between**. This design choice is intentional and important.

##### **Phase 1 — Preprocessing & QC**

Runs automatically without intervention:

1. Download raw reads from SRA (if needed)
2. Trim adapters and low-quality bases with fastp
3. Screen for contamination with FastQ Screen
4. Align to hg38 with STAR or HISAT2
5. Collect alignment statistics (flagstat, idxstats)
6. Detect library strandedness
7. Measure gene body coverage
8. Run Picard and/or Qualimap RNA-seq QC
9. Quantify gene expression (featureCounts, Salmon)
10. Compile everything into a MultiQC report

##### **QC Review — Your Turn**

After Phase 1, the pipeline **pauses** and shows you the quality metrics.

You see the PCA plot of all samples and a per-sample table. Pre-ticked samples are the auto-detected outliers.

Review carefully:

- Are the flagged samples genuinely bad, or just naturally variable?
- Are there samples you want to remove that weren't auto-flagged?
- Do the PCA clusters make biological sense?

##### **Phase 2 — Statistical Analysis**

After you confirm your sample selection, Phase 2 runs DESeq2 and DRomics on the approved samples.

This two-phase approach prevents a common problem: running statistical analysis on bad data and not realizing it until much later. Every decision is visible and recorded.

#### **9. Bioinformatics Tools Reference**

All tools are installed automatically by setup.sh. Here's what each one does:

##### **Preprocessing**

| **Tool** | **Purpose** |
| --- | --- |
| **SRA Tools** (prefetch + fasterq-dump) | Downloads reads from NCBI's Sequence Read Archive |
| **fastp** | Adapter trimming, quality filtering, per-sample QC reports |
| **FastQ Screen** | Checks reads against 13 reference genomes to detect contamination |

##### **Alignment**

| **Tool** | **Purpose** |
| --- | --- |
| **STAR** | Fast splice-aware aligner; needs 32 GB RAM; gold standard for RNA-seq |
| **HISAT2** | Splice-aware aligner; needs only 8 GB RAM; excellent alternative |
| **Bowtie2** | Used internally by FastQ Screen for contamination screening |

##### **Post-alignment QC**

| **Tool** | **Purpose** |
| --- | --- |
| **samtools** | Sorts, indexes BAM files; generates flagstat and idxstats reports |
| **RSeQC** | Gene body coverage, read distribution, strandedness inference |
| **Picard** | RNA-seq metrics: ribosomal fraction, 5'/3' bias, median CV coverage |
| **Qualimap** | Comprehensive BAM QC and RNA-seq specific statistics |
| **MultiQC** | Aggregates all QC reports into one interactive HTML report |

##### **Quantification**

| **Tool** | **Purpose** |
| --- | --- |
| **featureCounts** (Subread) | Counts reads per gene from BAM files using GTF annotation |
| **Salmon** | Transcript quantification; supports both alignment-based and pseudo-alignment modes |

##### **Statistical Analysis (R)**

| **Package** | **Purpose** |
| --- | --- |
| **DESeq2** | Negative binomial model for differential expression; size factor normalization |
| **edgeR** | Alternative DE method; used for low-count filtering |
| **SVA** | Surrogate variable analysis for batch effect correction |
| **DRomics** | Dose-response modelling; fits sigmoidal, exponential, linear, and other models |
| **org.Hs.eg.db** | Human gene annotation database (Entrez IDs, symbols, GO terms) |
| **GO.db** | Gene Ontology database for pathway enrichment |
| **KEGGREST** | KEGG pathway database access |
| **AnnotationDbi** | Interface layer for Bioconductor annotation packages |
| **ggplot2 + ggrepel** | Publication-quality plot generation |

##### **Infrastructure**

| **Tool** | **Purpose** |
| --- | --- |
| **Nextflow** | Workflow manager; handles job scheduling, parallelism, and resumability |
| **Conda / Mamba** | Package and environment manager; keeps all tool versions isolated |
| **Streamlit** | Web framework that powers the browser interface |

#### **10. Output Files & Folders**

After a run, your output directory (~/aracra_star_work/results by default) contains:

```
results/
├── fastp/              ← Per-sample trimming stats and HTML reports
├── fastq_screen/       ← Contamination screening results
├── star/ or hisat2/    ← Aligned BAM files
├── samtools/           ← flagstat, idxstats per sample
├── rseqc/              ← Gene body coverage, strandedness, read distribution
├── picard/             ← RNA metrics
├── qualimap/           ← BAM and RNA-seq QC
├── featurecounts/      ← Raw count matrix (all samples)
├── salmon/             ← Transcript-level quantification
├── qc/
│   └── outlier/
│       ├── outlier_flag.json     ← Machine-readable QC flags
│       └── pca_qc_plot.png       ← PCA scatter for QC review
├── multiqc_report.html           ← Interactive QC summary (open in browser)
├── counts_matrix.csv             ← Gene × sample count matrix
└── runs/
    └── TreatmentName_vs_Control/
        ├── deg_results/
        │   ├── All_Results.csv               ← All genes with stats
        │   ├── Custom_Filtered_DEGs.csv      ← Significant DEGs
        │   ├── Upregulated_Genes.csv
        │   ├── Downregulated_Genes.csv
        │   ├── Final_PCA_Plot.png
        │   ├── Volcano_Plot.png
        │   └── analysis_summary.json
        └── dromics_results/
            ├── bmd_results.csv               ← Gene-level BMD estimates
            ├── top50_sensitive_genes.csv
            ├── dose_response_curves.png
            ├── bmd_distribution.png
            ├── pathway_sensitivity.png
            ├── pathway_bmd_summary.csv       ← Pathway-level tPOD
            ├── model_quality_metrics.csv
            └── dromics_summary.json

```

The **MultiQC report** (multiqc_report.html) is worth opening directly in your browser — it's an interactive dashboard with all QC metrics across all samples side by side. Use it to spot outliers, check alignment rates, and verify library quality before your manuscript.

#### **11. Troubleshooting**

##### **"⚠ .env not found — run setup.sh first"**

The sidebar shows this warning when setup hasn't been run or didn't complete. Run bash setup.sh from the ARACRA directory and wait for it to finish fully.

##### **Setup fails partway through**

The setup script uses set -euo pipefail, meaning it stops on any error. Common causes:

**No internet access:**

```
curl -I https://ftp.ebi.ac.uk
# Should return HTTP/1.1 200. If it times out, check your network/proxy.

```

**Not enough disk space:**

```
df -h ~/databases
# Need at least 80 GB free for full setup

```

**conda conflicts:** If you have an existing conda installation that's causing issues:

conda clean --all -y
bash setup.sh

##### **Pipeline starts but immediately shows FAILED**

Check the live log in the app, or read it directly:

tail -100 ~/aracra_star_work/pipeline.log

Common causes:

**SRA download failure:**

ERROR ~ process DOWNLOAD_SRA failed

→ Check your SRR accession numbers are correct and you have internet access. SRA can also be temporarily slow; try enabling "Resume" and re-running — it will pick up where it left off.

**STAR out of memory:**

STAR: Out of memory

→ Your machine has less than 32 GB RAM, or other processes are consuming RAM. Either free up memory, or switch to HISAT2 in the sidebar before running.

**Nextflow not found:**

conda activate ~/miniforge3/envs/test_ARACRA
nextflow -version

If this fails, the conda environment isn't activating. Check that run_app.sh references the correct conda path.

##### **The QC panel says "PCA plot not found"**

The PCA plot is generated during Step 1 of the DESeq2/DRomics tab (QC Check), not during Phase 1 preprocessing. Click **▶ Step 1: QC Check** in the DESeq2/DRomics tab to generate it.

##### **DESeq2 fails with "0 genes passed the filter"**

Your read count threshold is too high, or the samples have very low sequencing depth. In the Parameters tab:

- Lower the **Read count threshold** from 1,000,000 to something like 500,000 or 100,000
- Check the fastp reports (or MultiQC) to see actual read counts per sample

##### **DRomics finds 0 dose-responsive genes**

DRomics requires at least 3–4 dose levels (not counting the control) to fit dose-response curves. If you have fewer dose groups, DRomics won't have enough data points to fit a model.

Also check:

- **FDR threshold** in Parameters — try relaxing from 0.05 to 0.1
- **Transformation method** — try log or sqrt instead of auto if your data has a skewed distribution

##### **R package errors ("package 'DESeq2' not found")**

The R packages aren't loading from the conda environment. This usually means the conda environment isn't activated when the app launches.

```
# Check which R is being used
which Rscript
# Should show: ~/miniforge3/envs/test_ARACRA/bin/Rscript

# If not, activate manually and try
conda activate ~/miniforge3/envs/test_ARACRA
Rscript -e 'library(DESeq2); cat("OK\n")'

```

If DESeq2 still isn't found, reinstall it:

conda activate ~/miniforge3/envs/test_ARACRA
Rscript -e 'BiocManager::install("DESeq2", ask=FALSE)'

##### **App won't load in browser (Windows/WSL2)**

1. Make sure the terminal running run_app.sh is still open and hasn't errored
2. Try <http://127.0.0.1:8501> instead of localhost:8501
3. Check Windows Firewall isn't blocking WSL2's loopback:
   a. Open Windows Security → Firewall → Allow an app → check that your browser is allowed on private networks
4. Restart WSL: in PowerShell run wsl --shutdown, then reopen Ubuntu and relaunch the app

##### **"Resume" keeps re-running completed steps**

This can happen if the Nextflow work directory was deleted or if the .nextflow/ cache was removed. If you want a truly clean run:

rm -rf ~/aracra_star_work/work/.nextflow

Then uncheck "Resume" in the sidebar for the next run (it will redo everything from scratch).

##### **Dependency version conflicts halt the setup**

Mamba occasionally hits unsolvable conflicts when it tries to satisfy all package constraints simultaneously — especially on machines with an existing conda base that has pinned versions. You'll see something like:

```
LibMambaUnsatisfiableError: Encountered problems while solving:
  - package star-2.7.11a requires libstdc++ >=12, but none of the providers can be installed

```

or a slower variant where conda just spins for many minutes and never resolves.

**Step 1 — Try the libmamba solver explicitly:**

conda install -n base conda-libmamba-solver -y
conda config --set solver libmamba
bash setup.sh

**Step 2 — If still stuck, create the environment from scratch with a pinned Python:**

```
# Remove the broken environment
conda env remove -p ~/miniforge3/envs/test_ARACRA -y

# Re-run setup — it will rebuild cleanly
bash setup.sh

```

**Step 3 — Conflict is between two specific packages:**

Identify the conflicting packages from the error message, then install them in two passes:

```
conda activate ~/miniforge3/envs/test_ARACRA

# Install the core aligner stack first (no R)
mamba install -c conda-forge -c bioconda \
    openjdk=17 nextflow star hisat2 samtools fastp salmon subread -y

# Then install R stack separately
mamba install -c conda-forge -c bioconda \
    r-base bioconductor-deseq2 bioconductor-edger -y

```

**Step 4 — Nuclear option (completely fresh conda):**

If your existing conda base is deeply conflicted (old pinned packages, stale channels):

```
# Back up any other environments you care about first!
# Then remove miniforge/miniconda entirely
rm -rf ~/miniforge3   # or ~/miniconda3 / ~/anaconda3

# Re-run setup — it will install a fresh Miniforge3 automatically
bash setup.sh

```

**Step 5 — Salmon GLIBC incompatibility (separate issue):**

On some older Linux systems or certain WSL2 configurations, Salmon installs but fails to run:

salmon: /lib/x86_64-linux-gnu/libc.so.6: version GLIBC_2.34 not found

Fix:

conda activate ~/miniforge3/envs/test_ARACRA
mamba install -c bioconda 'salmon>=1.10' --force-reinstall -y

If that doesn't resolve it, run setup with --skip-index to at least get the rest of the pipeline working, then use featureCounts for quantification instead of Salmon.

##### **Getting more help**

- Check the live log first (~/aracra_star_work/pipeline.log) — it usually contains the exact error
- Use the **▶ Check Tools** button in the sidebar to confirm all tools are installed
- For Nextflow-specific issues: the Nextflow log is at .nextflow.log in the ARACRA directory
- For any issue, create a ticket in issues section of GitHub.

*ARACRA Pipeline · v2.0 · For research use*
