## Supplementary material for "ARACRA: Automated RNA-seq Analysis for Chemical Risk Assessment": HTML file: S3_multiqc.html

MultiQC Report


### Toggle navigation v1.25.1

Loading report..

- General Stats
- Samtools
  - Flagstat
  - Mapped reads per contig
- STAR
  - Summary Statistics
  - Alignment Scores
- FastQ Screen
- fastp
  - Filtered Reads
  - Sequence Quality
  - GC Content
  - N content
- Software Versions

Toolbox

##### MultiQC Toolbox

###### Apply Highlight Samples

+

Regex mode off
help
 Clear

###### Apply Rename Samples

+

Click here for bulk input.

Paste two columns of a tab-delimited table here (eg. from Excel).

First column should be the old name, second column the new name.

Add

Regex mode off
help
 Clear

###### Apply Show / Hide Samples

Hide matching samples

Show only matching samples

+

Regex mode off
help
 Clear

###### Export Plots

- Images
- Data

px

px

Aspect ratio

PNG
SVG

Plot scaling

X

Download the raw data used to create the plots in this report below:

Format:

Tab-separated
Comma-separated
JSON

Note that additional data was saved in `multiqc_data` when this report was generated.

---

###### Choose Plots

 All
 None


---


   Download Plot Images

If you use plots from MultiQC in a publication or presentation, please cite:

> **MultiQC: Summarize analysis results for multiple tools and samples in a single report**  
> *Philip Ewels, Måns Magnusson, Sverker Lundin and Max Käller*  
> Bioinformatics (2016)  
> doi: 10.1093/bioinformatics/btw354  
> PMID: 27312411

###### Save Settings

You can save the toolbox settings for this report to the browser.

 Save


---

###### Load Settings

Choose a saved report profile from the dropdown box below:

[ select ]

Load
 Delete
 Set default
 Clear default

###### Tool Citations

Please remember to cite the tools that you use in your analysis.

To help with this, you can download publication details of the tools mentioned in this report:

List of DOIs

BibTeX file

###### About MultiQC

This report was generated using MultiQC, version 1.25.1

You can see a YouTube video describing how to use MultiQC reports here:
https://youtu.be/qPbIlO\_KWN0

For more information about MultiQC, including other videos and
extensive documentation, please visit http://multiqc.info

You can report bugs, suggest improvements and find the source code for MultiQC on GitHub:
https://github.com/MultiQC/MultiQC

MultiQC is published in Bioinformatics:

> **MultiQC: Summarize analysis results for multiple tools and samples in a single report**  
> *Philip Ewels, Måns Magnusson, Sverker Lundin and Max Käller*  
> Bioinformatics (2016)  
> doi: 10.1093/bioinformatics/btw354  
> PMID: 27312411

---

MultiQC is developed by:

A modular tool to aggregate results from bioinformatics analyses across many samples into a single report.

###### JavaScript Disabled

MultiQC reports use JavaScript for plots and toolbox functions. It looks like
you have JavaScript disabled in your web browser. Please note that many of the report
functions will not work as intended.

Loading report..

Report
generated on 2026-03-18, 21:27 CET
based on data in:
`/home/saurav/aracra_star_work/work/b9/1846559dfa4969250c1040daba3ce3`

---

×
don't show again

**Welcome!** Not sure where to start?  
Watch a tutorial video
  *(6:06)*

×

Because this report contains a lot of samples, you may need to click 'Show plot' to see some graphs.
Render all plots

#### General Statistics

Showing 572 samples.

Configure columns
Export Plot

Created with MultiQC

#### Samtools

Toolkit for interacting with BAM/CRAM files.*URL: http://www.htslib.org**DOI: 10.1093/bioinformatics/btp352*

##### Flagstat

This module parses the output from `samtools flagstat`

Configure columns
 Table

Read counts
Percentage of total

Export Plot

Created with MultiQC

Copy table

 Configure columns

 Sort by highlight

 Scatter plot

 Violin plot
Export as CSV
Showing ^0^/_286_ rows and ^10^/_10_ columns.

| Sample Name | Total Reads | Total Passed QC | Mapped | Duplicates | Paired in Sequencing | Properly Paired | Self and mate mapped | Singletons | Mate mapped to diff chr | Diff chr (mapQ >= 5) |
| --- | --- | --- | --- | --- | --- | --- | --- | --- | --- | --- |
| SRR29684827\_flagstat | 3.9M | 3.9M | 3.9M | 0.0M | 0.0M | 0.0M | 0.0M | 0.0M | 0.0M | 0.0M |
| SRR29684828\_flagstat | 0.1M | 0.1M | 0.1M | 0.0M | 0.0M | 0.0M | 0.0M | 0.0M | 0.0M | 0.0M |
| SRR29684829\_flagstat | 3.7M | 3.7M | 3.7M | 0.0M | 0.0M | 0.0M | 0.0M | 0.0M | 0.0M | 0.0M |
| SRR29684830\_flagstat | 3.4M | 3.4M | 3.4M | 0.0M | 0.0M | 0.0M | 0.0M | 0.0M | 0.0M | 0.0M |
| SRR29684831\_flagstat | 4.3M | 4.3M | 4.3M | 0.0M | 0.0M | 0.0M | 0.0M | 0.0M | 0.0M | 0.0M |
| SRR29684832\_flagstat | 4.4M | 4.4M | 4.4M | 0.0M | 0.0M | 0.0M | 0.0M | 0.0M | 0.0M | 0.0M |
| SRR29684833\_flagstat | 4.4M | 4.4M | 4.4M | 0.0M | 0.0M | 0.0M | 0.0M | 0.0M | 0.0M | 0.0M |
| SRR29684834\_flagstat | 3.7M | 3.7M | 3.7M | 0.0M | 0.0M | 0.0M | 0.0M | 0.0M | 0.0M | 0.0M |
| SRR29684835\_flagstat | 3.7M | 3.7M | 3.7M | 0.0M | 0.0M | 0.0M | 0.0M | 0.0M | 0.0M | 0.0M |
| SRR29684836\_flagstat | 4.0M | 4.0M | 4.0M | 0.0M | 0.0M | 0.0M | 0.0M | 0.0M | 0.0M | 0.0M |
| SRR29684837\_flagstat | 4.8M | 4.8M | 4.8M | 0.0M | 0.0M | 0.0M | 0.0M | 0.0M | 0.0M | 0.0M |
| SRR29684838\_flagstat | 3.9M | 3.9M | 3.9M | 0.0M | 0.0M | 0.0M | 0.0M | 0.0M | 0.0M | 0.0M |
| SRR29684839\_flagstat | 4.6M | 4.6M | 4.6M | 0.0M | 0.0M | 0.0M | 0.0M | 0.0M | 0.0M | 0.0M |
| SRR29684840\_flagstat | 4.1M | 4.1M | 4.1M | 0.0M | 0.0M | 0.0M | 0.0M | 0.0M | 0.0M | 0.0M |
| SRR29684841\_flagstat | 3.5M | 3.5M | 3.5M | 0.0M | 0.0M | 0.0M | 0.0M | 0.0M | 0.0M | 0.0M |
| SRR29684842\_flagstat | 3.0M | 3.0M | 3.0M | 0.0M | 0.0M | 0.0M | 0.0M | 0.0M | 0.0M | 0.0M |
| SRR29684843\_flagstat | 4.3M | 4.3M | 4.3M | 0.0M | 0.0M | 0.0M | 0.0M | 0.0M | 0.0M | 0.0M |
| SRR29684844\_flagstat | 5.7M | 5.7M | 5.7M | 0.0M | 0.0M | 0.0M | 0.0M | 0.0M | 0.0M | 0.0M |
| SRR29684845\_flagstat | 6.4M | 6.4M | 6.4M | 0.0M | 0.0M | 0.0M | 0.0M | 0.0M | 0.0M | 0.0M |
| SRR29684846\_flagstat | 4.6M | 4.6M | 4.6M | 0.0M | 0.0M | 0.0M | 0.0M | 0.0M | 0.0M | 0.0M |
| SRR29684847\_flagstat | 6.4M | 6.4M | 6.4M | 0.0M | 0.0M | 0.0M | 0.0M | 0.0M | 0.0M | 0.0M |
| SRR29684848\_flagstat | 5.6M | 5.6M | 5.6M | 0.0M | 0.0M | 0.0M | 0.0M | 0.0M | 0.0M | 0.0M |
| SRR29684849\_flagstat | 4.6M | 4.6M | 4.6M | 0.0M | 0.0M | 0.0M | 0.0M | 0.0M | 0.0M | 0.0M |
| SRR29684850\_flagstat | 3.5M | 3.5M | 3.5M | 0.0M | 0.0M | 0.0M | 0.0M | 0.0M | 0.0M | 0.0M |
| SRR29684851\_flagstat | 1.8M | 1.8M | 1.8M | 0.0M | 0.0M | 0.0M | 0.0M | 0.0M | 0.0M | 0.0M |
| SRR29684852\_flagstat | 1.4M | 1.4M | 1.4M | 0.0M | 0.0M | 0.0M | 0.0M | 0.0M | 0.0M | 0.0M |
| SRR29684853\_flagstat | 2.6M | 2.6M | 2.6M | 0.0M | 0.0M | 0.0M | 0.0M | 0.0M | 0.0M | 0.0M |
| SRR29684854\_flagstat | 2.6M | 2.6M | 2.6M | 0.0M | 0.0M | 0.0M | 0.0M | 0.0M | 0.0M | 0.0M |
| SRR29684855\_flagstat | 1.3M | 1.3M | 1.3M | 0.0M | 0.0M | 0.0M | 0.0M | 0.0M | 0.0M | 0.0M |
| SRR29684856\_flagstat | 0.4M | 0.4M | 0.4M | 0.0M | 0.0M | 0.0M | 0.0M | 0.0M | 0.0M | 0.0M |
| SRR29684857\_flagstat | 0.8M | 0.8M | 0.8M | 0.0M | 0.0M | 0.0M | 0.0M | 0.0M | 0.0M | 0.0M |
| SRR29684858\_flagstat | 1.5M | 1.5M | 1.5M | 0.0M | 0.0M | 0.0M | 0.0M | 0.0M | 0.0M | 0.0M |
| SRR29684859\_flagstat | 2.2M | 2.2M | 2.2M | 0.0M | 0.0M | 0.0M | 0.0M | 0.0M | 0.0M | 0.0M |
| SRR29684860\_flagstat | 2.4M | 2.4M | 2.4M | 0.0M | 0.0M | 0.0M | 0.0M | 0.0M | 0.0M | 0.0M |
| SRR29684861\_flagstat | 2.4M | 2.4M | 2.4M | 0.0M | 0.0M | 0.0M | 0.0M | 0.0M | 0.0M | 0.0M |
| SRR29684862\_flagstat | 1.6M | 1.6M | 1.6M | 0.0M | 0.0M | 0.0M | 0.0M | 0.0M | 0.0M | 0.0M |
| SRR29684863\_flagstat | 2.2M | 2.2M | 2.2M | 0.0M | 0.0M | 0.0M | 0.0M | 0.0M | 0.0M | 0.0M |
| SRR29684864\_flagstat | 1.8M | 1.8M | 1.8M | 0.0M | 0.0M | 0.0M | 0.0M | 0.0M | 0.0M | 0.0M |
| SRR29684865\_flagstat | 1.6M | 1.6M | 1.6M | 0.0M | 0.0M | 0.0M | 0.0M | 0.0M | 0.0M | 0.0M |
| SRR29684866\_flagstat | 1.4M | 1.4M | 1.4M | 0.0M | 0.0M | 0.0M | 0.0M | 0.0M | 0.0M | 0.0M |
| SRR29684867\_flagstat | 1.7M | 1.7M | 1.7M | 0.0M | 0.0M | 0.0M | 0.0M | 0.0M | 0.0M | 0.0M |
| SRR29684868\_flagstat | 3.2M | 3.2M | 3.2M | 0.0M | 0.0M | 0.0M | 0.0M | 0.0M | 0.0M | 0.0M |
| SRR29684869\_flagstat | 2.8M | 2.8M | 2.8M | 0.0M | 0.0M | 0.0M | 0.0M | 0.0M | 0.0M | 0.0M |
| SRR29684870\_flagstat | 2.3M | 2.3M | 2.3M | 0.0M | 0.0M | 0.0M | 0.0M | 0.0M | 0.0M | 0.0M |
| SRR29684871\_flagstat | 3.3M | 3.3M | 3.3M | 0.0M | 0.0M | 0.0M | 0.0M | 0.0M | 0.0M | 0.0M |
| SRR29684872\_flagstat | 3.3M | 3.3M | 3.3M | 0.0M | 0.0M | 0.0M | 0.0M | 0.0M | 0.0M | 0.0M |
| SRR29684873\_flagstat | 3.3M | 3.3M | 3.3M | 0.0M | 0.0M | 0.0M | 0.0M | 0.0M | 0.0M | 0.0M |
| SRR29684874\_flagstat | 0.7M | 0.7M | 0.7M | 0.0M | 0.0M | 0.0M | 0.0M | 0.0M | 0.0M | 0.0M |
| SRR29684875\_flagstat | 2.6M | 2.6M | 2.6M | 0.0M | 0.0M | 0.0M | 0.0M | 0.0M | 0.0M | 0.0M |
| SRR29684876\_flagstat | 4.1M | 4.1M | 4.1M | 0.0M | 0.0M | 0.0M | 0.0M | 0.0M | 0.0M | 0.0M |
| SRR29684877\_flagstat | 3.8M | 3.8M | 3.8M | 0.0M | 0.0M | 0.0M | 0.0M | 0.0M | 0.0M | 0.0M |
| SRR29684878\_flagstat | 5.1M | 5.1M | 5.1M | 0.0M | 0.0M | 0.0M | 0.0M | 0.0M | 0.0M | 0.0M |
| SRR29684879\_flagstat | 4.4M | 4.4M | 4.4M | 0.0M | 0.0M | 0.0M | 0.0M | 0.0M | 0.0M | 0.0M |
| SRR29684880\_flagstat | 4.8M | 4.8M | 4.8M | 0.0M | 0.0M | 0.0M | 0.0M | 0.0M | 0.0M | 0.0M |
| SRR29684881\_flagstat | 4.3M | 4.3M | 4.3M | 0.0M | 0.0M | 0.0M | 0.0M | 0.0M | 0.0M | 0.0M |
| SRR29684882\_flagstat | 4.2M | 4.2M | 4.2M | 0.0M | 0.0M | 0.0M | 0.0M | 0.0M | 0.0M | 0.0M |
| SRR29684883\_flagstat | 3.1M | 3.1M | 3.1M | 0.0M | 0.0M | 0.0M | 0.0M | 0.0M | 0.0M | 0.0M |
| SRR29684884\_flagstat | 3.3M | 3.3M | 3.3M | 0.0M | 0.0M | 0.0M | 0.0M | 0.0M | 0.0M | 0.0M |
| SRR29684885\_flagstat | 2.2M | 2.2M | 2.2M | 0.0M | 0.0M | 0.0M | 0.0M | 0.0M | 0.0M | 0.0M |
| SRR29684886\_flagstat | 5.0M | 5.0M | 5.0M | 0.0M | 0.0M | 0.0M | 0.0M | 0.0M | 0.0M | 0.0M |
| SRR29684887\_flagstat | 3.7M | 3.7M | 3.7M | 0.0M | 0.0M | 0.0M | 0.0M | 0.0M | 0.0M | 0.0M |
| SRR29684888\_flagstat | 4.2M | 4.2M | 4.2M | 0.0M | 0.0M | 0.0M | 0.0M | 0.0M | 0.0M | 0.0M |
| SRR29684889\_flagstat | 4.2M | 4.2M | 4.2M | 0.0M | 0.0M | 0.0M | 0.0M | 0.0M | 0.0M | 0.0M |
| SRR29684890\_flagstat | 3.1M | 3.1M | 3.1M | 0.0M | 0.0M | 0.0M | 0.0M | 0.0M | 0.0M | 0.0M |
| SRR29684891\_flagstat | 2.7M | 2.7M | 2.7M | 0.0M | 0.0M | 0.0M | 0.0M | 0.0M | 0.0M | 0.0M |
| SRR29684892\_flagstat | 3.5M | 3.5M | 3.5M | 0.0M | 0.0M | 0.0M | 0.0M | 0.0M | 0.0M | 0.0M |
| SRR29684893\_flagstat | 4.2M | 4.2M | 4.2M | 0.0M | 0.0M | 0.0M | 0.0M | 0.0M | 0.0M | 0.0M |
| SRR29684894\_flagstat | 6.4M | 6.4M | 6.4M | 0.0M | 0.0M | 0.0M | 0.0M | 0.0M | 0.0M | 0.0M |
| SRR29684895\_flagstat | 4.7M | 4.7M | 4.7M | 0.0M | 0.0M | 0.0M | 0.0M | 0.0M | 0.0M | 0.0M |
| SRR29684896\_flagstat | 5.0M | 5.0M | 5.0M | 0.0M | 0.0M | 0.0M | 0.0M | 0.0M | 0.0M | 0.0M |
| SRR29684897\_flagstat | 4.5M | 4.5M | 4.5M | 0.0M | 0.0M | 0.0M | 0.0M | 0.0M | 0.0M | 0.0M |
| SRR29684898\_flagstat | 4.8M | 4.8M | 4.8M | 0.0M | 0.0M | 0.0M | 0.0M | 0.0M | 0.0M | 0.0M |
| SRR29684899\_flagstat | 3.0M | 3.0M | 3.0M | 0.0M | 0.0M | 0.0M | 0.0M | 0.0M | 0.0M | 0.0M |
| SRR29684900\_flagstat | 3.4M | 3.4M | 3.4M | 0.0M | 0.0M | 0.0M | 0.0M | 0.0M | 0.0M | 0.0M |
| SRR29684901\_flagstat | 4.7M | 4.7M | 4.7M | 0.0M | 0.0M | 0.0M | 0.0M | 0.0M | 0.0M | 0.0M |
| SRR29684902\_flagstat | 4.2M | 4.2M | 4.2M | 0.0M | 0.0M | 0.0M | 0.0M | 0.0M | 0.0M | 0.0M |
| SRR29684903\_flagstat | 4.7M | 4.7M | 4.7M | 0.0M | 0.0M | 0.0M | 0.0M | 0.0M | 0.0M | 0.0M |
| SRR29684904\_flagstat | 4.0M | 4.0M | 4.0M | 0.0M | 0.0M | 0.0M | 0.0M | 0.0M | 0.0M | 0.0M |
| SRR29684905\_flagstat | 1.0M | 1.0M | 1.0M | 0.0M | 0.0M | 0.0M | 0.0M | 0.0M | 0.0M | 0.0M |
| SRR29684906\_flagstat | 3.3M | 3.3M | 3.3M | 0.0M | 0.0M | 0.0M | 0.0M | 0.0M | 0.0M | 0.0M |
| SRR29684907\_flagstat | 3.9M | 3.9M | 3.9M | 0.0M | 0.0M | 0.0M | 0.0M | 0.0M | 0.0M | 0.0M |
| SRR29684908\_flagstat | 1.9M | 1.9M | 1.9M | 0.0M | 0.0M | 0.0M | 0.0M | 0.0M | 0.0M | 0.0M |
| SRR29684909\_flagstat | 5.0M | 5.0M | 5.0M | 0.0M | 0.0M | 0.0M | 0.0M | 0.0M | 0.0M | 0.0M |
| SRR29684910\_flagstat | 3.6M | 3.6M | 3.6M | 0.0M | 0.0M | 0.0M | 0.0M | 0.0M | 0.0M | 0.0M |
| SRR29684911\_flagstat | 3.8M | 3.8M | 3.8M | 0.0M | 0.0M | 0.0M | 0.0M | 0.0M | 0.0M | 0.0M |
| SRR29684912\_flagstat | 4.4M | 4.4M | 4.4M | 0.0M | 0.0M | 0.0M | 0.0M | 0.0M | 0.0M | 0.0M |
| SRR29684913\_flagstat | 4.5M | 4.5M | 4.5M | 0.0M | 0.0M | 0.0M | 0.0M | 0.0M | 0.0M | 0.0M |
| SRR29684914\_flagstat | 1.0M | 1.0M | 1.0M | 0.0M | 0.0M | 0.0M | 0.0M | 0.0M | 0.0M | 0.0M |
| SRR29684915\_flagstat | 1.2M | 1.2M | 1.2M | 0.0M | 0.0M | 0.0M | 0.0M | 0.0M | 0.0M | 0.0M |
| SRR29684916\_flagstat | 1.5M | 1.5M | 1.5M | 0.0M | 0.0M | 0.0M | 0.0M | 0.0M | 0.0M | 0.0M |
| SRR29684917\_flagstat | 2.6M | 2.6M | 2.6M | 0.0M | 0.0M | 0.0M | 0.0M | 0.0M | 0.0M | 0.0M |
| SRR29684918\_flagstat | 0.0M | 0.0M | 0.0M | 0.0M | 0.0M | 0.0M | 0.0M | 0.0M | 0.0M | 0.0M |
| SRR29684919\_flagstat | 2.5M | 2.5M | 2.5M | 0.0M | 0.0M | 0.0M | 0.0M | 0.0M | 0.0M | 0.0M |
| SRR29684920\_flagstat | 2.0M | 2.0M | 2.0M | 0.0M | 0.0M | 0.0M | 0.0M | 0.0M | 0.0M | 0.0M |
| SRR29684921\_flagstat | 2.1M | 2.1M | 2.1M | 0.0M | 0.0M | 0.0M | 0.0M | 0.0M | 0.0M | 0.0M |
| SRR29684922\_flagstat | 1.4M | 1.4M | 1.4M | 0.0M | 0.0M | 0.0M | 0.0M | 0.0M | 0.0M | 0.0M |
| SRR29684923\_flagstat | 1.9M | 1.9M | 1.9M | 0.0M | 0.0M | 0.0M | 0.0M | 0.0M | 0.0M | 0.0M |
| SRR29684924\_flagstat | 2.1M | 2.1M | 2.1M | 0.0M | 0.0M | 0.0M | 0.0M | 0.0M | 0.0M | 0.0M |
| SRR29684925\_flagstat | 1.1M | 1.1M | 1.1M | 0.0M | 0.0M | 0.0M | 0.0M | 0.0M | 0.0M | 0.0M |
| SRR29684926\_flagstat | 3.0M | 3.0M | 3.0M | 0.0M | 0.0M | 0.0M | 0.0M | 0.0M | 0.0M | 0.0M |
| SRR29684927\_flagstat | 2.5M | 2.5M | 2.5M | 0.0M | 0.0M | 0.0M | 0.0M | 0.0M | 0.0M | 0.0M |
| SRR29684928\_flagstat | 2.6M | 2.6M | 2.6M | 0.0M | 0.0M | 0.0M | 0.0M | 0.0M | 0.0M | 0.0M |
| SRR29684929\_flagstat | 2.5M | 2.5M | 2.5M | 0.0M | 0.0M | 0.0M | 0.0M | 0.0M | 0.0M | 0.0M |
| SRR29684930\_flagstat | 2.9M | 2.9M | 2.9M | 0.0M | 0.0M | 0.0M | 0.0M | 0.0M | 0.0M | 0.0M |
| SRR29684931\_flagstat | 3.2M | 3.2M | 3.2M | 0.0M | 0.0M | 0.0M | 0.0M | 0.0M | 0.0M | 0.0M |
| SRR29684932\_flagstat | 3.0M | 3.0M | 3.0M | 0.0M | 0.0M | 0.0M | 0.0M | 0.0M | 0.0M | 0.0M |
| SRR29684933\_flagstat | 2.0M | 2.0M | 2.0M | 0.0M | 0.0M | 0.0M | 0.0M | 0.0M | 0.0M | 0.0M |
| SRR29684934\_flagstat | 2.7M | 2.7M | 2.7M | 0.0M | 0.0M | 0.0M | 0.0M | 0.0M | 0.0M | 0.0M |
| SRR29684935\_flagstat | 2.3M | 2.3M | 2.3M | 0.0M | 0.0M | 0.0M | 0.0M | 0.0M | 0.0M | 0.0M |
| SRR29684936\_flagstat | 2.0M | 2.0M | 2.0M | 0.0M | 0.0M | 0.0M | 0.0M | 0.0M | 0.0M | 0.0M |
| SRR29684937\_flagstat | 2.5M | 2.5M | 2.5M | 0.0M | 0.0M | 0.0M | 0.0M | 0.0M | 0.0M | 0.0M |
| SRR29684938\_flagstat | 3.8M | 3.8M | 3.8M | 0.0M | 0.0M | 0.0M | 0.0M | 0.0M | 0.0M | 0.0M |
| SRR29684939\_flagstat | 3.8M | 3.8M | 3.8M | 0.0M | 0.0M | 0.0M | 0.0M | 0.0M | 0.0M | 0.0M |
| SRR29684940\_flagstat | 3.3M | 3.3M | 3.3M | 0.0M | 0.0M | 0.0M | 0.0M | 0.0M | 0.0M | 0.0M |
| SRR29684941\_flagstat | 3.1M | 3.1M | 3.1M | 0.0M | 0.0M | 0.0M | 0.0M | 0.0M | 0.0M | 0.0M |
| SRR29684942\_flagstat | 4.8M | 4.8M | 4.8M | 0.0M | 0.0M | 0.0M | 0.0M | 0.0M | 0.0M | 0.0M |
| SRR29684943\_flagstat | 0.8M | 0.8M | 0.8M | 0.0M | 0.0M | 0.0M | 0.0M | 0.0M | 0.0M | 0.0M |
| SRR29684944\_flagstat | 4.6M | 4.6M | 4.6M | 0.0M | 0.0M | 0.0M | 0.0M | 0.0M | 0.0M | 0.0M |
| SRR29684945\_flagstat | 3.9M | 3.9M | 3.9M | 0.0M | 0.0M | 0.0M | 0.0M | 0.0M | 0.0M | 0.0M |
| SRR29684946\_flagstat | 4.1M | 4.1M | 4.1M | 0.0M | 0.0M | 0.0M | 0.0M | 0.0M | 0.0M | 0.0M |
| SRR29684947\_flagstat | 3.0M | 3.0M | 3.0M | 0.0M | 0.0M | 0.0M | 0.0M | 0.0M | 0.0M | 0.0M |
| SRR29684948\_flagstat | 4.8M | 4.8M | 4.8M | 0.0M | 0.0M | 0.0M | 0.0M | 0.0M | 0.0M | 0.0M |
| SRR29684949\_flagstat | 5.2M | 5.2M | 5.2M | 0.0M | 0.0M | 0.0M | 0.0M | 0.0M | 0.0M | 0.0M |
| SRR29684950\_flagstat | 7.1M | 7.1M | 7.1M | 0.0M | 0.0M | 0.0M | 0.0M | 0.0M | 0.0M | 0.0M |
| SRR29684951\_flagstat | 5.6M | 5.6M | 5.6M | 0.0M | 0.0M | 0.0M | 0.0M | 0.0M | 0.0M | 0.0M |
| SRR29684952\_flagstat | 6.4M | 6.4M | 6.4M | 0.0M | 0.0M | 0.0M | 0.0M | 0.0M | 0.0M | 0.0M |
| SRR29684953\_flagstat | 6.3M | 6.3M | 6.3M | 0.0M | 0.0M | 0.0M | 0.0M | 0.0M | 0.0M | 0.0M |
| SRR29684954\_flagstat | 5.8M | 5.8M | 5.8M | 0.0M | 0.0M | 0.0M | 0.0M | 0.0M | 0.0M | 0.0M |
| SRR29684955\_flagstat | 3.9M | 3.9M | 3.9M | 0.0M | 0.0M | 0.0M | 0.0M | 0.0M | 0.0M | 0.0M |
| SRR29684956\_flagstat | 2.9M | 2.9M | 2.9M | 0.0M | 0.0M | 0.0M | 0.0M | 0.0M | 0.0M | 0.0M |
| SRR29684957\_flagstat | 2.5M | 2.5M | 2.5M | 0.0M | 0.0M | 0.0M | 0.0M | 0.0M | 0.0M | 0.0M |
| SRR29684958\_flagstat | 2.6M | 2.6M | 2.6M | 0.0M | 0.0M | 0.0M | 0.0M | 0.0M | 0.0M | 0.0M |
| SRR29684959\_flagstat | 1.1M | 1.1M | 1.1M | 0.0M | 0.0M | 0.0M | 0.0M | 0.0M | 0.0M | 0.0M |
| SRR29684960\_flagstat | 0.8M | 0.8M | 0.8M | 0.0M | 0.0M | 0.0M | 0.0M | 0.0M | 0.0M | 0.0M |
| SRR29684961\_flagstat | 0.7M | 0.7M | 0.7M | 0.0M | 0.0M | 0.0M | 0.0M | 0.0M | 0.0M | 0.0M |
| SRR29684962\_flagstat | 2.5M | 2.5M | 2.5M | 0.0M | 0.0M | 0.0M | 0.0M | 0.0M | 0.0M | 0.0M |
| SRR29684963\_flagstat | 2.4M | 2.4M | 2.4M | 0.0M | 0.0M | 0.0M | 0.0M | 0.0M | 0.0M | 0.0M |
| SRR29684964\_flagstat | 2.5M | 2.5M | 2.5M | 0.0M | 0.0M | 0.0M | 0.0M | 0.0M | 0.0M | 0.0M |
| SRR29684965\_flagstat | 2.3M | 2.3M | 2.3M | 0.0M | 0.0M | 0.0M | 0.0M | 0.0M | 0.0M | 0.0M |
| SRR29684966\_flagstat | 1.4M | 1.4M | 1.4M | 0.0M | 0.0M | 0.0M | 0.0M | 0.0M | 0.0M | 0.0M |
| SRR29684967\_flagstat | 2.2M | 2.2M | 2.2M | 0.0M | 0.0M | 0.0M | 0.0M | 0.0M | 0.0M | 0.0M |
| SRR29684968\_flagstat | 1.7M | 1.7M | 1.7M | 0.0M | 0.0M | 0.0M | 0.0M | 0.0M | 0.0M | 0.0M |
| SRR29684969\_flagstat | 1.7M | 1.7M | 1.7M | 0.0M | 0.0M | 0.0M | 0.0M | 0.0M | 0.0M | 0.0M |
| SRR29684970\_flagstat | 1.1M | 1.1M | 1.1M | 0.0M | 0.0M | 0.0M | 0.0M | 0.0M | 0.0M | 0.0M |
| SRR29684971\_flagstat | 3.5M | 3.5M | 3.5M | 0.0M | 0.0M | 0.0M | 0.0M | 0.0M | 0.0M | 0.0M |
| SRR29684972\_flagstat | 3.3M | 3.3M | 3.3M | 0.0M | 0.0M | 0.0M | 0.0M | 0.0M | 0.0M | 0.0M |
| SRR29684973\_flagstat | 2.2M | 2.2M | 2.2M | 0.0M | 0.0M | 0.0M | 0.0M | 0.0M | 0.0M | 0.0M |
| SRR29684974\_flagstat | 4.3M | 4.3M | 4.3M | 0.0M | 0.0M | 0.0M | 0.0M | 0.0M | 0.0M | 0.0M |
| SRR29684975\_flagstat | 3.3M | 3.3M | 3.3M | 0.0M | 0.0M | 0.0M | 0.0M | 0.0M | 0.0M | 0.0M |
| SRR29684976\_flagstat | 2.1M | 2.1M | 2.1M | 0.0M | 0.0M | 0.0M | 0.0M | 0.0M | 0.0M | 0.0M |
| SRR29684977\_flagstat | 3.5M | 3.5M | 3.5M | 0.0M | 0.0M | 0.0M | 0.0M | 0.0M | 0.0M | 0.0M |
| SRR29684978\_flagstat | 1.9M | 1.9M | 1.9M | 0.0M | 0.0M | 0.0M | 0.0M | 0.0M | 0.0M | 0.0M |
| SRR29684979\_flagstat | 3.5M | 3.5M | 3.5M | 0.0M | 0.0M | 0.0M | 0.0M | 0.0M | 0.0M | 0.0M |
| SRR29684980\_flagstat | 3.1M | 3.1M | 3.1M | 0.0M | 0.0M | 0.0M | 0.0M | 0.0M | 0.0M | 0.0M |
| SRR29684981\_flagstat | 3.3M | 3.3M | 3.3M | 0.0M | 0.0M | 0.0M | 0.0M | 0.0M | 0.0M | 0.0M |
| SRR29684982\_flagstat | 3.5M | 3.5M | 3.5M | 0.0M | 0.0M | 0.0M | 0.0M | 0.0M | 0.0M | 0.0M |
| SRR29684983\_flagstat | 3.3M | 3.3M | 3.3M | 0.0M | 0.0M | 0.0M | 0.0M | 0.0M | 0.0M | 0.0M |
| SRR29684984\_flagstat | 3.4M | 3.4M | 3.4M | 0.0M | 0.0M | 0.0M | 0.0M | 0.0M | 0.0M | 0.0M |
| SRR29684985\_flagstat | 3.7M | 3.7M | 3.7M | 0.0M | 0.0M | 0.0M | 0.0M | 0.0M | 0.0M | 0.0M |
| SRR29684986\_flagstat | 3.4M | 3.4M | 3.4M | 0.0M | 0.0M | 0.0M | 0.0M | 0.0M | 0.0M | 0.0M |
| SRR29684987\_flagstat | 3.9M | 3.9M | 3.9M | 0.0M | 0.0M | 0.0M | 0.0M | 0.0M | 0.0M | 0.0M |
| SRR29684988\_flagstat | 2.6M | 2.6M | 2.6M | 0.0M | 0.0M | 0.0M | 0.0M | 0.0M | 0.0M | 0.0M |
| SRR29684989\_flagstat | 5.7M | 5.7M | 5.7M | 0.0M | 0.0M | 0.0M | 0.0M | 0.0M | 0.0M | 0.0M |
| SRR29684990\_flagstat | 4.6M | 4.6M | 4.6M | 0.0M | 0.0M | 0.0M | 0.0M | 0.0M | 0.0M | 0.0M |
| SRR29684991\_flagstat | 3.9M | 3.9M | 3.9M | 0.0M | 0.0M | 0.0M | 0.0M | 0.0M | 0.0M | 0.0M |
| SRR29684992\_flagstat | 4.8M | 4.8M | 4.8M | 0.0M | 0.0M | 0.0M | 0.0M | 0.0M | 0.0M | 0.0M |
| SRR29684993\_flagstat | 4.8M | 4.8M | 4.8M | 0.0M | 0.0M | 0.0M | 0.0M | 0.0M | 0.0M | 0.0M |
| SRR29684994\_flagstat | 3.2M | 3.2M | 3.2M | 0.0M | 0.0M | 0.0M | 0.0M | 0.0M | 0.0M | 0.0M |
| SRR29684995\_flagstat | 3.7M | 3.7M | 3.7M | 0.0M | 0.0M | 0.0M | 0.0M | 0.0M | 0.0M | 0.0M |
| SRR29684996\_flagstat | 3.9M | 3.9M | 3.9M | 0.0M | 0.0M | 0.0M | 0.0M | 0.0M | 0.0M | 0.0M |
| SRR29684997\_flagstat | 5.2M | 5.2M | 5.2M | 0.0M | 0.0M | 0.0M | 0.0M | 0.0M | 0.0M | 0.0M |
| SRR29684998\_flagstat | 3.8M | 3.8M | 3.8M | 0.0M | 0.0M | 0.0M | 0.0M | 0.0M | 0.0M | 0.0M |
| SRR29684999\_flagstat | 4.6M | 4.6M | 4.6M | 0.0M | 0.0M | 0.0M | 0.0M | 0.0M | 0.0M | 0.0M |
| SRR29685000\_flagstat | 4.2M | 4.2M | 4.2M | 0.0M | 0.0M | 0.0M | 0.0M | 0.0M | 0.0M | 0.0M |
| SRR29685001\_flagstat | 4.0M | 4.0M | 4.0M | 0.0M | 0.0M | 0.0M | 0.0M | 0.0M | 0.0M | 0.0M |
| SRR29685002\_flagstat | 3.9M | 3.9M | 3.9M | 0.0M | 0.0M | 0.0M | 0.0M | 0.0M | 0.0M | 0.0M |
| SRR29685003\_flagstat | 4.0M | 4.0M | 4.0M | 0.0M | 0.0M | 0.0M | 0.0M | 0.0M | 0.0M | 0.0M |
| SRR29685004\_flagstat | 5.7M | 5.7M | 5.7M | 0.0M | 0.0M | 0.0M | 0.0M | 0.0M | 0.0M | 0.0M |
| SRR29685005\_flagstat | 2.7M | 2.7M | 2.7M | 0.0M | 0.0M | 0.0M | 0.0M | 0.0M | 0.0M | 0.0M |
| SRR29685006\_flagstat | 4.7M | 4.7M | 4.7M | 0.0M | 0.0M | 0.0M | 0.0M | 0.0M | 0.0M | 0.0M |
| SRR29685007\_flagstat | 4.6M | 4.6M | 4.6M | 0.0M | 0.0M | 0.0M | 0.0M | 0.0M | 0.0M | 0.0M |
| SRR29685008\_flagstat | 4.4M | 4.4M | 4.4M | 0.0M | 0.0M | 0.0M | 0.0M | 0.0M | 0.0M | 0.0M |
| SRR29685009\_flagstat | 3.5M | 3.5M | 3.5M | 0.0M | 0.0M | 0.0M | 0.0M | 0.0M | 0.0M | 0.0M |
| SRR29685010\_flagstat | 3.2M | 3.2M | 3.2M | 0.0M | 0.0M | 0.0M | 0.0M | 0.0M | 0.0M | 0.0M |
| SRR29685011\_flagstat | 4.2M | 4.2M | 4.2M | 0.0M | 0.0M | 0.0M | 0.0M | 0.0M | 0.0M | 0.0M |
| SRR29685012\_flagstat | 6.0M | 6.0M | 6.0M | 0.0M | 0.0M | 0.0M | 0.0M | 0.0M | 0.0M | 0.0M |
| SRR29685013\_flagstat | 5.3M | 5.3M | 5.3M | 0.0M | 0.0M | 0.0M | 0.0M | 0.0M | 0.0M | 0.0M |
| SRR29685014\_flagstat | 5.5M | 5.5M | 5.5M | 0.0M | 0.0M | 0.0M | 0.0M | 0.0M | 0.0M | 0.0M |
| SRR29685015\_flagstat | 4.9M | 4.9M | 4.9M | 0.0M | 0.0M | 0.0M | 0.0M | 0.0M | 0.0M | 0.0M |
| SRR29685016\_flagstat | 4.7M | 4.7M | 4.7M | 0.0M | 0.0M | 0.0M | 0.0M | 0.0M | 0.0M | 0.0M |
| SRR29685017\_flagstat | 4.2M | 4.2M | 4.2M | 0.0M | 0.0M | 0.0M | 0.0M | 0.0M | 0.0M | 0.0M |
| SRR29685018\_flagstat | 3.7M | 3.7M | 3.7M | 0.0M | 0.0M | 0.0M | 0.0M | 0.0M | 0.0M | 0.0M |
| SRR29685019\_flagstat | 5.4M | 5.4M | 5.4M | 0.0M | 0.0M | 0.0M | 0.0M | 0.0M | 0.0M | 0.0M |
| SRR29685020\_flagstat | 6.6M | 6.6M | 6.6M | 0.0M | 0.0M | 0.0M | 0.0M | 0.0M | 0.0M | 0.0M |
| SRR29685021\_flagstat | 4.3M | 4.3M | 4.3M | 0.0M | 0.0M | 0.0M | 0.0M | 0.0M | 0.0M | 0.0M |
| SRR29685022\_flagstat | 5.7M | 5.7M | 5.7M | 0.0M | 0.0M | 0.0M | 0.0M | 0.0M | 0.0M | 0.0M |
| SRR29685023\_flagstat | 6.2M | 6.2M | 6.2M | 0.0M | 0.0M | 0.0M | 0.0M | 0.0M | 0.0M | 0.0M |
| SRR29685024\_flagstat | 4.6M | 4.6M | 4.6M | 0.0M | 0.0M | 0.0M | 0.0M | 0.0M | 0.0M | 0.0M |
| SRR29685025\_flagstat | 4.2M | 4.2M | 4.2M | 0.0M | 0.0M | 0.0M | 0.0M | 0.0M | 0.0M | 0.0M |
| SRR29685026\_flagstat | 3.5M | 3.5M | 3.5M | 0.0M | 0.0M | 0.0M | 0.0M | 0.0M | 0.0M | 0.0M |
| SRR29685027\_flagstat | 4.2M | 4.2M | 4.2M | 0.0M | 0.0M | 0.0M | 0.0M | 0.0M | 0.0M | 0.0M |
| SRR29685028\_flagstat | 2.4M | 2.4M | 2.4M | 0.0M | 0.0M | 0.0M | 0.0M | 0.0M | 0.0M | 0.0M |
| SRR29685029\_flagstat | 3.8M | 3.8M | 3.8M | 0.0M | 0.0M | 0.0M | 0.0M | 0.0M | 0.0M | 0.0M |
| SRR29685030\_flagstat | 4.5M | 4.5M | 4.5M | 0.0M | 0.0M | 0.0M | 0.0M | 0.0M | 0.0M | 0.0M |
| SRR29685031\_flagstat | 4.9M | 4.9M | 4.9M | 0.0M | 0.0M | 0.0M | 0.0M | 0.0M | 0.0M | 0.0M |
| SRR29685032\_flagstat | 4.4M | 4.4M | 4.4M | 0.0M | 0.0M | 0.0M | 0.0M | 0.0M | 0.0M | 0.0M |
| SRR29685033\_flagstat | 3.0M | 3.0M | 3.0M | 0.0M | 0.0M | 0.0M | 0.0M | 0.0M | 0.0M | 0.0M |
| SRR29685034\_flagstat | 3.7M | 3.7M | 3.7M | 0.0M | 0.0M | 0.0M | 0.0M | 0.0M | 0.0M | 0.0M |
| SRR29685035\_flagstat | 3.4M | 3.4M | 3.4M | 0.0M | 0.0M | 0.0M | 0.0M | 0.0M | 0.0M | 0.0M |
| SRR29685036\_flagstat | 4.2M | 4.2M | 4.2M | 0.0M | 0.0M | 0.0M | 0.0M | 0.0M | 0.0M | 0.0M |
| SRR29685037\_flagstat | 3.4M | 3.4M | 3.4M | 0.0M | 0.0M | 0.0M | 0.0M | 0.0M | 0.0M | 0.0M |
| SRR29685038\_flagstat | 3.6M | 3.6M | 3.6M | 0.0M | 0.0M | 0.0M | 0.0M | 0.0M | 0.0M | 0.0M |
| SRR29685039\_flagstat | 3.7M | 3.7M | 3.7M | 0.0M | 0.0M | 0.0M | 0.0M | 0.0M | 0.0M | 0.0M |
| SRR29685040\_flagstat | 4.0M | 4.0M | 4.0M | 0.0M | 0.0M | 0.0M | 0.0M | 0.0M | 0.0M | 0.0M |
| SRR29685041\_flagstat | 3.8M | 3.8M | 3.8M | 0.0M | 0.0M | 0.0M | 0.0M | 0.0M | 0.0M | 0.0M |
| SRR29685042\_flagstat | 4.1M | 4.1M | 4.1M | 0.0M | 0.0M | 0.0M | 0.0M | 0.0M | 0.0M | 0.0M |
| SRR29685043\_flagstat | 3.6M | 3.6M | 3.6M | 0.0M | 0.0M | 0.0M | 0.0M | 0.0M | 0.0M | 0.0M |
| SRR29685044\_flagstat | 5.3M | 5.3M | 5.3M | 0.0M | 0.0M | 0.0M | 0.0M | 0.0M | 0.0M | 0.0M |
| SRR29685045\_flagstat | 4.0M | 4.0M | 4.0M | 0.0M | 0.0M | 0.0M | 0.0M | 0.0M | 0.0M | 0.0M |
| SRR29685046\_flagstat | 4.1M | 4.1M | 4.1M | 0.0M | 0.0M | 0.0M | 0.0M | 0.0M | 0.0M | 0.0M |
| SRR29685047\_flagstat | 3.8M | 3.8M | 3.8M | 0.0M | 0.0M | 0.0M | 0.0M | 0.0M | 0.0M | 0.0M |
| SRR29685048\_flagstat | 4.0M | 4.0M | 4.0M | 0.0M | 0.0M | 0.0M | 0.0M | 0.0M | 0.0M | 0.0M |
| SRR29685049\_flagstat | 3.9M | 3.9M | 3.9M | 0.0M | 0.0M | 0.0M | 0.0M | 0.0M | 0.0M | 0.0M |
| SRR29685050\_flagstat | 3.5M | 3.5M | 3.5M | 0.0M | 0.0M | 0.0M | 0.0M | 0.0M | 0.0M | 0.0M |
| SRR29685051\_flagstat | 4.9M | 4.9M | 4.9M | 0.0M | 0.0M | 0.0M | 0.0M | 0.0M | 0.0M | 0.0M |
| SRR29685052\_flagstat | 2.6M | 2.6M | 2.6M | 0.0M | 0.0M | 0.0M | 0.0M | 0.0M | 0.0M | 0.0M |
| SRR29685053\_flagstat | 4.1M | 4.1M | 4.1M | 0.0M | 0.0M | 0.0M | 0.0M | 0.0M | 0.0M | 0.0M |
| SRR29685054\_flagstat | 4.4M | 4.4M | 4.4M | 0.0M | 0.0M | 0.0M | 0.0M | 0.0M | 0.0M | 0.0M |
| SRR29685055\_flagstat | 3.8M | 3.8M | 3.8M | 0.0M | 0.0M | 0.0M | 0.0M | 0.0M | 0.0M | 0.0M |
| SRR29685056\_flagstat | 3.5M | 3.5M | 3.5M | 0.0M | 0.0M | 0.0M | 0.0M | 0.0M | 0.0M | 0.0M |
| SRR29685057\_flagstat | 3.7M | 3.7M | 3.7M | 0.0M | 0.0M | 0.0M | 0.0M | 0.0M | 0.0M | 0.0M |
| SRR29685058\_flagstat | 3.3M | 3.3M | 3.3M | 0.0M | 0.0M | 0.0M | 0.0M | 0.0M | 0.0M | 0.0M |
| SRR29685059\_flagstat | 4.4M | 4.4M | 4.4M | 0.0M | 0.0M | 0.0M | 0.0M | 0.0M | 0.0M | 0.0M |
| SRR29685060\_flagstat | 3.8M | 3.8M | 3.8M | 0.0M | 0.0M | 0.0M | 0.0M | 0.0M | 0.0M | 0.0M |
| SRR29685061\_flagstat | 4.0M | 4.0M | 4.0M | 0.0M | 0.0M | 0.0M | 0.0M | 0.0M | 0.0M | 0.0M |
| SRR29685062\_flagstat | 4.1M | 4.1M | 4.1M | 0.0M | 0.0M | 0.0M | 0.0M | 0.0M | 0.0M | 0.0M |
| SRR29685063\_flagstat | 3.6M | 3.6M | 3.6M | 0.0M | 0.0M | 0.0M | 0.0M | 0.0M | 0.0M | 0.0M |
| SRR29685064\_flagstat | 2.7M | 2.7M | 2.7M | 0.0M | 0.0M | 0.0M | 0.0M | 0.0M | 0.0M | 0.0M |
| SRR29685065\_flagstat | 3.5M | 3.5M | 3.5M | 0.0M | 0.0M | 0.0M | 0.0M | 0.0M | 0.0M | 0.0M |
| SRR29685066\_flagstat | 3.6M | 3.6M | 3.6M | 0.0M | 0.0M | 0.0M | 0.0M | 0.0M | 0.0M | 0.0M |
| SRR29685067\_flagstat | 5.0M | 5.0M | 5.0M | 0.0M | 0.0M | 0.0M | 0.0M | 0.0M | 0.0M | 0.0M |
| SRR29685068\_flagstat | 4.0M | 4.0M | 4.0M | 0.0M | 0.0M | 0.0M | 0.0M | 0.0M | 0.0M | 0.0M |
| SRR29685069\_flagstat | 2.6M | 2.6M | 2.6M | 0.0M | 0.0M | 0.0M | 0.0M | 0.0M | 0.0M | 0.0M |
| SRR29685070\_flagstat | 4.3M | 4.3M | 4.3M | 0.0M | 0.0M | 0.0M | 0.0M | 0.0M | 0.0M | 0.0M |
| SRR29685071\_flagstat | 4.2M | 4.2M | 4.2M | 0.0M | 0.0M | 0.0M | 0.0M | 0.0M | 0.0M | 0.0M |
| SRR29685072\_flagstat | 3.5M | 3.5M | 3.5M | 0.0M | 0.0M | 0.0M | 0.0M | 0.0M | 0.0M | 0.0M |
| SRR29685073\_flagstat | 2.2M | 2.2M | 2.2M | 0.0M | 0.0M | 0.0M | 0.0M | 0.0M | 0.0M | 0.0M |
| SRR29685074\_flagstat | 1.5M | 1.5M | 1.5M | 0.0M | 0.0M | 0.0M | 0.0M | 0.0M | 0.0M | 0.0M |
| SRR29685075\_flagstat | 2.9M | 2.9M | 2.9M | 0.0M | 0.0M | 0.0M | 0.0M | 0.0M | 0.0M | 0.0M |
| SRR29685076\_flagstat | 2.5M | 2.5M | 2.5M | 0.0M | 0.0M | 0.0M | 0.0M | 0.0M | 0.0M | 0.0M |
| SRR29685077\_flagstat | 2.6M | 2.6M | 2.6M | 0.0M | 0.0M | 0.0M | 0.0M | 0.0M | 0.0M | 0.0M |
| SRR29685078\_flagstat | 2.7M | 2.7M | 2.7M | 0.0M | 0.0M | 0.0M | 0.0M | 0.0M | 0.0M | 0.0M |
| SRR29685079\_flagstat | 2.4M | 2.4M | 2.4M | 0.0M | 0.0M | 0.0M | 0.0M | 0.0M | 0.0M | 0.0M |
| SRR29685080\_flagstat | 2.5M | 2.5M | 2.5M | 0.0M | 0.0M | 0.0M | 0.0M | 0.0M | 0.0M | 0.0M |
| SRR29685081\_flagstat | 1.0M | 1.0M | 1.0M | 0.0M | 0.0M | 0.0M | 0.0M | 0.0M | 0.0M | 0.0M |
| SRR29685082\_flagstat | 0.1M | 0.1M | 0.1M | 0.0M | 0.0M | 0.0M | 0.0M | 0.0M | 0.0M | 0.0M |
| SRR29685083\_flagstat | 2.6M | 2.6M | 2.6M | 0.0M | 0.0M | 0.0M | 0.0M | 0.0M | 0.0M | 0.0M |
| SRR29685084\_flagstat | 1.3M | 1.3M | 1.3M | 0.0M | 0.0M | 0.0M | 0.0M | 0.0M | 0.0M | 0.0M |
| SRR29685085\_flagstat | 2.1M | 2.1M | 2.1M | 0.0M | 0.0M | 0.0M | 0.0M | 0.0M | 0.0M | 0.0M |
| SRR29685086\_flagstat | 1.6M | 1.6M | 1.6M | 0.0M | 0.0M | 0.0M | 0.0M | 0.0M | 0.0M | 0.0M |
| SRR29685087\_flagstat | 1.7M | 1.7M | 1.7M | 0.0M | 0.0M | 0.0M | 0.0M | 0.0M | 0.0M | 0.0M |
| SRR29685088\_flagstat | 1.5M | 1.5M | 1.5M | 0.0M | 0.0M | 0.0M | 0.0M | 0.0M | 0.0M | 0.0M |
| SRR29685089\_flagstat | 3.5M | 3.5M | 3.5M | 0.0M | 0.0M | 0.0M | 0.0M | 0.0M | 0.0M | 0.0M |
| SRR29685090\_flagstat | 3.3M | 3.3M | 3.3M | 0.0M | 0.0M | 0.0M | 0.0M | 0.0M | 0.0M | 0.0M |
| SRR29685091\_flagstat | 5.1M | 5.1M | 5.1M | 0.0M | 0.0M | 0.0M | 0.0M | 0.0M | 0.0M | 0.0M |
| SRR29685092\_flagstat | 6.2M | 6.2M | 6.2M | 0.0M | 0.0M | 0.0M | 0.0M | 0.0M | 0.0M | 0.0M |
| SRR29685093\_flagstat | 6.2M | 6.2M | 6.2M | 0.0M | 0.0M | 0.0M | 0.0M | 0.0M | 0.0M | 0.0M |
| SRR29685094\_flagstat | 5.1M | 5.1M | 5.1M | 0.0M | 0.0M | 0.0M | 0.0M | 0.0M | 0.0M | 0.0M |
| SRR29685095\_flagstat | 5.9M | 5.9M | 5.9M | 0.0M | 0.0M | 0.0M | 0.0M | 0.0M | 0.0M | 0.0M |
| SRR29685096\_flagstat | 4.0M | 4.0M | 4.0M | 0.0M | 0.0M | 0.0M | 0.0M | 0.0M | 0.0M | 0.0M |
| SRR29685097\_flagstat | 2.6M | 2.6M | 2.6M | 0.0M | 0.0M | 0.0M | 0.0M | 0.0M | 0.0M | 0.0M |
| SRR29685098\_flagstat | 3.2M | 3.2M | 3.2M | 0.0M | 0.0M | 0.0M | 0.0M | 0.0M | 0.0M | 0.0M |
| SRR29685099\_flagstat | 3.5M | 3.5M | 3.5M | 0.0M | 0.0M | 0.0M | 0.0M | 0.0M | 0.0M | 0.0M |
| SRR29685100\_flagstat | 5.0M | 5.0M | 5.0M | 0.0M | 0.0M | 0.0M | 0.0M | 0.0M | 0.0M | 0.0M |
| SRR29685101\_flagstat | 3.3M | 3.3M | 3.3M | 0.0M | 0.0M | 0.0M | 0.0M | 0.0M | 0.0M | 0.0M |
| SRR29685102\_flagstat | 4.1M | 4.1M | 4.1M | 0.0M | 0.0M | 0.0M | 0.0M | 0.0M | 0.0M | 0.0M |
| SRR29685103\_flagstat | 4.7M | 4.7M | 4.7M | 0.0M | 0.0M | 0.0M | 0.0M | 0.0M | 0.0M | 0.0M |
| SRR29685104\_flagstat | 3.6M | 3.6M | 3.6M | 0.0M | 0.0M | 0.0M | 0.0M | 0.0M | 0.0M | 0.0M |
| SRR29685105\_flagstat | 3.5M | 3.5M | 3.5M | 0.0M | 0.0M | 0.0M | 0.0M | 0.0M | 0.0M | 0.0M |
| SRR29685106\_flagstat | 1.8M | 1.8M | 1.8M | 0.0M | 0.0M | 0.0M | 0.0M | 0.0M | 0.0M | 0.0M |
| SRR29685107\_flagstat | 0.1M | 0.1M | 0.1M | 0.0M | 0.0M | 0.0M | 0.0M | 0.0M | 0.0M | 0.0M |
| SRR29685108\_flagstat | 5.0M | 5.0M | 5.0M | 0.0M | 0.0M | 0.0M | 0.0M | 0.0M | 0.0M | 0.0M |
| SRR29685109\_flagstat | 4.1M | 4.1M | 4.1M | 0.0M | 0.0M | 0.0M | 0.0M | 0.0M | 0.0M | 0.0M |
| SRR29685110\_flagstat | 4.7M | 4.7M | 4.7M | 0.0M | 0.0M | 0.0M | 0.0M | 0.0M | 0.0M | 0.0M |
| SRR29685111\_flagstat | 4.5M | 4.5M | 4.5M | 0.0M | 0.0M | 0.0M | 0.0M | 0.0M | 0.0M | 0.0M |
| SRR29685112\_flagstat | 3.8M | 3.8M | 3.8M | 0.0M | 0.0M | 0.0M | 0.0M | 0.0M | 0.0M | 0.0M |

×

###### Samtools: flagstat: read count: Columns

Uncheck the tick box to hide columns. Click and drag the handle on the left to change order. Table ID: `samtools-flagstat-dp_table-1`

Show All
Show None

| Sort | Visible | Group | Column | Description | ID | Scale |
| --- | --- | --- | --- | --- | --- | --- |
| \|\| |  |  | Total Reads | Total Reads | `flagstat_total` | read\_count |
| \|\| |  |  | Total Passed QC | Total Passed QC | `total_passed` | read\_count |
| \|\| |  |  | Mapped | Mapped | `mapped_passed` | read\_count |
| \|\| |  |  | Duplicates | Duplicates | `duplicates_passed` | read\_count |
| \|\| |  |  | Paired in Sequencing | Paired in Sequencing | `paired_in_sequencing_passed` | read\_count |
| \|\| |  |  | Properly Paired | Properly Paired | `properly_paired_passed` | read\_count |
| \|\| |  |  | Self and mate mapped | Reads with itself and mate mapped | `with_itself_and_mate_mapped_passed` | read\_count |
| \|\| |  |  | Singletons | Singletons | `singletons_passed` | read\_count |
| \|\| |  |  | Mate mapped to diff chr | Mate mapped to different chromosome | `with_mate_mapped_to_a_different_chr_passed` | read\_count |
| \|\| |  |  | Diff chr (mapQ >= 5) | Mate mapped to different chromosome (mapQ >= 5) | `with_mate_mapped_to_a_different_chr_mapQ_5__passed` | read\_count |

Close

---

##### Mapped reads per contig

The `samtools idxstats` tool counts the number of mapped reads per chromosome / contig. Chromosomes with < 0.1% of the total aligned reads are omitted from this plot.

Log10

Normalised Counts
Observed over Expected Counts
Raw Counts

Export Plot

Created with MultiQC

---

#### STAR

Universal RNA-seq aligner.*URL: https://github.com/alexdobin/STAR**DOI: 10.1093/bioinformatics/bts635*

##### Summary Statistics

Summary statistics from the STAR alignment

Configure columns
 Table
Export Plot

Created with MultiQC

Copy table

 Configure columns

 Sort by highlight

 Scatter plot

 Violin plot
Export as CSV
Showing ^0^/_286_ rows and ^10^/_19_ columns.

| Sample Name | Total reads | Aligned | Aligned | Uniq aligned | Uniq aligned | Multimapped | Avg. read len | Avg. mapped len | Splices | Annotated splices | GT/AG splices | GC/AG splices | AT/AC splices | Non-canonical splices | Mismatch rate | Del rate | Del len | Ins rate | Ins len |
| --- | --- | --- | --- | --- | --- | --- | --- | --- | --- | --- | --- | --- | --- | --- | --- | --- | --- | --- | --- |
| SRR29684827 | 4.0M | 3.9M | 97.6% | 3.9M | 97.6% | 0.0M | 50.0bp | 49.7bp | 0.0M | 0.0M | 0.0M | 0.0M | 0.0M | 0.0M | 0.1% | 0.1% | 1.0bp | 0.0% | 1.0bp |
| SRR29684828 | 1.6M | 0.1M | 6.7% | 0.1M | 6.7% | 0.0M | 51.0bp | 49.7bp | 0.0M | 0.0M | 0.0M | 0.0M | 0.0M | 0.0M | 0.2% | 0.2% | 1.0bp | 0.0% | 1.0bp |
| SRR29684829 | 4.2M | 3.7M | 89.4% | 3.7M | 89.4% | 0.0M | 51.0bp | 49.8bp | 0.0M | 0.0M | 0.0M | 0.0M | 0.0M | 0.0M | 0.2% | 0.1% | 1.0bp | 0.0% | 1.0bp |
| SRR29684830 | 3.5M | 3.4M | 97.1% | 3.4M | 97.1% | 0.0M | 50.0bp | 49.8bp | 0.0M | 0.0M | 0.0M | 0.0M | 0.0M | 0.0M | 0.1% | 0.1% | 1.0bp | 0.0% | 1.0bp |
| SRR29684831 | 4.5M | 4.3M | 96.4% | 4.3M | 96.4% | 0.0M | 51.0bp | 49.8bp | 0.0M | 0.0M | 0.0M | 0.0M | 0.0M | 0.0M | 0.1% | 0.1% | 1.0bp | 0.0% | 1.0bp |
| SRR29684832 | 4.5M | 4.4M | 97.1% | 4.4M | 97.1% | 0.0M | 51.0bp | 49.8bp | 0.0M | 0.0M | 0.0M | 0.0M | 0.0M | 0.0M | 0.1% | 0.1% | 1.0bp | 0.0% | 1.0bp |
| SRR29684833 | 4.5M | 4.4M | 98.2% | 4.4M | 98.2% | 0.0M | 50.0bp | 49.8bp | 0.0M | 0.0M | 0.0M | 0.0M | 0.0M | 0.0M | 0.1% | 0.1% | 1.0bp | 0.0% | 1.0bp |
| SRR29684834 | 3.7M | 3.7M | 97.9% | 3.7M | 97.9% | 0.0M | 50.0bp | 49.8bp | 0.0M | 0.0M | 0.0M | 0.0M | 0.0M | 0.0M | 0.1% | 0.1% | 1.0bp | 0.0% | 1.0bp |
| SRR29684835 | 3.8M | 3.7M | 97.7% | 3.7M | 97.7% | 0.0M | 51.0bp | 49.8bp | 0.0M | 0.0M | 0.0M | 0.0M | 0.0M | 0.0M | 0.1% | 0.1% | 1.0bp | 0.0% | 1.0bp |
| SRR29684836 | 4.0M | 4.0M | 98.4% | 4.0M | 98.4% | 0.0M | 51.0bp | 49.8bp | 0.0M | 0.0M | 0.0M | 0.0M | 0.0M | 0.0M | 0.1% | 0.1% | 1.0bp | 0.0% | 1.0bp |
| SRR29684837 | 5.0M | 4.8M | 96.9% | 4.8M | 96.9% | 0.0M | 51.0bp | 49.8bp | 0.0M | 0.0M | 0.0M | 0.0M | 0.0M | 0.0M | 0.1% | 0.1% | 1.0bp | 0.0% | 1.0bp |
| SRR29684838 | 4.0M | 3.9M | 97.2% | 3.9M | 97.2% | 0.0M | 51.0bp | 49.8bp | 0.0M | 0.0M | 0.0M | 0.0M | 0.0M | 0.0M | 0.1% | 0.1% | 1.0bp | 0.0% | 1.0bp |
| SRR29684839 | 4.7M | 4.6M | 97.6% | 4.6M | 97.6% | 0.0M | 51.0bp | 49.8bp | 0.0M | 0.0M | 0.0M | 0.0M | 0.0M | 0.0M | 0.1% | 0.1% | 1.0bp | 0.0% | 1.0bp |
| SRR29684840 | 4.3M | 4.1M | 97.1% | 4.1M | 97.1% | 0.0M | 51.0bp | 49.8bp | 0.0M | 0.0M | 0.0M | 0.0M | 0.0M | 0.0M | 0.1% | 0.1% | 1.0bp | 0.0% | 1.0bp |
| SRR29684841 | 3.6M | 3.5M | 97.6% | 3.5M | 97.6% | 0.0M | 50.0bp | 49.8bp | 0.0M | 0.0M | 0.0M | 0.0M | 0.0M | 0.0M | 0.1% | 0.1% | 1.0bp | 0.0% | 1.0bp |
| SRR29684842 | 3.1M | 3.0M | 97.7% | 3.0M | 97.7% | 0.0M | 51.0bp | 49.8bp | 0.0M | 0.0M | 0.0M | 0.0M | 0.0M | 0.0M | 0.1% | 0.1% | 1.0bp | 0.0% | 1.0bp |
| SRR29684843 | 4.4M | 4.3M | 97.6% | 4.3M | 97.6% | 0.0M | 51.0bp | 49.8bp | 0.0M | 0.0M | 0.0M | 0.0M | 0.0M | 0.0M | 0.1% | 0.1% | 1.0bp | 0.0% | 1.0bp |
| SRR29684844 | 5.8M | 5.7M | 98.5% | 5.7M | 98.5% | 0.0M | 50.0bp | 49.7bp | 0.0M | 0.0M | 0.0M | 0.0M | 0.0M | 0.0M | 0.1% | 0.1% | 1.0bp | 0.0% | 1.0bp |
| SRR29684845 | 6.6M | 6.4M | 97.1% | 6.4M | 97.1% | 0.0M | 51.0bp | 49.8bp | 0.0M | 0.0M | 0.0M | 0.0M | 0.0M | 0.0M | 0.1% | 0.1% | 1.0bp | 0.0% | 1.0bp |
| SRR29684846 | 4.8M | 4.6M | 97.5% | 4.6M | 97.5% | 0.0M | 51.0bp | 49.8bp | 0.0M | 0.0M | 0.0M | 0.0M | 0.0M | 0.0M | 0.1% | 0.1% | 1.0bp | 0.0% | 1.0bp |
| SRR29684847 | 6.6M | 6.4M | 97.8% | 6.4M | 97.8% | 0.0M | 51.0bp | 49.8bp | 0.0M | 0.0M | 0.0M | 0.0M | 0.0M | 0.0M | 0.1% | 0.1% | 1.0bp | 0.0% | 1.0bp |
| SRR29684848 | 5.8M | 5.6M | 97.4% | 5.6M | 97.4% | 0.0M | 51.0bp | 49.8bp | 0.0M | 0.0M | 0.0M | 0.0M | 0.0M | 0.0M | 0.1% | 0.1% | 1.0bp | 0.0% | 1.0bp |
| SRR29684849 | 4.7M | 4.6M | 97.8% | 4.6M | 97.8% | 0.0M | 51.0bp | 49.8bp | 0.0M | 0.0M | 0.0M | 0.0M | 0.0M | 0.0M | 0.1% | 0.1% | 1.0bp | 0.0% | 1.0bp |
| SRR29684850 | 3.6M | 3.5M | 97.3% | 3.5M | 97.3% | 0.0M | 51.0bp | 49.8bp | 0.0M | 0.0M | 0.0M | 0.0M | 0.0M | 0.0M | 0.1% | 0.1% | 1.0bp | 0.0% | 1.0bp |
| SRR29684851 | 1.9M | 1.8M | 93.9% | 1.8M | 93.9% | 0.0M | 51.0bp | 49.8bp | 0.0M | 0.0M | 0.0M | 0.0M | 0.0M | 0.0M | 0.2% | 0.1% | 1.0bp | 0.0% | 1.0bp |
| SRR29684852 | 1.5M | 1.4M | 93.4% | 1.4M | 93.4% | 0.0M | 51.0bp | 49.8bp | 0.0M | 0.0M | 0.0M | 0.0M | 0.0M | 0.0M | 0.2% | 0.1% | 1.0bp | 0.0% | 1.0bp |
| SRR29684853 | 2.7M | 2.6M | 96.3% | 2.6M | 96.3% | 0.0M | 51.0bp | 49.8bp | 0.0M | 0.0M | 0.0M | 0.0M | 0.0M | 0.0M | 0.2% | 0.1% | 1.0bp | 0.0% | 1.0bp |
| SRR29684854 | 2.7M | 2.6M | 97.5% | 2.6M | 97.5% | 0.0M | 51.0bp | 49.8bp | 0.0M | 0.0M | 0.0M | 0.0M | 0.0M | 0.0M | 0.1% | 0.1% | 1.0bp | 0.0% | 1.0bp |
| SRR29684855 | 1.5M | 1.3M | 84.8% | 1.3M | 84.8% | 0.0M | 51.0bp | 49.8bp | 0.0M | 0.0M | 0.0M | 0.0M | 0.0M | 0.0M | 0.2% | 0.1% | 1.0bp | 0.0% | 1.0bp |
| SRR29684856 | 0.6M | 0.4M | 65.8% | 0.4M | 65.8% | 0.0M | 51.0bp | 49.8bp | 0.0M | 0.0M | 0.0M | 0.0M | 0.0M | 0.0M | 0.2% | 0.1% | 1.0bp | 0.0% | 1.0bp |
| SRR29684857 | 1.0M | 0.8M | 80.6% | 0.8M | 80.6% | 0.0M | 51.0bp | 49.8bp | 0.0M | 0.0M | 0.0M | 0.0M | 0.0M | 0.0M | 0.2% | 0.1% | 1.0bp | 0.0% | 1.0bp |
| SRR29684858 | 1.6M | 1.5M | 92.9% | 1.5M | 92.9% | 0.0M | 51.0bp | 49.8bp | 0.0M | 0.0M | 0.0M | 0.0M | 0.0M | 0.0M | 0.2% | 0.1% | 1.0bp | 0.0% | 1.0bp |
| SRR29684859 | 2.3M | 2.2M | 94.5% | 2.2M | 94.5% | 0.0M | 51.0bp | 49.8bp | 0.0M | 0.0M | 0.0M | 0.0M | 0.0M | 0.0M | 0.2% | 0.1% | 1.0bp | 0.0% | 1.0bp |
| SRR29684860 | 2.5M | 2.4M | 97.4% | 2.4M | 97.4% | 0.0M | 51.0bp | 49.8bp | 0.0M | 0.0M | 0.0M | 0.0M | 0.0M | 0.0M | 0.2% | 0.1% | 1.0bp | 0.0% | 1.0bp |
| SRR29684861 | 2.5M | 2.4M | 97.9% | 2.4M | 97.9% | 0.0M | 51.0bp | 49.8bp | 0.0M | 0.0M | 0.0M | 0.0M | 0.0M | 0.0M | 0.1% | 0.1% | 1.0bp | 0.0% | 1.0bp |
| SRR29684862 | 1.7M | 1.6M | 95.3% | 1.6M | 95.3% | 0.0M | 51.0bp | 49.8bp | 0.0M | 0.0M | 0.0M | 0.0M | 0.0M | 0.0M | 0.2% | 0.1% | 1.0bp | 0.0% | 1.0bp |
| SRR29684863 | 2.3M | 2.2M | 93.7% | 2.2M | 93.7% | 0.0M | 51.0bp | 49.8bp | 0.0M | 0.0M | 0.0M | 0.0M | 0.0M | 0.0M | 0.2% | 0.1% | 1.0bp | 0.0% | 1.0bp |
| SRR29684864 | 1.9M | 1.8M | 93.1% | 1.8M | 93.1% | 0.0M | 51.0bp | 49.8bp | 0.0M | 0.0M | 0.0M | 0.0M | 0.0M | 0.0M | 0.2% | 0.1% | 1.0bp | 0.0% | 1.0bp |
| SRR29684865 | 1.7M | 1.6M | 94.0% | 1.6M | 94.0% | 0.0M | 51.0bp | 49.8bp | 0.0M | 0.0M | 0.0M | 0.0M | 0.0M | 0.0M | 0.2% | 0.1% | 1.0bp | 0.0% | 1.0bp |
| SRR29684866 | 1.5M | 1.4M | 93.9% | 1.4M | 93.9% | 0.0M | 51.0bp | 49.8bp | 0.0M | 0.0M | 0.0M | 0.0M | 0.0M | 0.0M | 0.2% | 0.1% | 1.0bp | 0.0% | 1.0bp |
| SRR29684867 | 1.9M | 1.7M | 92.7% | 1.7M | 92.7% | 0.0M | 51.0bp | 49.8bp | 0.0M | 0.0M | 0.0M | 0.0M | 0.0M | 0.0M | 0.2% | 0.1% | 1.0bp | 0.0% | 1.0bp |
| SRR29684868 | 3.3M | 3.2M | 97.7% | 3.2M | 97.7% | 0.0M | 51.0bp | 49.8bp | 0.0M | 0.0M | 0.0M | 0.0M | 0.0M | 0.0M | 0.2% | 0.1% | 1.0bp | 0.0% | 1.0bp |
| SRR29684869 | 2.8M | 2.8M | 97.6% | 2.8M | 97.6% | 0.0M | 51.0bp | 49.8bp | 0.0M | 0.0M | 0.0M | 0.0M | 0.0M | 0.0M | 0.2% | 0.1% | 1.0bp | 0.0% | 1.0bp |
| SRR29684870 | 2.4M | 2.3M | 96.3% | 2.3M | 96.3% | 0.0M | 51.0bp | 49.8bp | 0.0M | 0.0M | 0.0M | 0.0M | 0.0M | 0.0M | 0.1% | 0.1% | 1.0bp | 0.0% | 1.0bp |
| SRR29684871 | 3.4M | 3.3M | 97.5% | 3.3M | 97.5% | 0.0M | 51.0bp | 49.8bp | 0.0M | 0.0M | 0.0M | 0.0M | 0.0M | 0.0M | 0.2% | 0.1% | 1.0bp | 0.0% | 1.0bp |
| SRR29684872 | 3.4M | 3.3M | 97.5% | 3.3M | 97.5% | 0.0M | 51.0bp | 49.8bp | 0.0M | 0.0M | 0.0M | 0.0M | 0.0M | 0.0M | 0.1% | 0.1% | 1.0bp | 0.0% | 1.0bp |
| SRR29684873 | 3.4M | 3.3M | 97.7% | 3.3M | 97.7% | 0.0M | 51.0bp | 49.8bp | 0.0M | 0.0M | 0.0M | 0.0M | 0.0M | 0.0M | 0.2% | 0.1% | 1.0bp | 0.0% | 1.0bp |
| SRR29684874 | 0.9M | 0.7M | 77.1% | 0.7M | 77.1% | 0.0M | 51.0bp | 49.8bp | 0.0M | 0.0M | 0.0M | 0.0M | 0.0M | 0.0M | 0.2% | 0.1% | 1.0bp | 0.0% | 1.0bp |
| SRR29684875 | 2.7M | 2.6M | 95.7% | 2.6M | 95.7% | 0.0M | 51.0bp | 49.8bp | 0.0M | 0.0M | 0.0M | 0.0M | 0.0M | 0.0M | 0.2% | 0.1% | 1.0bp | 0.0% | 1.0bp |
| SRR29684876 | 4.2M | 4.1M | 97.1% | 4.1M | 97.1% | 0.0M | 51.0bp | 49.8bp | 0.0M | 0.0M | 0.0M | 0.0M | 0.0M | 0.0M | 0.1% | 0.1% | 1.0bp | 0.0% | 1.0bp |
| SRR29684877 | 4.0M | 3.8M | 96.3% | 3.8M | 96.3% | 0.0M | 51.0bp | 49.8bp | 0.0M | 0.0M | 0.0M | 0.0M | 0.0M | 0.0M | 0.1% | 0.1% | 1.0bp | 0.0% | 1.0bp |
| SRR29684878 | 5.3M | 5.1M | 97.0% | 5.1M | 97.0% | 0.0M | 50.0bp | 49.8bp | 0.0M | 0.0M | 0.0M | 0.0M | 0.0M | 0.0M | 0.1% | 0.1% | 1.0bp | 0.0% | 1.0bp |
| SRR29684879 | 4.5M | 4.4M | 97.1% | 4.4M | 97.1% | 0.0M | 51.0bp | 49.8bp | 0.0M | 0.0M | 0.0M | 0.0M | 0.0M | 0.0M | 0.1% | 0.1% | 1.0bp | 0.0% | 1.0bp |
| SRR29684880 | 4.9M | 4.8M | 97.4% | 4.8M | 97.4% | 0.0M | 51.0bp | 49.8bp | 0.0M | 0.0M | 0.0M | 0.0M | 0.0M | 0.0M | 0.1% | 0.1% | 1.0bp | 0.0% | 1.0bp |
| SRR29684881 | 4.4M | 4.3M | 97.3% | 4.3M | 97.3% | 0.0M | 51.0bp | 49.8bp | 0.0M | 0.0M | 0.0M | 0.0M | 0.0M | 0.0M | 0.1% | 0.1% | 1.0bp | 0.0% | 1.0bp |
| SRR29684882 | 4.2M | 4.2M | 97.9% | 4.2M | 97.9% | 0.0M | 50.0bp | 49.8bp | 0.0M | 0.0M | 0.0M | 0.0M | 0.0M | 0.0M | 0.1% | 0.1% | 1.0bp | 0.0% | 1.0bp |
| SRR29684883 | 3.2M | 3.1M | 95.4% | 3.1M | 95.4% | 0.0M | 51.0bp | 49.8bp | 0.0M | 0.0M | 0.0M | 0.0M | 0.0M | 0.0M | 0.1% | 0.1% | 1.0bp | 0.0% | 1.0bp |
| SRR29684884 | 3.4M | 3.3M | 97.5% | 3.3M | 97.5% | 0.0M | 51.0bp | 49.8bp | 0.0M | 0.0M | 0.0M | 0.0M | 0.0M | 0.0M | 0.1% | 0.1% | 1.0bp | 0.0% | 1.0bp |
| SRR29684885 | 2.3M | 2.2M | 97.6% | 2.2M | 97.6% | 0.0M | 50.0bp | 49.7bp | 0.0M | 0.0M | 0.0M | 0.0M | 0.0M | 0.0M | 0.1% | 0.1% | 1.0bp | 0.0% | 1.0bp |
| SRR29684886 | 5.1M | 5.0M | 97.0% | 5.0M | 97.0% | 0.0M | 51.0bp | 49.8bp | 0.0M | 0.0M | 0.0M | 0.0M | 0.0M | 0.0M | 0.1% | 0.1% | 1.0bp | 0.0% | 1.0bp |
| SRR29684887 | 3.8M | 3.7M | 97.4% | 3.7M | 97.4% | 0.0M | 51.0bp | 49.8bp | 0.0M | 0.0M | 0.0M | 0.0M | 0.0M | 0.0M | 0.1% | 0.1% | 1.0bp | 0.0% | 1.0bp |
| SRR29684888 | 4.3M | 4.2M | 97.3% | 4.2M | 97.3% | 0.0M | 51.0bp | 49.8bp | 0.0M | 0.0M | 0.0M | 0.0M | 0.0M | 0.0M | 0.1% | 0.1% | 1.0bp | 0.0% | 1.0bp |
| SRR29684889 | 4.4M | 4.2M | 97.0% | 4.2M | 97.0% | 0.0M | 51.0bp | 49.8bp | 0.0M | 0.0M | 0.0M | 0.0M | 0.0M | 0.0M | 0.1% | 0.1% | 1.0bp | 0.0% | 1.0bp |
| SRR29684890 | 3.3M | 3.1M | 94.8% | 3.1M | 94.8% | 0.0M | 51.0bp | 49.8bp | 0.0M | 0.0M | 0.0M | 0.0M | 0.0M | 0.0M | 0.1% | 0.1% | 1.0bp | 0.0% | 1.0bp |
| SRR29684891 | 2.8M | 2.7M | 96.6% | 2.7M | 96.6% | 0.0M | 51.0bp | 49.8bp | 0.0M | 0.0M | 0.0M | 0.0M | 0.0M | 0.0M | 0.1% | 0.1% | 1.0bp | 0.0% | 1.0bp |
| SRR29684892 | 3.7M | 3.5M | 96.0% | 3.5M | 96.0% | 0.0M | 51.0bp | 49.8bp | 0.0M | 0.0M | 0.0M | 0.0M | 0.0M | 0.0M | 0.1% | 0.1% | 1.0bp | 0.0% | 1.0bp |
| SRR29684893 | 4.4M | 4.2M | 96.6% | 4.2M | 96.6% | 0.0M | 51.0bp | 49.8bp | 0.0M | 0.0M | 0.0M | 0.0M | 0.0M | 0.0M | 0.1% | 0.1% | 1.0bp | 0.0% | 1.0bp |
| SRR29684894 | 6.5M | 6.4M | 97.6% | 6.4M | 97.6% | 0.0M | 51.0bp | 49.8bp | 0.0M | 0.0M | 0.0M | 0.0M | 0.0M | 0.0M | 0.1% | 0.1% | 1.0bp | 0.0% | 1.0bp |
| SRR29684895 | 4.8M | 4.7M | 97.5% | 4.7M | 97.5% | 0.0M | 51.0bp | 49.8bp | 0.0M | 0.0M | 0.0M | 0.0M | 0.0M | 0.0M | 0.1% | 0.1% | 1.0bp | 0.0% | 1.0bp |
| SRR29684896 | 5.1M | 5.0M | 97.7% | 5.0M | 97.7% | 0.0M | 51.0bp | 49.8bp | 0.0M | 0.0M | 0.0M | 0.0M | 0.0M | 0.0M | 0.1% | 0.1% | 1.0bp | 0.0% | 1.0bp |
| SRR29684897 | 4.6M | 4.5M | 97.5% | 4.5M | 97.5% | 0.0M | 51.0bp | 49.8bp | 0.0M | 0.0M | 0.0M | 0.0M | 0.0M | 0.0M | 0.1% | 0.1% | 1.0bp | 0.0% | 1.0bp |
| SRR29684898 | 4.9M | 4.8M | 98.2% | 4.8M | 98.2% | 0.0M | 50.0bp | 49.7bp | 0.0M | 0.0M | 0.0M | 0.0M | 0.0M | 0.0M | 0.1% | 0.1% | 1.0bp | 0.0% | 1.0bp |
| SRR29684899 | 3.1M | 3.0M | 97.0% | 3.0M | 97.0% | 0.0M | 51.0bp | 49.8bp | 0.0M | 0.0M | 0.0M | 0.0M | 0.0M | 0.0M | 0.1% | 0.1% | 1.0bp | 0.0% | 1.0bp |
| SRR29684900 | 3.5M | 3.4M | 97.0% | 3.4M | 97.0% | 0.0M | 51.0bp | 49.8bp | 0.0M | 0.0M | 0.0M | 0.0M | 0.0M | 0.0M | 0.1% | 0.1% | 1.0bp | 0.0% | 1.0bp |
| SRR29684901 | 4.8M | 4.7M | 96.7% | 4.7M | 96.7% | 0.0M | 51.0bp | 49.8bp | 0.0M | 0.0M | 0.0M | 0.0M | 0.0M | 0.0M | 0.1% | 0.1% | 1.0bp | 0.0% | 1.0bp |
| SRR29684902 | 4.2M | 4.2M | 98.5% | 4.2M | 98.5% | 0.0M | 51.0bp | 49.8bp | 0.0M | 0.0M | 0.0M | 0.0M | 0.0M | 0.0M | 0.1% | 0.1% | 1.0bp | 0.0% | 1.0bp |
| SRR29684903 | 4.8M | 4.7M | 98.3% | 4.7M | 98.3% | 0.0M | 51.0bp | 49.8bp | 0.0M | 0.0M | 0.0M | 0.0M | 0.0M | 0.0M | 0.1% | 0.1% | 1.0bp | 0.0% | 1.0bp |
| SRR29684904 | 4.1M | 4.0M | 96.2% | 4.0M | 96.2% | 0.0M | 51.0bp | 49.8bp | 0.0M | 0.0M | 0.0M | 0.0M | 0.0M | 0.0M | 0.1% | 0.1% | 1.0bp | 0.0% | 1.0bp |
| SRR29684905 | 1.5M | 1.0M | 64.0% | 1.0M | 64.0% | 0.0M | 51.0bp | 49.8bp | 0.0M | 0.0M | 0.0M | 0.0M | 0.0M | 0.0M | 0.2% | 0.1% | 1.0bp | 0.0% | 1.0bp |
| SRR29684906 | 3.4M | 3.3M | 96.4% | 3.3M | 96.4% | 0.0M | 51.0bp | 49.8bp | 0.0M | 0.0M | 0.0M | 0.0M | 0.0M | 0.0M | 0.1% | 0.1% | 1.0bp | 0.0% | 1.0bp |
| SRR29684907 | 4.0M | 3.9M | 98.4% | 3.9M | 98.4% | 0.0M | 50.0bp | 49.7bp | 0.0M | 0.0M | 0.0M | 0.0M | 0.0M | 0.0M | 0.1% | 0.1% | 1.0bp | 0.0% | 1.0bp |
| SRR29684908 | 2.4M | 1.9M | 82.6% | 1.9M | 82.6% | 0.0M | 51.0bp | 49.8bp | 0.0M | 0.0M | 0.0M | 0.0M | 0.0M | 0.0M | 0.2% | 0.1% | 1.0bp | 0.0% | 1.0bp |
| SRR29684909 | 5.2M | 5.0M | 97.5% | 5.0M | 97.5% | 0.0M | 51.0bp | 49.8bp | 0.0M | 0.0M | 0.0M | 0.0M | 0.0M | 0.0M | 0.1% | 0.1% | 1.0bp | 0.0% | 1.0bp |
| SRR29684910 | 3.7M | 3.6M | 97.9% | 3.6M | 97.9% | 0.0M | 50.0bp | 49.8bp | 0.0M | 0.0M | 0.0M | 0.0M | 0.0M | 0.0M | 0.1% | 0.1% | 1.0bp | 0.0% | 1.0bp |
| SRR29684911 | 3.9M | 3.8M | 97.9% | 3.8M | 97.9% | 0.0M | 50.0bp | 49.8bp | 0.0M | 0.0M | 0.0M | 0.0M | 0.0M | 0.0M | 0.1% | 0.1% | 1.0bp | 0.0% | 1.0bp |
| SRR29684912 | 4.4M | 4.4M | 98.2% | 4.4M | 98.2% | 0.0M | 50.0bp | 49.8bp | 0.0M | 0.0M | 0.0M | 0.0M | 0.0M | 0.0M | 0.1% | 0.1% | 1.0bp | 0.0% | 1.0bp |
| SRR29684913 | 4.5M | 4.5M | 98.3% | 4.5M | 98.3% | 0.0M | 50.0bp | 49.8bp | 0.0M | 0.0M | 0.0M | 0.0M | 0.0M | 0.0M | 0.1% | 0.1% | 1.0bp | 0.0% | 1.0bp |
| SRR29684914 | 1.2M | 1.0M | 87.4% | 1.0M | 87.4% | 0.0M | 51.0bp | 49.8bp | 0.0M | 0.0M | 0.0M | 0.0M | 0.0M | 0.0M | 0.2% | 0.1% | 1.0bp | 0.0% | 1.0bp |
| SRR29684915 | 1.3M | 1.2M | 87.5% | 1.2M | 87.5% | 0.0M | 51.0bp | 49.8bp | 0.0M | 0.0M | 0.0M | 0.0M | 0.0M | 0.0M | 0.2% | 0.1% | 1.0bp | 0.0% | 1.0bp |
| SRR29684916 | 1.7M | 1.5M | 87.8% | 1.5M | 87.8% | 0.0M | 51.0bp | 49.8bp | 0.0M | 0.0M | 0.0M | 0.0M | 0.0M | 0.0M | 0.2% | 0.1% | 1.0bp | 0.0% | 1.0bp |
| SRR29684917 | 2.7M | 2.6M | 97.4% | 2.6M | 97.4% | 0.0M | 51.0bp | 49.8bp | 0.0M | 0.0M | 0.0M | 0.0M | 0.0M | 0.0M | 0.2% | 0.1% | 1.0bp | 0.0% | 1.0bp |
| SRR29684918 | 0.4M | 0.0M | 7.8% | 0.0M | 7.8% | 0.0M | 51.0bp | 49.7bp | 0.0M | 0.0M | 0.0M | 0.0M | 0.0M | 0.0M | 0.2% | 0.2% | 1.0bp | 0.0% | 1.0bp |
| SRR29684919 | 2.7M | 2.5M | 93.8% | 2.5M | 93.8% | 0.0M | 51.0bp | 49.8bp | 0.0M | 0.0M | 0.0M | 0.0M | 0.0M | 0.0M | 0.2% | 0.1% | 1.0bp | 0.0% | 1.0bp |
| SRR29684920 | 2.1M | 2.0M | 94.9% | 2.0M | 94.9% | 0.0M | 51.0bp | 49.8bp | 0.0M | 0.0M | 0.0M | 0.0M | 0.0M | 0.0M | 0.2% | 0.1% | 1.0bp | 0.0% | 1.0bp |
| SRR29684921 | 2.2M | 2.1M | 94.6% | 2.1M | 94.6% | 0.0M | 51.0bp | 49.8bp | 0.0M | 0.0M | 0.0M | 0.0M | 0.0M | 0.0M | 0.2% | 0.1% | 1.0bp | 0.0% | 1.0bp |
| SRR29684922 | 1.5M | 1.4M | 93.4% | 1.4M | 93.4% | 0.0M | 51.0bp | 49.8bp | 0.0M | 0.0M | 0.0M | 0.0M | 0.0M | 0.0M | 0.2% | 0.1% | 1.0bp | 0.0% | 1.0bp |
| SRR29684923 | 2.0M | 1.9M | 92.0% | 1.9M | 92.0% | 0.0M | 51.0bp | 49.8bp | 0.0M | 0.0M | 0.0M | 0.0M | 0.0M | 0.0M | 0.2% | 0.1% | 1.0bp | 0.0% | 1.0bp |
| SRR29684924 | 2.2M | 2.1M | 95.3% | 2.1M | 95.3% | 0.0M | 51.0bp | 49.8bp | 0.0M | 0.0M | 0.0M | 0.0M | 0.0M | 0.0M | 0.2% | 0.1% | 1.0bp | 0.0% | 1.0bp |
| SRR29684925 | 1.3M | 1.1M | 89.7% | 1.1M | 89.7% | 0.0M | 51.0bp | 49.8bp | 0.0M | 0.0M | 0.0M | 0.0M | 0.0M | 0.0M | 0.2% | 0.1% | 1.0bp | 0.0% | 1.0bp |
| SRR29684926 | 3.1M | 3.0M | 95.5% | 3.0M | 95.5% | 0.0M | 51.0bp | 49.8bp | 0.0M | 0.0M | 0.0M | 0.0M | 0.0M | 0.0M | 0.2% | 0.1% | 1.0bp | 0.0% | 1.0bp |
| SRR29684927 | 2.7M | 2.5M | 95.0% | 2.5M | 95.0% | 0.0M | 51.0bp | 49.8bp | 0.0M | 0.0M | 0.0M | 0.0M | 0.0M | 0.0M | 0.2% | 0.1% | 1.0bp | 0.0% | 1.0bp |
| SRR29684928 | 2.7M | 2.6M | 95.1% | 2.6M | 95.1% | 0.0M | 51.0bp | 49.8bp | 0.0M | 0.0M | 0.0M | 0.0M | 0.0M | 0.0M | 0.2% | 0.1% | 1.0bp | 0.0% | 1.0bp |
| SRR29684929 | 2.6M | 2.5M | 95.3% | 2.5M | 95.3% | 0.0M | 51.0bp | 49.8bp | 0.0M | 0.0M | 0.0M | 0.0M | 0.0M | 0.0M | 0.2% | 0.1% | 1.0bp | 0.0% | 1.0bp |
| SRR29684930 | 3.0M | 2.9M | 98.2% | 2.9M | 98.2% | 0.0M | 50.0bp | 49.8bp | 0.0M | 0.0M | 0.0M | 0.0M | 0.0M | 0.0M | 0.1% | 0.1% | 1.0bp | 0.0% | 1.0bp |
| SRR29684931 | 3.3M | 3.2M | 97.6% | 3.2M | 97.6% | 0.0M | 51.0bp | 49.8bp | 0.0M | 0.0M | 0.0M | 0.0M | 0.0M | 0.0M | 0.2% | 0.1% | 1.0bp | 0.0% | 1.0bp |
| SRR29684932 | 3.1M | 3.0M | 98.0% | 3.0M | 98.0% | 0.0M | 51.0bp | 49.8bp | 0.0M | 0.0M | 0.0M | 0.0M | 0.0M | 0.0M | 0.2% | 0.1% | 1.0bp | 0.0% | 1.0bp |
| SRR29684933 | 2.1M | 2.0M | 95.7% | 2.0M | 95.7% | 0.0M | 51.0bp | 49.8bp | 0.0M | 0.0M | 0.0M | 0.0M | 0.0M | 0.0M | 0.2% | 0.1% | 1.0bp | 0.0% | 1.0bp |
| SRR29684934 | 2.8M | 2.7M | 95.0% | 2.7M | 95.0% | 0.0M | 51.0bp | 49.8bp | 0.0M | 0.0M | 0.0M | 0.0M | 0.0M | 0.0M | 0.2% | 0.1% | 1.0bp | 0.0% | 1.0bp |
| SRR29684935 | 2.5M | 2.3M | 95.4% | 2.3M | 95.4% | 0.0M | 51.0bp | 49.8bp | 0.0M | 0.0M | 0.0M | 0.0M | 0.0M | 0.0M | 0.2% | 0.1% | 1.0bp | 0.0% | 1.0bp |
| SRR29684936 | 2.1M | 2.0M | 93.8% | 2.0M | 93.8% | 0.0M | 51.0bp | 49.8bp | 0.0M | 0.0M | 0.0M | 0.0M | 0.0M | 0.0M | 0.2% | 0.1% | 1.0bp | 0.0% | 1.0bp |
| SRR29684937 | 2.6M | 2.5M | 96.5% | 2.5M | 96.5% | 0.0M | 51.0bp | 49.8bp | 0.0M | 0.0M | 0.0M | 0.0M | 0.0M | 0.0M | 0.2% | 0.1% | 1.0bp | 0.0% | 1.0bp |
| SRR29684938 | 3.9M | 3.8M | 97.5% | 3.8M | 97.5% | 0.0M | 51.0bp | 49.8bp | 0.0M | 0.0M | 0.0M | 0.0M | 0.0M | 0.0M | 0.1% | 0.1% | 1.0bp | 0.0% | 1.0bp |
| SRR29684939 | 3.9M | 3.8M | 97.9% | 3.8M | 97.9% | 0.0M | 51.0bp | 49.8bp | 0.0M | 0.0M | 0.0M | 0.0M | 0.0M | 0.0M | 0.1% | 0.1% | 1.0bp | 0.0% | 1.0bp |
| SRR29684940 | 3.4M | 3.3M | 97.5% | 3.3M | 97.5% | 0.0M | 50.0bp | 49.8bp | 0.0M | 0.0M | 0.0M | 0.0M | 0.0M | 0.0M | 0.1% | 0.1% | 1.0bp | 0.0% | 1.0bp |
| SRR29684941 | 3.2M | 3.1M | 97.5% | 3.1M | 97.5% | 0.0M | 51.0bp | 49.8bp | 0.0M | 0.0M | 0.0M | 0.0M | 0.0M | 0.0M | 0.1% | 0.1% | 1.0bp | 0.0% | 1.0bp |
| SRR29684942 | 4.9M | 4.8M | 97.8% | 4.8M | 97.8% | 0.0M | 50.0bp | 49.8bp | 0.0M | 0.0M | 0.0M | 0.0M | 0.0M | 0.0M | 0.1% | 0.1% | 1.0bp | 0.0% | 1.0bp |
| SRR29684943 | 0.8M | 0.8M | 97.8% | 0.8M | 97.8% | 0.0M | 50.0bp | 49.8bp | 0.0M | 0.0M | 0.0M | 0.0M | 0.0M | 0.0M | 0.1% | 0.1% | 1.0bp | 0.0% | 1.0bp |
| SRR29684944 | 4.7M | 4.6M | 98.3% | 4.6M | 98.3% | 0.0M | 50.0bp | 49.8bp | 0.0M | 0.0M | 0.0M | 0.0M | 0.0M | 0.0M | 0.1% | 0.1% | 1.0bp | 0.0% | 1.0bp |
| SRR29684945 | 3.9M | 3.9M | 98.2% | 3.9M | 98.2% | 0.0M | 50.0bp | 49.8bp | 0.0M | 0.0M | 0.0M | 0.0M | 0.0M | 0.0M | 0.1% | 0.1% | 1.0bp | 0.0% | 1.0bp |
| SRR29684946 | 4.2M | 4.1M | 98.2% | 4.1M | 98.2% | 0.0M | 50.0bp | 49.8bp | 0.0M | 0.0M | 0.0M | 0.0M | 0.0M | 0.0M | 0.1% | 0.1% | 1.0bp | 0.0% | 1.0bp |
| SRR29684947 | 3.1M | 3.0M | 98.1% | 3.0M | 98.1% | 0.0M | 50.0bp | 49.8bp | 0.0M | 0.0M | 0.0M | 0.0M | 0.0M | 0.0M | 0.1% | 0.1% | 1.0bp | 0.0% | 1.0bp |
| SRR29684948 | 4.8M | 4.8M | 98.0% | 4.8M | 98.0% | 0.0M | 51.0bp | 49.8bp | 0.0M | 0.0M | 0.0M | 0.0M | 0.0M | 0.0M | 0.1% | 0.1% | 1.0bp | 0.0% | 1.0bp |
| SRR29684949 | 5.3M | 5.2M | 98.5% | 5.2M | 98.5% | 0.0M | 50.0bp | 49.7bp | 0.0M | 0.0M | 0.0M | 0.0M | 0.0M | 0.0M | 0.1% | 0.1% | 1.0bp | 0.0% | 1.0bp |
| SRR29684950 | 7.3M | 7.1M | 97.0% | 7.1M | 97.0% | 0.0M | 51.0bp | 49.8bp | 0.0M | 0.0M | 0.0M | 0.0M | 0.0M | 0.0M | 0.1% | 0.1% | 1.0bp | 0.0% | 1.0bp |
| SRR29684951 | 5.8M | 5.6M | 97.2% | 5.6M | 97.2% | 0.0M | 51.0bp | 49.8bp | 0.0M | 0.0M | 0.0M | 0.0M | 0.0M | 0.0M | 0.1% | 0.1% | 1.0bp | 0.0% | 1.0bp |
| SRR29684952 | 6.6M | 6.4M | 97.6% | 6.4M | 97.6% | 0.0M | 51.0bp | 49.8bp | 0.0M | 0.0M | 0.0M | 0.0M | 0.0M | 0.0M | 0.1% | 0.1% | 1.0bp | 0.0% | 1.0bp |
| SRR29684953 | 6.5M | 6.3M | 97.3% | 6.3M | 97.3% | 0.0M | 51.0bp | 49.8bp | 0.0M | 0.0M | 0.0M | 0.0M | 0.0M | 0.0M | 0.1% | 0.1% | 1.0bp | 0.0% | 1.0bp |
| SRR29684954 | 5.9M | 5.8M | 97.8% | 5.8M | 97.8% | 0.0M | 51.0bp | 49.8bp | 0.0M | 0.0M | 0.0M | 0.0M | 0.0M | 0.0M | 0.1% | 0.1% | 1.0bp | 0.0% | 1.0bp |
| SRR29684955 | 4.0M | 3.9M | 97.5% | 3.9M | 97.5% | 0.0M | 51.0bp | 49.8bp | 0.0M | 0.0M | 0.0M | 0.0M | 0.0M | 0.0M | 0.1% | 0.1% | 1.0bp | 0.0% | 1.0bp |
| SRR29684956 | 3.0M | 2.9M | 97.0% | 2.9M | 97.0% | 0.0M | 51.0bp | 49.8bp | 0.0M | 0.0M | 0.0M | 0.0M | 0.0M | 0.0M | 0.2% | 0.1% | 1.0bp | 0.0% | 1.0bp |
| SRR29684957 | 2.6M | 2.5M | 96.7% | 2.5M | 96.7% | 0.0M | 51.0bp | 49.8bp | 0.0M | 0.0M | 0.0M | 0.0M | 0.0M | 0.0M | 0.2% | 0.1% | 1.0bp | 0.0% | 1.0bp |
| SRR29684958 | 2.6M | 2.6M | 96.9% | 2.6M | 96.9% | 0.0M | 51.0bp | 49.8bp | 0.0M | 0.0M | 0.0M | 0.0M | 0.0M | 0.0M | 0.2% | 0.1% | 1.0bp | 0.0% | 1.0bp |
| SRR29684959 | 1.3M | 1.1M | 80.3% | 1.1M | 80.3% | 0.0M | 51.0bp | 49.8bp | 0.0M | 0.0M | 0.0M | 0.0M | 0.0M | 0.0M | 0.2% | 0.1% | 1.0bp | 0.0% | 1.0bp |
| SRR29684960 | 0.9M | 0.8M | 80.8% | 0.8M | 80.8% | 0.0M | 51.0bp | 49.8bp | 0.0M | 0.0M | 0.0M | 0.0M | 0.0M | 0.0M | 0.2% | 0.1% | 1.0bp | 0.0% | 1.0bp |
| SRR29684961 | 1.0M | 0.7M | 76.1% | 0.7M | 76.1% | 0.0M | 51.0bp | 49.8bp | 0.0M | 0.0M | 0.0M | 0.0M | 0.0M | 0.0M | 0.2% | 0.1% | 1.0bp | 0.0% | 1.0bp |
| SRR29684962 | 2.6M | 2.5M | 96.6% | 2.5M | 96.6% | 0.0M | 51.0bp | 49.8bp | 0.0M | 0.0M | 0.0M | 0.0M | 0.0M | 0.0M | 0.2% | 0.1% | 1.0bp | 0.0% | 1.0bp |
| SRR29684963 | 2.5M | 2.4M | 96.8% | 2.4M | 96.8% | 0.0M | 51.0bp | 49.8bp | 0.0M | 0.0M | 0.0M | 0.0M | 0.0M | 0.0M | 0.1% | 0.1% | 1.0bp | 0.0% | 1.0bp |
| SRR29684964 | 2.5M | 2.5M | 96.9% | 2.5M | 96.9% | 0.0M | 51.0bp | 49.8bp | 0.0M | 0.0M | 0.0M | 0.0M | 0.0M | 0.0M | 0.2% | 0.1% | 1.0bp | 0.0% | 1.0bp |
| SRR29684965 | 2.4M | 2.3M | 97.5% | 2.3M | 97.5% | 0.0M | 51.0bp | 49.8bp | 0.0M | 0.0M | 0.0M | 0.0M | 0.0M | 0.0M | 0.2% | 0.1% | 1.0bp | 0.0% | 1.0bp |
| SRR29684966 | 1.5M | 1.4M | 93.6% | 1.4M | 93.6% | 0.0M | 51.0bp | 49.8bp | 0.0M | 0.0M | 0.0M | 0.0M | 0.0M | 0.0M | 0.2% | 0.1% | 1.0bp | 0.0% | 1.0bp |
| SRR29684967 | 2.4M | 2.2M | 92.8% | 2.2M | 92.8% | 0.0M | 51.0bp | 49.8bp | 0.0M | 0.0M | 0.0M | 0.0M | 0.0M | 0.0M | 0.2% | 0.1% | 1.0bp | 0.0% | 1.0bp |
| SRR29684968 | 1.8M | 1.7M | 92.8% | 1.7M | 92.8% | 0.0M | 51.0bp | 49.8bp | 0.0M | 0.0M | 0.0M | 0.0M | 0.0M | 0.0M | 0.2% | 0.1% | 1.0bp | 0.0% | 1.0bp |
| SRR29684969 | 1.8M | 1.7M | 92.3% | 1.7M | 92.3% | 0.0M | 51.0bp | 49.8bp | 0.0M | 0.0M | 0.0M | 0.0M | 0.0M | 0.0M | 0.2% | 0.1% | 1.0bp | 0.0% | 1.0bp |
| SRR29684970 | 1.2M | 1.1M | 91.3% | 1.1M | 91.3% | 0.0M | 51.0bp | 49.8bp | 0.0M | 0.0M | 0.0M | 0.0M | 0.0M | 0.0M | 0.2% | 0.1% | 1.0bp | 0.0% | 1.0bp |
| SRR29684971 | 3.6M | 3.5M | 97.2% | 3.5M | 97.2% | 0.0M | 51.0bp | 49.8bp | 0.0M | 0.0M | 0.0M | 0.0M | 0.0M | 0.0M | 0.2% | 0.1% | 1.0bp | 0.0% | 1.0bp |
| SRR29684972 | 3.4M | 3.3M | 97.7% | 3.3M | 97.7% | 0.0M | 51.0bp | 49.8bp | 0.0M | 0.0M | 0.0M | 0.0M | 0.0M | 0.0M | 0.1% | 0.1% | 1.0bp | 0.0% | 1.0bp |
| SRR29684973 | 2.3M | 2.2M | 95.9% | 2.2M | 95.9% | 0.0M | 51.0bp | 49.8bp | 0.0M | 0.0M | 0.0M | 0.0M | 0.0M | 0.0M | 0.1% | 0.1% | 1.0bp | 0.0% | 1.0bp |
| SRR29684974 | 4.4M | 4.3M | 97.6% | 4.3M | 97.6% | 0.0M | 51.0bp | 49.8bp | 0.0M | 0.0M | 0.0M | 0.0M | 0.0M | 0.0M | 0.1% | 0.1% | 1.0bp | 0.0% | 1.0bp |
| SRR29684975 | 3.4M | 3.3M | 97.4% | 3.3M | 97.4% | 0.0M | 51.0bp | 49.8bp | 0.0M | 0.0M | 0.0M | 0.0M | 0.0M | 0.0M | 0.1% | 0.1% | 1.0bp | 0.0% | 1.0bp |
| SRR29684976 | 2.3M | 2.1M | 91.1% | 2.1M | 91.1% | 0.0M | 51.0bp | 49.8bp | 0.0M | 0.0M | 0.0M | 0.0M | 0.0M | 0.0M | 0.2% | 0.1% | 1.0bp | 0.0% | 1.0bp |
| SRR29684977 | 3.6M | 3.5M | 97.6% | 3.5M | 97.6% | 0.0M | 50.0bp | 49.7bp | 0.0M | 0.0M | 0.0M | 0.0M | 0.0M | 0.0M | 0.1% | 0.1% | 1.0bp | 0.0% | 1.0bp |
| SRR29684978 | 2.0M | 1.9M | 94.0% | 1.9M | 94.0% | 0.0M | 51.0bp | 49.8bp | 0.0M | 0.0M | 0.0M | 0.0M | 0.0M | 0.0M | 0.2% | 0.1% | 1.0bp | 0.0% | 1.0bp |
| SRR29684979 | 3.6M | 3.5M | 97.8% | 3.5M | 97.8% | 0.0M | 51.0bp | 49.8bp | 0.0M | 0.0M | 0.0M | 0.0M | 0.0M | 0.0M | 0.2% | 0.1% | 1.0bp | 0.0% | 1.0bp |
| SRR29684980 | 3.2M | 3.1M | 97.8% | 3.1M | 97.8% | 0.0M | 51.0bp | 49.8bp | 0.0M | 0.0M | 0.0M | 0.0M | 0.0M | 0.0M | 0.2% | 0.1% | 1.0bp | 0.0% | 1.0bp |
| SRR29684981 | 3.4M | 3.3M | 97.0% | 3.3M | 97.0% | 0.0M | 51.0bp | 49.8bp | 0.0M | 0.0M | 0.0M | 0.0M | 0.0M | 0.0M | 0.2% | 0.1% | 1.0bp | 0.0% | 1.0bp |
| SRR29684982 | 3.6M | 3.5M | 97.2% | 3.5M | 97.2% | 0.0M | 51.0bp | 49.8bp | 0.0M | 0.0M | 0.0M | 0.0M | 0.0M | 0.0M | 0.2% | 0.1% | 1.0bp | 0.0% | 1.0bp |
| SRR29684983 | 3.4M | 3.3M | 97.4% | 3.3M | 97.4% | 0.0M | 51.0bp | 49.8bp | 0.0M | 0.0M | 0.0M | 0.0M | 0.0M | 0.0M | 0.2% | 0.1% | 1.0bp | 0.0% | 1.0bp |
| SRR29684984 | 3.5M | 3.4M | 97.3% | 3.4M | 97.3% | 0.0M | 51.0bp | 49.8bp | 0.0M | 0.0M | 0.0M | 0.0M | 0.0M | 0.0M | 0.2% | 0.1% | 1.0bp | 0.0% | 1.0bp |
| SRR29684985 | 3.8M | 3.7M | 97.8% | 3.7M | 97.8% | 0.0M | 51.0bp | 49.8bp | 0.0M | 0.0M | 0.0M | 0.0M | 0.0M | 0.0M | 0.2% | 0.1% | 1.0bp | 0.0% | 1.0bp |
| SRR29684986 | 3.5M | 3.4M | 97.7% | 3.4M | 97.7% | 0.0M | 51.0bp | 49.8bp | 0.0M | 0.0M | 0.0M | 0.0M | 0.0M | 0.0M | 0.1% | 0.1% | 1.0bp | 0.0% | 1.0bp |
| SRR29684987 | 4.0M | 3.9M | 97.6% | 3.9M | 97.6% | 0.0M | 51.0bp | 49.8bp | 0.0M | 0.0M | 0.0M | 0.0M | 0.0M | 0.0M | 0.1% | 0.1% | 1.0bp | 0.0% | 1.0bp |
| SRR29684988 | 3.1M | 2.6M | 82.6% | 2.6M | 82.6% | 0.0M | 51.0bp | 49.8bp | 0.0M | 0.0M | 0.0M | 0.0M | 0.0M | 0.0M | 0.2% | 0.1% | 1.0bp | 0.0% | 1.0bp |
| SRR29684989 | 5.9M | 5.7M | 97.4% | 5.7M | 97.4% | 0.0M | 51.0bp | 49.8bp | 0.0M | 0.0M | 0.0M | 0.0M | 0.0M | 0.0M | 0.1% | 0.1% | 1.0bp | 0.0% | 1.0bp |
| SRR29684990 | 4.7M | 4.6M | 97.6% | 4.6M | 97.6% | 0.0M | 51.0bp | 49.8bp | 0.0M | 0.0M | 0.0M | 0.0M | 0.0M | 0.0M | 0.1% | 0.1% | 1.0bp | 0.0% | 1.0bp |
| SRR29684991 | 4.0M | 3.9M | 97.8% | 3.9M | 97.8% | 0.0M | 51.0bp | 49.8bp | 0.0M | 0.0M | 0.0M | 0.0M | 0.0M | 0.0M | 0.1% | 0.1% | 1.0bp | 0.0% | 1.0bp |
| SRR29684992 | 5.0M | 4.8M | 97.2% | 4.8M | 97.2% | 0.0M | 51.0bp | 49.8bp | 0.0M | 0.0M | 0.0M | 0.0M | 0.0M | 0.0M | 0.1% | 0.1% | 1.0bp | 0.0% | 1.0bp |
| SRR29684993 | 4.9M | 4.8M | 97.7% | 4.8M | 97.7% | 0.0M | 51.0bp | 49.8bp | 0.0M | 0.0M | 0.0M | 0.0M | 0.0M | 0.0M | 0.1% | 0.1% | 1.0bp | 0.0% | 1.0bp |
| SRR29684994 | 3.3M | 3.2M | 98.2% | 3.2M | 98.2% | 0.0M | 50.0bp | 49.7bp | 0.0M | 0.0M | 0.0M | 0.0M | 0.0M | 0.0M | 0.1% | 0.1% | 1.0bp | 0.0% | 1.0bp |
| SRR29684995 | 3.8M | 3.7M | 97.3% | 3.7M | 97.3% | 0.0M | 51.0bp | 49.8bp | 0.0M | 0.0M | 0.0M | 0.0M | 0.0M | 0.0M | 0.1% | 0.1% | 1.0bp | 0.0% | 1.0bp |
| SRR29684996 | 4.0M | 3.9M | 97.1% | 3.9M | 97.1% | 0.0M | 51.0bp | 49.8bp | 0.0M | 0.0M | 0.0M | 0.0M | 0.0M | 0.0M | 0.1% | 0.1% | 1.0bp | 0.0% | 1.0bp |
| SRR29684997 | 5.3M | 5.2M | 97.5% | 5.2M | 97.5% | 0.0M | 51.0bp | 49.8bp | 0.0M | 0.0M | 0.0M | 0.0M | 0.0M | 0.0M | 0.1% | 0.1% | 1.0bp | 0.0% | 1.0bp |
| SRR29684998 | 3.9M | 3.8M | 97.2% | 3.8M | 97.2% | 0.0M | 51.0bp | 49.8bp | 0.0M | 0.0M | 0.0M | 0.0M | 0.0M | 0.0M | 0.1% | 0.1% | 1.0bp | 0.0% | 1.0bp |
| SRR29684999 | 4.7M | 4.6M | 97.5% | 4.6M | 97.5% | 0.0M | 51.0bp | 49.8bp | 0.0M | 0.0M | 0.0M | 0.0M | 0.0M | 0.0M | 0.1% | 0.1% | 1.0bp | 0.0% | 1.0bp |
| SRR29685000 | 4.3M | 4.2M | 97.7% | 4.2M | 97.7% | 0.0M | 51.0bp | 49.8bp | 0.0M | 0.0M | 0.0M | 0.0M | 0.0M | 0.0M | 0.1% | 0.1% | 1.0bp | 0.0% | 1.0bp |
| SRR29685001 | 4.1M | 4.0M | 97.7% | 4.0M | 97.7% | 0.0M | 51.0bp | 49.8bp | 0.0M | 0.0M | 0.0M | 0.0M | 0.0M | 0.0M | 0.1% | 0.1% | 1.0bp | 0.0% | 1.0bp |
| SRR29685002 | 4.0M | 3.9M | 97.6% | 3.9M | 97.6% | 0.0M | 51.0bp | 49.8bp | 0.0M | 0.0M | 0.0M | 0.0M | 0.0M | 0.0M | 0.1% | 0.1% | 1.0bp | 0.0% | 1.0bp |
| SRR29685003 | 4.1M | 4.0M | 97.5% | 4.0M | 97.5% | 0.0M | 50.0bp | 49.8bp | 0.0M | 0.0M | 0.0M | 0.0M | 0.0M | 0.0M | 0.1% | 0.1% | 1.0bp | 0.0% | 1.0bp |
| SRR29685004 | 5.8M | 5.7M | 98.3% | 5.7M | 98.3% | 0.0M | 51.0bp | 49.8bp | 0.0M | 0.0M | 0.0M | 0.0M | 0.0M | 0.0M | 0.1% | 0.1% | 1.0bp | 0.0% | 1.0bp |
| SRR29685005 | 2.8M | 2.7M | 98.3% | 2.7M | 98.3% | 0.0M | 50.0bp | 49.8bp | 0.0M | 0.0M | 0.0M | 0.0M | 0.0M | 0.0M | 0.1% | 0.1% | 1.0bp | 0.0% | 1.0bp |
| SRR29685006 | 4.8M | 4.7M | 98.1% | 4.7M | 98.1% | 0.0M | 51.0bp | 49.8bp | 0.0M | 0.0M | 0.0M | 0.0M | 0.0M | 0.0M | 0.1% | 0.1% | 1.0bp | 0.0% | 1.0bp |
| SRR29685007 | 4.7M | 4.6M | 97.8% | 4.6M | 97.8% | 0.0M | 51.0bp | 49.8bp | 0.0M | 0.0M | 0.0M | 0.0M | 0.0M | 0.0M | 0.1% | 0.1% | 1.0bp | 0.0% | 1.0bp |
| SRR29685008 | 4.5M | 4.4M | 98.0% | 4.4M | 98.0% | 0.0M | 50.0bp | 49.8bp | 0.0M | 0.0M | 0.0M | 0.0M | 0.0M | 0.0M | 0.1% | 0.1% | 1.0bp | 0.0% | 1.0bp |
| SRR29685009 | 3.5M | 3.5M | 97.6% | 3.5M | 97.6% | 0.0M | 51.0bp | 49.8bp | 0.0M | 0.0M | 0.0M | 0.0M | 0.0M | 0.0M | 0.1% | 0.1% | 1.0bp | 0.0% | 1.0bp |
| SRR29685010 | 3.3M | 3.2M | 97.8% | 3.2M | 97.8% | 0.0M | 51.0bp | 49.8bp | 0.0M | 0.0M | 0.0M | 0.0M | 0.0M | 0.0M | 0.1% | 0.1% | 1.0bp | 0.0% | 1.0bp |
| SRR29685011 | 4.3M | 4.2M | 98.3% | 4.2M | 98.3% | 0.0M | 50.0bp | 49.7bp | 0.0M | 0.0M | 0.0M | 0.0M | 0.0M | 0.0M | 0.1% | 0.1% | 1.0bp | 0.0% | 1.0bp |
| SRR29685012 | 6.2M | 6.0M | 97.3% | 6.0M | 97.3% | 0.0M | 51.0bp | 49.8bp | 0.0M | 0.0M | 0.0M | 0.0M | 0.0M | 0.0M | 0.1% | 0.1% | 1.0bp | 0.0% | 1.0bp |
| SRR29685013 | 5.4M | 5.3M | 98.1% | 5.3M | 98.1% | 0.0M | 51.0bp | 49.8bp | 0.0M | 0.0M | 0.0M | 0.0M | 0.0M | 0.0M | 0.1% | 0.1% | 1.0bp | 0.0% | 1.0bp |
| SRR29685014 | 5.6M | 5.5M | 97.9% | 5.5M | 97.9% | 0.0M | 51.0bp | 49.8bp | 0.0M | 0.0M | 0.0M | 0.0M | 0.0M | 0.0M | 0.1% | 0.1% | 1.0bp | 0.0% | 1.0bp |
| SRR29685015 | 5.1M | 4.9M | 97.1% | 4.9M | 97.1% | 0.0M | 51.0bp | 49.8bp | 0.0M | 0.0M | 0.0M | 0.0M | 0.0M | 0.0M | 0.1% | 0.1% | 1.0bp | 0.0% | 1.0bp |
| SRR29685016 | 4.8M | 4.7M | 98.0% | 4.7M | 98.0% | 0.0M | 50.0bp | 49.8bp | 0.0M | 0.0M | 0.0M | 0.0M | 0.0M | 0.0M | 0.1% | 0.1% | 1.0bp | 0.0% | 1.0bp |
| SRR29685017 | 4.3M | 4.2M | 97.7% | 4.2M | 97.7% | 0.0M | 50.0bp | 49.8bp | 0.0M | 0.0M | 0.0M | 0.0M | 0.0M | 0.0M | 0.1% | 0.1% | 1.0bp | 0.0% | 1.0bp |
| SRR29685018 | 3.8M | 3.7M | 96.8% | 3.7M | 96.8% | 0.0M | 51.0bp | 49.8bp | 0.0M | 0.0M | 0.0M | 0.0M | 0.0M | 0.0M | 0.1% | 0.1% | 1.0bp | 0.0% | 1.0bp |
| SRR29685019 | 5.5M | 5.4M | 98.7% | 5.4M | 98.7% | 0.0M | 50.0bp | 49.7bp | 0.0M | 0.0M | 0.0M | 0.0M | 0.0M | 0.0M | 0.1% | 0.1% | 1.0bp | 0.0% | 1.0bp |
| SRR29685020 | 6.8M | 6.6M | 96.1% | 6.6M | 96.1% | 0.0M | 51.0bp | 49.8bp | 0.0M | 0.0M | 0.0M | 0.0M | 0.0M | 0.0M | 0.1% | 0.1% | 1.0bp | 0.0% | 1.0bp |
| SRR29685021 | 4.4M | 4.3M | 98.1% | 4.3M | 98.1% | 0.0M | 51.0bp | 49.8bp | 0.0M | 0.0M | 0.0M | 0.0M | 0.0M | 0.0M | 0.1% | 0.1% | 1.0bp | 0.0% | 1.0bp |
| SRR29685022 | 5.8M | 5.7M | 98.3% | 5.7M | 98.3% | 0.0M | 51.0bp | 49.8bp | 0.0M | 0.0M | 0.0M | 0.0M | 0.0M | 0.0M | 0.1% | 0.1% | 1.0bp | 0.0% | 1.0bp |
| SRR29685023 | 6.3M | 6.2M | 98.3% | 6.2M | 98.3% | 0.0M | 51.0bp | 49.8bp | 0.0M | 0.0M | 0.0M | 0.0M | 0.0M | 0.0M | 0.1% | 0.1% | 1.0bp | 0.0% | 1.0bp |
| SRR29685024 | 4.7M | 4.6M | 98.2% | 4.6M | 98.2% | 0.0M | 51.0bp | 49.8bp | 0.0M | 0.0M | 0.0M | 0.0M | 0.0M | 0.0M | 0.1% | 0.1% | 1.0bp | 0.0% | 1.0bp |
| SRR29685025 | 4.3M | 4.2M | 97.5% | 4.2M | 97.5% | 0.0M | 51.0bp | 49.8bp | 0.0M | 0.0M | 0.0M | 0.0M | 0.0M | 0.0M | 0.1% | 0.1% | 1.0bp | 0.0% | 1.0bp |
| SRR29685026 | 3.6M | 3.5M | 97.3% | 3.5M | 97.3% | 0.0M | 51.0bp | 49.8bp | 0.0M | 0.0M | 0.0M | 0.0M | 0.0M | 0.0M | 0.1% | 0.1% | 1.0bp | 0.0% | 1.0bp |
| SRR29685027 | 4.3M | 4.2M | 96.8% | 4.2M | 96.8% | 0.0M | 51.0bp | 49.8bp | 0.0M | 0.0M | 0.0M | 0.0M | 0.0M | 0.0M | 0.1% | 0.1% | 1.0bp | 0.0% | 1.0bp |
| SRR29685028 | 3.3M | 2.4M | 73.1% | 2.4M | 73.1% | 0.0M | 51.0bp | 49.8bp | 0.0M | 0.0M | 0.0M | 0.0M | 0.0M | 0.0M | 0.2% | 0.1% | 1.0bp | 0.0% | 1.0bp |
| SRR29685029 | 3.9M | 3.8M | 97.0% | 3.8M | 97.0% | 0.0M | 51.0bp | 49.8bp | 0.0M | 0.0M | 0.0M | 0.0M | 0.0M | 0.0M | 0.1% | 0.1% | 1.0bp | 0.0% | 1.0bp |
| SRR29685030 | 4.6M | 4.5M | 97.0% | 4.5M | 97.0% | 0.0M | 51.0bp | 49.8bp | 0.0M | 0.0M | 0.0M | 0.0M | 0.0M | 0.0M | 0.1% | 0.1% | 1.0bp | 0.0% | 1.0bp |
| SRR29685031 | 5.0M | 4.9M | 97.4% | 4.9M | 97.4% | 0.0M | 51.0bp | 49.8bp | 0.0M | 0.0M | 0.0M | 0.0M | 0.0M | 0.0M | 0.1% | 0.1% | 1.0bp | 0.0% | 1.0bp |
| SRR29685032 | 4.5M | 4.4M | 97.5% | 4.4M | 97.5% | 0.0M | 51.0bp | 49.8bp | 0.0M | 0.0M | 0.0M | 0.0M | 0.0M | 0.0M | 0.1% | 0.1% | 1.0bp | 0.0% | 1.0bp |
| SRR29685033 | 3.1M | 3.0M | 97.4% | 3.0M | 97.4% | 0.0M | 51.0bp | 49.8bp | 0.0M | 0.0M | 0.0M | 0.0M | 0.0M | 0.0M | 0.1% | 0.1% | 1.0bp | 0.0% | 1.0bp |
| SRR29685034 | 3.8M | 3.7M | 98.6% | 3.7M | 98.6% | 0.0M | 50.0bp | 49.7bp | 0.0M | 0.0M | 0.0M | 0.0M | 0.0M | 0.0M | 0.1% | 0.1% | 1.0bp | 0.0% | 1.0bp |
| SRR29685035 | 3.5M | 3.4M | 96.6% | 3.4M | 96.6% | 0.0M | 51.0bp | 49.8bp | 0.0M | 0.0M | 0.0M | 0.0M | 0.0M | 0.0M | 0.1% | 0.1% | 1.0bp | 0.0% | 1.0bp |
| SRR29685036 | 4.4M | 4.2M | 95.3% | 4.2M | 95.3% | 0.0M | 51.0bp | 49.8bp | 0.0M | 0.0M | 0.0M | 0.0M | 0.0M | 0.0M | 0.1% | 0.1% | 1.0bp | 0.0% | 1.0bp |
| SRR29685037 | 3.5M | 3.4M | 96.7% | 3.4M | 96.7% | 0.0M | 51.0bp | 49.8bp | 0.0M | 0.0M | 0.0M | 0.0M | 0.0M | 0.0M | 0.1% | 0.1% | 1.0bp | 0.0% | 1.0bp |
| SRR29685038 | 3.7M | 3.6M | 97.0% | 3.6M | 97.0% | 0.0M | 51.0bp | 49.8bp | 0.0M | 0.0M | 0.0M | 0.0M | 0.0M | 0.0M | 0.1% | 0.1% | 1.0bp | 0.0% | 1.0bp |
| SRR29685039 | 3.8M | 3.7M | 96.9% | 3.7M | 96.9% | 0.0M | 51.0bp | 49.8bp | 0.0M | 0.0M | 0.0M | 0.0M | 0.0M | 0.0M | 0.1% | 0.1% | 1.0bp | 0.0% | 1.0bp |
| SRR29685040 | 4.1M | 4.0M | 97.0% | 4.0M | 97.0% | 0.0M | 51.0bp | 49.8bp | 0.0M | 0.0M | 0.0M | 0.0M | 0.0M | 0.0M | 0.1% | 0.1% | 1.0bp | 0.0% | 1.0bp |
| SRR29685041 | 3.9M | 3.8M | 97.6% | 3.8M | 97.6% | 0.0M | 51.0bp | 49.8bp | 0.0M | 0.0M | 0.0M | 0.0M | 0.0M | 0.0M | 0.1% | 0.1% | 1.0bp | 0.0% | 1.0bp |
| SRR29685042 | 4.2M | 4.1M | 97.3% | 4.1M | 97.3% | 0.0M | 51.0bp | 49.8bp | 0.0M | 0.0M | 0.0M | 0.0M | 0.0M | 0.0M | 0.1% | 0.1% | 1.0bp | 0.0% | 1.0bp |
| SRR29685043 | 3.7M | 3.6M | 96.3% | 3.6M | 96.3% | 0.0M | 51.0bp | 49.8bp | 0.0M | 0.0M | 0.0M | 0.0M | 0.0M | 0.0M | 0.1% | 0.1% | 1.0bp | 0.0% | 1.0bp |
| SRR29685044 | 5.4M | 5.3M | 97.1% | 5.3M | 97.1% | 0.0M | 51.0bp | 49.8bp | 0.0M | 0.0M | 0.0M | 0.0M | 0.0M | 0.0M | 0.1% | 0.1% | 1.0bp | 0.0% | 1.0bp |
| SRR29685045 | 4.1M | 4.0M | 96.8% | 4.0M | 96.8% | 0.0M | 51.0bp | 49.8bp | 0.0M | 0.0M | 0.0M | 0.0M | 0.0M | 0.0M | 0.1% | 0.1% | 1.0bp | 0.0% | 1.0bp |
| SRR29685046 | 4.2M | 4.1M | 96.9% | 4.1M | 96.9% | 0.0M | 51.0bp | 49.8bp | 0.0M | 0.0M | 0.0M | 0.0M | 0.0M | 0.0M | 0.1% | 0.1% | 1.0bp | 0.0% | 1.0bp |
| SRR29685047 | 3.9M | 3.8M | 96.7% | 3.8M | 96.7% | 0.0M | 51.0bp | 49.8bp | 0.0M | 0.0M | 0.0M | 0.0M | 0.0M | 0.0M | 0.1% | 0.1% | 1.0bp | 0.0% | 1.0bp |
| SRR29685048 | 4.1M | 4.0M | 96.9% | 4.0M | 96.9% | 0.0M | 51.0bp | 49.8bp | 0.0M | 0.0M | 0.0M | 0.0M | 0.0M | 0.0M | 0.1% | 0.1% | 1.0bp | 0.0% | 1.0bp |
| SRR29685049 | 4.0M | 3.9M | 98.3% | 3.9M | 98.3% | 0.0M | 51.0bp | 49.8bp | 0.0M | 0.0M | 0.0M | 0.0M | 0.0M | 0.0M | 0.1% | 0.1% | 1.0bp | 0.0% | 1.0bp |
| SRR29685050 | 3.7M | 3.5M | 96.7% | 3.5M | 96.7% | 0.0M | 51.0bp | 49.8bp | 0.0M | 0.0M | 0.0M | 0.0M | 0.0M | 0.0M | 0.1% | 0.1% | 1.0bp | 0.0% | 1.0bp |
| SRR29685051 | 5.1M | 4.9M | 96.0% | 4.9M | 96.0% | 0.0M | 51.0bp | 49.8bp | 0.0M | 0.0M | 0.0M | 0.0M | 0.0M | 0.0M | 0.1% | 0.1% | 1.0bp | 0.0% | 1.0bp |
| SRR29685052 | 2.7M | 2.6M | 97.5% | 2.6M | 97.5% | 0.0M | 50.0bp | 49.8bp | 0.0M | 0.0M | 0.0M | 0.0M | 0.0M | 0.0M | 0.1% | 0.1% | 1.0bp | 0.0% | 1.0bp |
| SRR29685053 | 4.2M | 4.1M | 97.3% | 4.1M | 97.3% | 0.0M | 51.0bp | 49.8bp | 0.0M | 0.0M | 0.0M | 0.0M | 0.0M | 0.0M | 0.1% | 0.1% | 1.0bp | 0.0% | 1.0bp |
| SRR29685054 | 4.5M | 4.4M | 97.7% | 4.4M | 97.7% | 0.0M | 51.0bp | 49.8bp | 0.0M | 0.0M | 0.0M | 0.0M | 0.0M | 0.0M | 0.1% | 0.1% | 1.0bp | 0.0% | 1.0bp |
| SRR29685055 | 3.8M | 3.8M | 97.7% | 3.8M | 97.7% | 0.0M | 51.0bp | 49.8bp | 0.0M | 0.0M | 0.0M | 0.0M | 0.0M | 0.0M | 0.1% | 0.1% | 1.0bp | 0.0% | 1.0bp |
| SRR29685056 | 3.6M | 3.5M | 97.4% | 3.5M | 97.4% | 0.0M | 51.0bp | 49.8bp | 0.0M | 0.0M | 0.0M | 0.0M | 0.0M | 0.0M | 0.1% | 0.1% | 1.0bp | 0.0% | 1.0bp |
| SRR29685057 | 3.8M | 3.7M | 98.2% | 3.7M | 98.2% | 0.0M | 50.0bp | 49.7bp | 0.0M | 0.0M | 0.0M | 0.0M | 0.0M | 0.0M | 0.1% | 0.1% | 1.0bp | 0.0% | 1.0bp |
| SRR29685058 | 3.4M | 3.3M | 96.7% | 3.3M | 96.7% | 0.0M | 51.0bp | 49.8bp | 0.0M | 0.0M | 0.0M | 0.0M | 0.0M | 0.0M | 0.1% | 0.1% | 1.0bp | 0.0% | 1.0bp |
| SRR29685059 | 4.6M | 4.4M | 97.2% | 4.4M | 97.2% | 0.0M | 51.0bp | 49.8bp | 0.0M | 0.0M | 0.0M | 0.0M | 0.0M | 0.0M | 0.1% | 0.1% | 1.0bp | 0.0% | 1.0bp |
| SRR29685060 | 3.9M | 3.8M | 97.6% | 3.8M | 97.6% | 0.0M | 51.0bp | 49.8bp | 0.0M | 0.0M | 0.0M | 0.0M | 0.0M | 0.0M | 0.1% | 0.1% | 1.0bp | 0.0% | 1.0bp |
| SRR29685061 | 4.1M | 4.0M | 97.5% | 4.0M | 97.5% | 0.0M | 51.0bp | 49.8bp | 0.0M | 0.0M | 0.0M | 0.0M | 0.0M | 0.0M | 0.1% | 0.1% | 1.0bp | 0.0% | 1.0bp |
| SRR29685062 | 4.2M | 4.1M | 97.8% | 4.1M | 97.8% | 0.0M | 51.0bp | 49.8bp | 0.0M | 0.0M | 0.0M | 0.0M | 0.0M | 0.0M | 0.1% | 0.1% | 1.0bp | 0.0% | 1.0bp |
| SRR29685063 | 3.7M | 3.6M | 97.5% | 3.6M | 97.5% | 0.0M | 51.0bp | 49.8bp | 0.0M | 0.0M | 0.0M | 0.0M | 0.0M | 0.0M | 0.1% | 0.1% | 1.0bp | 0.0% | 1.0bp |
| SRR29685064 | 2.8M | 2.7M | 96.6% | 2.7M | 96.6% | 0.0M | 51.0bp | 49.8bp | 0.0M | 0.0M | 0.0M | 0.0M | 0.0M | 0.0M | 0.1% | 0.1% | 1.0bp | 0.0% | 1.0bp |
| SRR29685065 | 3.6M | 3.5M | 97.5% | 3.5M | 97.5% | 0.0M | 51.0bp | 49.8bp | 0.0M | 0.0M | 0.0M | 0.0M | 0.0M | 0.0M | 0.1% | 0.1% | 1.0bp | 0.0% | 1.0bp |
| SRR29685066 | 3.6M | 3.6M | 98.7% | 3.6M | 98.7% | 0.0M | 50.0bp | 49.7bp | 0.0M | 0.0M | 0.0M | 0.0M | 0.0M | 0.0M | 0.1% | 0.1% | 1.0bp | 0.0% | 1.0bp |
| SRR29685067 | 5.1M | 5.0M | 97.4% | 5.0M | 97.4% | 0.0M | 51.0bp | 49.8bp | 0.0M | 0.0M | 0.0M | 0.0M | 0.0M | 0.0M | 0.1% | 0.1% | 1.0bp | 0.0% | 1.0bp |
| SRR29685068 | 4.1M | 4.0M | 97.8% | 4.0M | 97.8% | 0.0M | 51.0bp | 49.8bp | 0.0M | 0.0M | 0.0M | 0.0M | 0.0M | 0.0M | 0.1% | 0.1% | 1.0bp | 0.0% | 1.0bp |
| SRR29685069 | 2.7M | 2.6M | 97.0% | 2.6M | 97.0% | 0.0M | 51.0bp | 49.8bp | 0.0M | 0.0M | 0.0M | 0.0M | 0.0M | 0.0M | 0.1% | 0.1% | 1.0bp | 0.0% | 1.0bp |
| SRR29685070 | 4.4M | 4.3M | 98.2% | 4.3M | 98.2% | 0.0M | 50.0bp | 49.7bp | 0.0M | 0.0M | 0.0M | 0.0M | 0.0M | 0.0M | 0.1% | 0.1% | 1.0bp | 0.0% | 1.0bp |
| SRR29685071 | 4.3M | 4.2M | 98.2% | 4.2M | 98.2% | 0.0M | 50.0bp | 49.7bp | 0.0M | 0.0M | 0.0M | 0.0M | 0.0M | 0.0M | 0.1% | 0.1% | 1.0bp | 0.0% | 1.0bp |
| SRR29685072 | 3.6M | 3.5M | 98.4% | 3.5M | 98.4% | 0.0M | 50.0bp | 49.7bp | 0.0M | 0.0M | 0.0M | 0.0M | 0.0M | 0.0M | 0.1% | 0.1% | 1.0bp | 0.0% | 1.0bp |
| SRR29685073 | 2.3M | 2.2M | 95.6% | 2.2M | 95.6% | 0.0M | 51.0bp | 49.8bp | 0.0M | 0.0M | 0.0M | 0.0M | 0.0M | 0.0M | 0.1% | 0.1% | 1.0bp | 0.0% | 1.0bp |
| SRR29685074 | 1.7M | 1.5M | 88.4% | 1.5M | 88.4% | 0.0M | 51.0bp | 49.8bp | 0.0M | 0.0M | 0.0M | 0.0M | 0.0M | 0.0M | 0.2% | 0.1% | 1.0bp | 0.0% | 1.0bp |
| SRR29685075 | 3.0M | 2.9M | 97.5% | 2.9M | 97.5% | 0.0M | 51.0bp | 49.8bp | 0.0M | 0.0M | 0.0M | 0.0M | 0.0M | 0.0M | 0.2% | 0.1% | 1.0bp | 0.0% | 1.0bp |
| SRR29685076 | 2.6M | 2.5M | 97.4% | 2.5M | 97.4% | 0.0M | 51.0bp | 49.8bp | 0.0M | 0.0M | 0.0M | 0.0M | 0.0M | 0.0M | 0.2% | 0.1% | 1.0bp | 0.0% | 1.0bp |
| SRR29685077 | 2.7M | 2.6M | 97.9% | 2.6M | 97.9% | 0.0M | 50.0bp | 49.7bp | 0.0M | 0.0M | 0.0M | 0.0M | 0.0M | 0.0M | 0.1% | 0.1% | 1.0bp | 0.0% | 1.0bp |
| SRR29685078 | 2.8M | 2.7M | 95.5% | 2.7M | 95.5% | 0.0M | 51.0bp | 49.8bp | 0.0M | 0.0M | 0.0M | 0.0M | 0.0M | 0.0M | 0.2% | 0.1% | 1.0bp | 0.0% | 1.0bp |
| SRR29685079 | 2.5M | 2.4M | 95.6% | 2.4M | 95.6% | 0.0M | 51.0bp | 49.8bp | 0.0M | 0.0M | 0.0M | 0.0M | 0.0M | 0.0M | 0.2% | 0.1% | 1.0bp | 0.0% | 1.0bp |
| SRR29685080 | 2.6M | 2.5M | 95.7% | 2.5M | 95.7% | 0.0M | 51.0bp | 49.8bp | 0.0M | 0.0M | 0.0M | 0.0M | 0.0M | 0.0M | 0.2% | 0.1% | 1.0bp | 0.0% | 1.0bp |
| SRR29685081 | 1.1M | 1.0M | 88.8% | 1.0M | 88.8% | 0.0M | 51.0bp | 49.8bp | 0.0M | 0.0M | 0.0M | 0.0M | 0.0M | 0.0M | 0.2% | 0.1% | 1.0bp | 0.0% | 1.0bp |
| SRR29685082 | 0.5M | 0.1M | 13.0% | 0.1M | 13.0% | 0.0M | 51.0bp | 49.8bp | 0.0M | 0.0M | 0.0M | 0.0M | 0.0M | 0.0M | 0.2% | 0.1% | 1.0bp | 0.0% | 1.0bp |
| SRR29685083 | 2.7M | 2.6M | 98.1% | 2.6M | 98.1% | 0.0M | 50.0bp | 49.7bp | 0.0M | 0.0M | 0.0M | 0.0M | 0.0M | 0.0M | 0.1% | 0.1% | 1.0bp | 0.0% | 1.0bp |
| SRR29685084 | 1.4M | 1.3M | 93.0% | 1.3M | 93.0% | 0.0M | 51.0bp | 49.8bp | 0.0M | 0.0M | 0.0M | 0.0M | 0.0M | 0.0M | 0.2% | 0.1% | 1.0bp | 0.0% | 1.0bp |
| SRR29685085 | 2.2M | 2.1M | 93.0% | 2.1M | 93.0% | 0.0M | 51.0bp | 49.8bp | 0.0M | 0.0M | 0.0M | 0.0M | 0.0M | 0.0M | 0.2% | 0.1% | 1.0bp | 0.0% | 1.0bp |
| SRR29685086 | 1.7M | 1.6M | 94.4% | 1.6M | 94.4% | 0.0M | 51.0bp | 49.8bp | 0.0M | 0.0M | 0.0M | 0.0M | 0.0M | 0.0M | 0.2% | 0.1% | 1.0bp | 0.0% | 1.0bp |
| SRR29685087 | 1.8M | 1.7M | 94.2% | 1.7M | 94.2% | 0.0M | 51.0bp | 49.8bp | 0.0M | 0.0M | 0.0M | 0.0M | 0.0M | 0.0M | 0.2% | 0.1% | 1.0bp | 0.0% | 1.0bp |
| SRR29685088 | 1.6M | 1.5M | 92.8% | 1.5M | 92.8% | 0.0M | 51.0bp | 49.8bp | 0.0M | 0.0M | 0.0M | 0.0M | 0.0M | 0.0M | 0.2% | 0.1% | 1.0bp | 0.0% | 1.0bp |
| SRR29685089 | 3.6M | 3.5M | 97.3% | 3.5M | 97.3% | 0.0M | 51.0bp | 49.8bp | 0.0M | 0.0M | 0.0M | 0.0M | 0.0M | 0.0M | 0.1% | 0.1% | 1.0bp | 0.0% | 1.0bp |
| SRR29685090 | 3.4M | 3.3M | 97.2% | 3.3M | 97.2% | 0.0M | 51.0bp | 49.8bp | 0.0M | 0.0M | 0.0M | 0.0M | 0.0M | 0.0M | 0.1% | 0.1% | 1.0bp | 0.0% | 1.0bp |
| SRR29685091 | 5.3M | 5.1M | 97.5% | 5.1M | 97.5% | 0.0M | 50.0bp | 49.8bp | 0.0M | 0.0M | 0.0M | 0.0M | 0.0M | 0.0M | 0.1% | 0.1% | 1.0bp | 0.0% | 1.0bp |
| SRR29685092 | 6.4M | 6.2M | 96.8% | 6.2M | 96.8% | 0.0M | 51.0bp | 49.8bp | 0.0M | 0.0M | 0.0M | 0.0M | 0.0M | 0.0M | 0.1% | 0.1% | 1.0bp | 0.0% | 1.0bp |
| SRR29685093 | 6.3M | 6.2M | 97.6% | 6.2M | 97.6% | 0.0M | 50.0bp | 49.8bp | 0.0M | 0.0M | 0.0M | 0.0M | 0.0M | 0.0M | 0.1% | 0.1% | 1.0bp | 0.0% | 1.0bp |
| SRR29685094 | 5.2M | 5.1M | 97.6% | 5.1M | 97.6% | 0.0M | 51.0bp | 49.8bp | 0.0M | 0.0M | 0.0M | 0.0M | 0.0M | 0.0M | 0.1% | 0.1% | 1.0bp | 0.0% | 1.0bp |
| SRR29685095 | 6.0M | 5.9M | 97.9% | 5.9M | 97.9% | 0.0M | 51.0bp | 49.8bp | 0.0M | 0.0M | 0.0M | 0.0M | 0.0M | 0.0M | 0.1% | 0.1% | 1.0bp | 0.0% | 1.0bp |
| SRR29685096 | 4.1M | 4.0M | 96.5% | 4.0M | 96.5% | 0.0M | 51.0bp | 49.8bp | 0.0M | 0.0M | 0.0M | 0.0M | 0.0M | 0.0M | 0.1% | 0.1% | 1.0bp | 0.0% | 1.0bp |
| SRR29685097 | 2.8M | 2.6M | 95.0% | 2.6M | 95.0% | 0.0M | 51.0bp | 49.8bp | 0.0M | 0.0M | 0.0M | 0.0M | 0.0M | 0.0M | 0.1% | 0.1% | 1.0bp | 0.0% | 1.0bp |
| SRR29685098 | 3.2M | 3.2M | 98.0% | 3.2M | 98.0% | 0.0M | 50.0bp | 49.8bp | 0.0M | 0.0M | 0.0M | 0.0M | 0.0M | 0.0M | 0.1% | 0.1% | 1.0bp | 0.0% | 1.0bp |
| SRR29685099 | 3.6M | 3.5M | 97.2% | 3.5M | 97.2% | 0.0M | 51.0bp | 49.8bp | 0.0M | 0.0M | 0.0M | 0.0M | 0.0M | 0.0M | 0.1% | 0.1% | 1.0bp | 0.0% | 1.0bp |
| SRR29685100 | 5.2M | 5.0M | 97.0% | 5.0M | 97.0% | 0.0M | 51.0bp | 49.8bp | 0.0M | 0.0M | 0.0M | 0.0M | 0.0M | 0.0M | 0.1% | 0.1% | 1.0bp | 0.0% | 1.0bp |
| SRR29685101 | 3.5M | 3.3M | 93.8% | 3.3M | 93.8% | 0.0M | 51.0bp | 49.8bp | 0.0M | 0.0M | 0.0M | 0.0M | 0.0M | 0.0M | 0.1% | 0.1% | 1.0bp | 0.0% | 1.0bp |
| SRR29685102 | 4.2M | 4.1M | 97.0% | 4.1M | 97.0% | 0.0M | 51.0bp | 49.8bp | 0.0M | 0.0M | 0.0M | 0.0M | 0.0M | 0.0M | 0.1% | 0.1% | 1.0bp | 0.0% | 1.0bp |
| SRR29685103 | 4.8M | 4.7M | 97.7% | 4.7M | 97.7% | 0.0M | 51.0bp | 49.8bp | 0.0M | 0.0M | 0.0M | 0.0M | 0.0M | 0.0M | 0.1% | 0.1% | 1.0bp | 0.0% | 1.0bp |
| SRR29685104 | 3.6M | 3.6M | 97.3% | 3.6M | 97.3% | 0.0M | 51.0bp | 49.8bp | 0.0M | 0.0M | 0.0M | 0.0M | 0.0M | 0.0M | 0.1% | 0.1% | 1.0bp | 0.0% | 1.0bp |
| SRR29685105 | 3.8M | 3.5M | 92.4% | 3.5M | 92.4% | 0.0M | 51.0bp | 49.8bp | 0.0M | 0.0M | 0.0M | 0.0M | 0.0M | 0.0M | 0.1% | 0.1% | 1.0bp | 0.0% | 1.0bp |
| SRR29685106 | 2.3M | 1.8M | 78.8% | 1.8M | 78.8% | 0.0M | 51.0bp | 49.8bp | 0.0M | 0.0M | 0.0M | 0.0M | 0.0M | 0.0M | 0.2% | 0.1% | 1.0bp | 0.0% | 1.0bp |
| SRR29685107 | 1.0M | 0.1M | 7.8% | 0.1M | 7.8% | 0.0M | 51.0bp | 49.7bp | 0.0M | 0.0M | 0.0M | 0.0M | 0.0M | 0.0M | 0.2% | 0.2% | 1.0bp | 0.0% | 1.1bp |
| SRR29685108 | 5.1M | 5.0M | 97.8% | 5.0M | 97.8% | 0.0M | 50.0bp | 49.8bp | 0.0M | 0.0M | 0.0M | 0.0M | 0.0M | 0.0M | 0.1% | 0.1% | 1.0bp | 0.0% | 1.0bp |
| SRR29685109 | 4.2M | 4.1M | 97.7% | 4.1M | 97.7% | 0.0M | 51.0bp | 49.8bp | 0.0M | 0.0M | 0.0M | 0.0M | 0.0M | 0.0M | 0.1% | 0.1% | 1.0bp | 0.0% | 1.0bp |
| SRR29685110 | 4.8M | 4.7M | 98.0% | 4.7M | 98.0% | 0.0M | 50.0bp | 49.7bp | 0.0M | 0.0M | 0.0M | 0.0M | 0.0M | 0.0M | 0.1% | 0.1% | 1.0bp | 0.0% | 1.0bp |
| SRR29685111 | 4.6M | 4.5M | 97.5% | 4.5M | 97.5% | 0.0M | 51.0bp | 49.8bp | 0.0M | 0.0M | 0.0M | 0.0M | 0.0M | 0.0M | 0.1% | 0.1% | 1.0bp | 0.0% | 1.0bp |
| SRR29685112 | 3.9M | 3.8M | 98.0% | 3.8M | 98.0% | 0.0M | 50.0bp | 49.7bp | 0.0M | 0.0M | 0.0M | 0.0M | 0.0M | 0.0M | 0.1% | 0.1% | 1.0bp | 0.0% | 1.0bp |

×

###### STAR: Summary Statistics: Columns

Uncheck the tick box to hide columns. Click and drag the handle on the left to change order. Table ID: `star_summary_table_table`

Show All
Show None

| Sort | Visible | Group | Column | Description | ID | Scale |
| --- | --- | --- | --- | --- | --- | --- |
| \|\| |  |  | Total reads | Number of input reads | `star-total_reads` | read\_count |
| \|\| |  |  | Aligned | Mapped reads | `star-mapped` | read\_count |
| \|\| |  |  | Aligned | % Mapped reads | `star-mapped_percent` |  |
| \|\| |  |  | Uniq aligned | Uniquely mapped reads | `star-uniquely_mapped` | read\_count |
| \|\| |  |  | Uniq aligned | % Uniquely mapped reads | `star-uniquely_mapped_percent` |  |
| \|\| |  |  | Multimapped | Multiple mapped reads | `star-multimapped` | read\_count |
| \|\| |  |  | Avg. read len | Average input read length | `star-avg_input_read_length` |  |
| \|\| |  |  | Avg. mapped len | Average mapped length | `star-avg_mapped_read_length` |  |
| \|\| |  |  | Splices | Number of splices: Total | `star-num_splices` | read\_count |
| \|\| |  |  | Annotated splices | Number of splices: Annotated (sjdb) | `star-num_annotated_splices` | read\_count |
| \|\| |  |  | GT/AG splices | Number of splices: GT/AG | `star-num_GTAG_splices` | read\_count |
| \|\| |  |  | GC/AG splices | Number of splices: GC/AG | `star-num_GCAG_splices` | read\_count |
| \|\| |  |  | AT/AC splices | Number of splices: AT/AC | `star-num_ATAC_splices` | read\_count |
| \|\| |  |  | Non-canonical splices | Number of splices: Non-canonical | `star-num_noncanonical_splices` | read\_count |
| \|\| |  |  | Mismatch rate | Mismatch rate per base | `star-mismatch_rate` |  |
| \|\| |  |  | Del rate | Deletion rate per base | `star-deletion_rate` |  |
| \|\| |  |  | Del len | Deletion average length | `star-deletion_length` |  |
| \|\| |  |  | Ins rate | Insertion rate per base | `star-insertion_rate` |  |
| \|\| |  |  | Ins len | Insertion average length | `star-insertion_length` |  |

Close

---

##### Alignment Scores

Percentages
Export Plot

Created with MultiQC

---

#### FastQ Screen

*Version:* 
 `0.16.0`

Screens a library of sequences in FastQ format against a set of sequence databases to see if the composition of the library matches with what you expect.*URL: http://www.bioinformatics.babraham.ac.uk/projects/fastq\_screen**DOI: 10.12688/f1000research.15931.2*

##### Mapped Reads

Percentages
Export Plot

Created with MultiQC

---

#### fastp

*Version:* 
 `1.0.1`

All-in-one FASTQ preprocessor (QC, adapters, trimming, filtering, splitting...).*URL: https://github.com/OpenGene/fastp**DOI: 10.1093/bioinformatics/bty560*

Fastp goes through fastq files in a folder and perform a series of quality control and filtering.
Quality control and reporting are displayed both before and after filtering, allowing for a clear
depiction of the consequences of the filtering process. Notably, the latter can be conducted on a
variety of parameters including quality scores, length, as well as the presence of adapters, polyG,
or polyX tailing.

##### Filtered Reads

Filtering statistics of sampled reads.

Percentages
Export Plot

Created with MultiQC

---

##### Sequence Quality

Average sequencing quality over each base of all reads.

Read 1: Before filtering
Read 1: After filtering

Export Plot

Created with MultiQC

---

##### GC Content

Average GC content over each base of all reads.

Read 1: Before filtering
Read 1: After filtering

Export Plot

Created with MultiQC

---

##### N content

Average N content over each base of all reads.

Read 1: Before filtering
Read 1: After filtering

Export Plot

Created with MultiQC

---

#### Software Versions

Software Versions lists versions of software tools extracted from file contents.

Copy table

| Software | Version |
| --- | --- |
| FastQ Screen | `0.16.0` |
| fastp | `1.0.1` |

**MultiQC v1.25.1** 
- Written by Phil Ewels,
available on GitHub.

This report uses Plotly,
jQuery,
jQuery UI,
Bootstrap and
FileSaver.js.

×

##### Plot Table Data

Select Column

Select Column

Please select two table columns.

Close

×

##### Regex Help

Toolbox search strings can behave as regular expressions (regexes). Click a button below to see an example of it in action. Try modifying them yourself in the text box.

`^` (start of string)
`$` (end of string)
`[]` (character choice)
`\d` (shorthand for `[0-9]`)
`\w` (shorthand for `[0-9a-zA-Z_]`)
`.` (any character)
`\.` (literal full stop)
`()` `|` (group / separator)
`*` (prev char 0 or more)
`+` (prev char 1 or more)
`?` (prev char 0 or 1)
`{}` (char num times)
`{,}` (count range)

```
samp_1
samp_1_edited
samp_2
samp_2_edited
samp_3
samp_3_edited
prepended_samp_1
tmp_samp_1_edited
tmpp_samp_1_edited
tmppp_samp_1_edited
#samp_1_edited.tmp
samp_11
samp_11111
```

See regex101.com for a more heavy duty testing suite.

Close
